# Switchgrass metabolomics reveals striking genotypic and developmental differences in specialized metabolic phenotypes

**DOI:** 10.1101/2020.06.01.127720

**Authors:** Xingxing Li, Saurav J. Sarma, Lloyd W. Sumner, A. Daniel Jones, Robert L. Last

**Affiliations:** Department of Biochemistry and Molecular Biology, Michigan State University, MI 48824; DOE Great Lakes Bioenergy Research Center, Michigan State University, MI 48824; Department of Plant Biology, Michigan State University, MI 48824; Department of Biochemistry, University of Missouri, Columbia, MO 65211; Bond Life Sciences Center, University of Missouri, Columbia, MO 65211; MU Metabolomics Center, University of Missouri, Columbia, MO 65211; Interdisciplinary Plant Group, University of Missouri, Columbia, MO 65211

**Keywords:** switchgrass, sustainability, biofuels, biomass, ecotype, metabolomics, NMR, specialized metabolite, diosgenin, saponin

## Abstract

Switchgrass (*Panicum virgatum* L.) is a bioenergy crop that grows productively on lands not suitable for food production, and is an excellent target for low-pesticide input biomass production. We hypothesize that resistance to insect pests and microbial pathogens is influenced by low molecular weight compounds known as specialized metabolites. We employed untargeted liquid chromatography-mass spectrometry (LC-MS), quantitative gas chromatography-mass spectrometry (GC-MS) and nuclear magnetic resonance (NMR) spectroscopy to identify differences in switchgrass ecotype metabolomes. This analysis revealed striking differences between upland and lowland switchgrass metabolomes as well as distinct developmental profiles. Terpenoid and polyphenol derived specialized metabolites were identified, including steroidal saponins, di- and sesqui-terpenoids and flavonoids. The saponins are especially abundant in switchgrass extracts and have diverse aglycone cores and sugar moieties. We report seven structurally distinct steroidal saponin classes with unique steroidal cores and glycosylated at one or two positions. Quantitative GC-MS revealed differences in total saponin concentrations in leaf blade, leaf sheath, stem, rhizome and root. The quantitative data also demonstrated that saponin concentrations is higher in roots of lowland than upland ecotype plants, suggesting ecotypic specific biosynthesis and/or biological functions. These results enable future testing of these specialized metabolites on biotic and abiotic stress tolerance and can inform development of low-input bioenergy crops.

**One sentence summary:** Integrated mass-spectrometry and nuclear magnetic resonance spectroscopy based metabolomics reveal that switchgrass accumulates structurally diverse terpenoids and phenolics, which vary in abundance and structure in a tissue- and ecotype-specific manner.

## Introduction

Development of environmentally sustainable and economical production of transportation fuels and industrial feedstocks using plant biomass is an important goal for the bioeconomy. Dedicated energy crops that are productive with low or no chemical fertilizers and pesticides on land that is unsuitable for food and fiber crops have received much attention ^1^. This requires development of plants with a suite of ‘ideal’ traits ^2^, including perennial life cycle, rapid growth under low soil fertility and water content as well as resilience to pests and pathogens.

Plants are master biochemists, producing a wide variety of general and specialized metabolites adapted to their ecological niches ^3^. The structurally diverse tissue- and clade-specific specialized metabolites play varied roles in how plants cope with biotic and abiotic stresses, both by reducing deleterious impacts and promoting beneficial interactions. For instance, glucosinolates produced by crucifers such as mustard, cabbage and horseradish, mediate interactions with insect herbivores ^4^, and flavonoids that induce the rhizobial lipochitooligosaccharides (‘Nod factors’) initiate the rhizobium-legume nitrogen fixation symbiosis ^5^. The root accumulating avenacin triterpene saponins, are well documented to protect oat (*Avena spp*.) from the fungal pathogen indued ‘take-all’ disease ^6–8^, contributing to oat productivity. Modifying plant specialized metabolism is an attractive target for bioengineering or trait breeding to create low-input bioenergy crops that can thrive on ‘marginal’ lands unsuitable for food and fiber crops.

While hundreds of thousands of specialized metabolites are estimated to be produced by plants ^3^, there are reasons why this number is almost certainly an underestimate. First, these metabolites are taxonomically restricted, often showing interspecies or even intraspecies variation ^3^; thus any sampled species, ecotype or cultivar will have a small subset of the overall plant kingdom’s phenotypic diversity. Second, specialized metabolites tend to be produced in a subset of cell- or tissue-types in any plant species analyzed; thus, cataloging the metabolic potential of even a single species requires extraction of multiple tissues over the plant’s development. Third, accumulation of these metabolites can be impacted by growth conditions and induced by abiotic or biotic stress ^9^. Finally, identification and structural characterization of newly discovered metabolites require specialized capabilities, typically a combination of mass spectrometry (MS) and nuclear magnetic resonance (NMR) spectroscopy analysis ^10^.

The North American native perennial switchgrass (*Panicum virgatum* L.) has the potential to be cultivated as a low-input bioenergy crop for growing on nonagricultural land ^11^. The two principal ecotypes of switchgrass are phenotypically distinct, with variation in flowering time, plant size, physiology and disease resistance. Upland ecotypes exhibit robust freezing tolerance but produce relatively low biomass yield in part due to early flowering ^12–15^. Plants of the lowland ecotype typically found in riparian areas produce large amounts of biomass and are more flooding- and heat-tolerant, pathogen-resistant and nutrient-use-efficient than the upland ecotype ^12,16,17^. However, these lowland ecotypes do not perform well in northern areas, largely due to cold intolerance.

Although morphological and physiological properties associated with the adaptive divergence of upland and lowland switchgrass have been intensively studied, the specialized metabolite diversity and their ecotypic differences remain underexplored, partially due to the technical challenges mentioned above. Lee et al. ^18^ detected large amounts of diosgenin-derived steroidal saponins from aerial tissues of four different switchgrass cultivars. Diosgenin is synthesized from cholesterol and cyclized and oxidized through several spontaneous steps and enzymatic reactions catalyzed by cytochrome P450 enzymes (CYP450s) ^19^. It is the backbone of spirostanol-type steroidal saponins that are important defensive compounds with documented antimicrobial and antiherbivory activities ^20–22^. These natural compounds also have pharmaceutical values. Diosgenin has been used as a major precursor for synthesizing steroidal drugs including hormonal contraceptives and corticosteroid anti-inflammatory agents ^20^. Beyond steroidal saponins, quercetin-derived flavonoids ^23^ and biotic/abiotic stress-elicited C_10_ – C_20_ terpenes ^24,25^ were also identified in switchgrass. A comprehensive metabolomics survey will be beneficial to understand the natural product diversity in this important bioenergy crop.

In this study we developed and deployed approaches to compare metabolomes of three upland and three lowland switchgrass cultivars by high-throughput untargeted LC-MS, targeted gas chromatography (GC)-MS and nuclear magnetic resonance spectroscopy (NMR). The two ecotypes were documented to have distinct metabolomes, especially in rhizome and root. We identified seven structurally distinct classes of diosgenin-derived steroidal saponins and cataloged a variety of flavonoid glycosides as well as di- and sesqui-terpenoids. Steroidal saponins were notably abundant, accounting for more than 30% total ion counts (averaging 5 μg/mg of dry tissue weight) in lowland root from the reproductive developmental stage. Furthermore, ecotype and/or tissue type specific accumulations were observed for the individual saponin classes as well as the total saponins. Our study provides a comprehensive analysis of the specialized metabolites produced by different switchgrass cultivars and sets the stage for developing dedicated bioenergy crops with varied plant and microbiome traits.

## Results

### Metabolome comparisons between tissue type, developmental stages and genotypes

LC-MS was used to develop an overview comparison of the metabolomes of upland and lowland ecotypes in different *tissue types and developmental stages*. Tissue extracts in 80% methanol were prepared from a sample panel (**Fig S1B**) containing three upland (Dacotah, Summer and Cave-in-Rock) and three lowland (Alamo, Kanlow and BoMaster) cultivars grown from seed in a controlled environment (**Materials and Methods**). Shoot, rhizome and root tissues were analyzed from plants at three developmental stages – vegetative, the transition between vegetative and reproductive, and early reproductive **(Fig S1A)** ^26^. In total, 4,668 distinct metabolite features were identified from the positive mode dataset; these are annotated as retention time/mass-to-charge ratio pairs and include multiple in-source fragments from single analytes. Downstream statistical analysis focused on the 2586 of 4,668 features with threshold maximum abundance ≥ 500 (**Table S1)**.

We employed two complementary approaches to annotate the features by metabolite class. First, Relative Mass Defect (RMD) filtering ^27^ was used to guide assignments of all metabolite signals to putative chemical classes **(Materials and Methods**). As a result, 42% and 15% of the 2586 features in the dataset were annotated as terpenoid glycosides and polyphenol derived metabolites, respectively (**Fig 1A and Table S1 column E and F**). We then searched the molecular ion mass-to-charge ratios (*m/z*) and associated fragment ion information in available online mass-spectral databases and found strong matches to 169 previously characterized metabolites (**Table S1 column KL – KT**).

**Figure 1.**
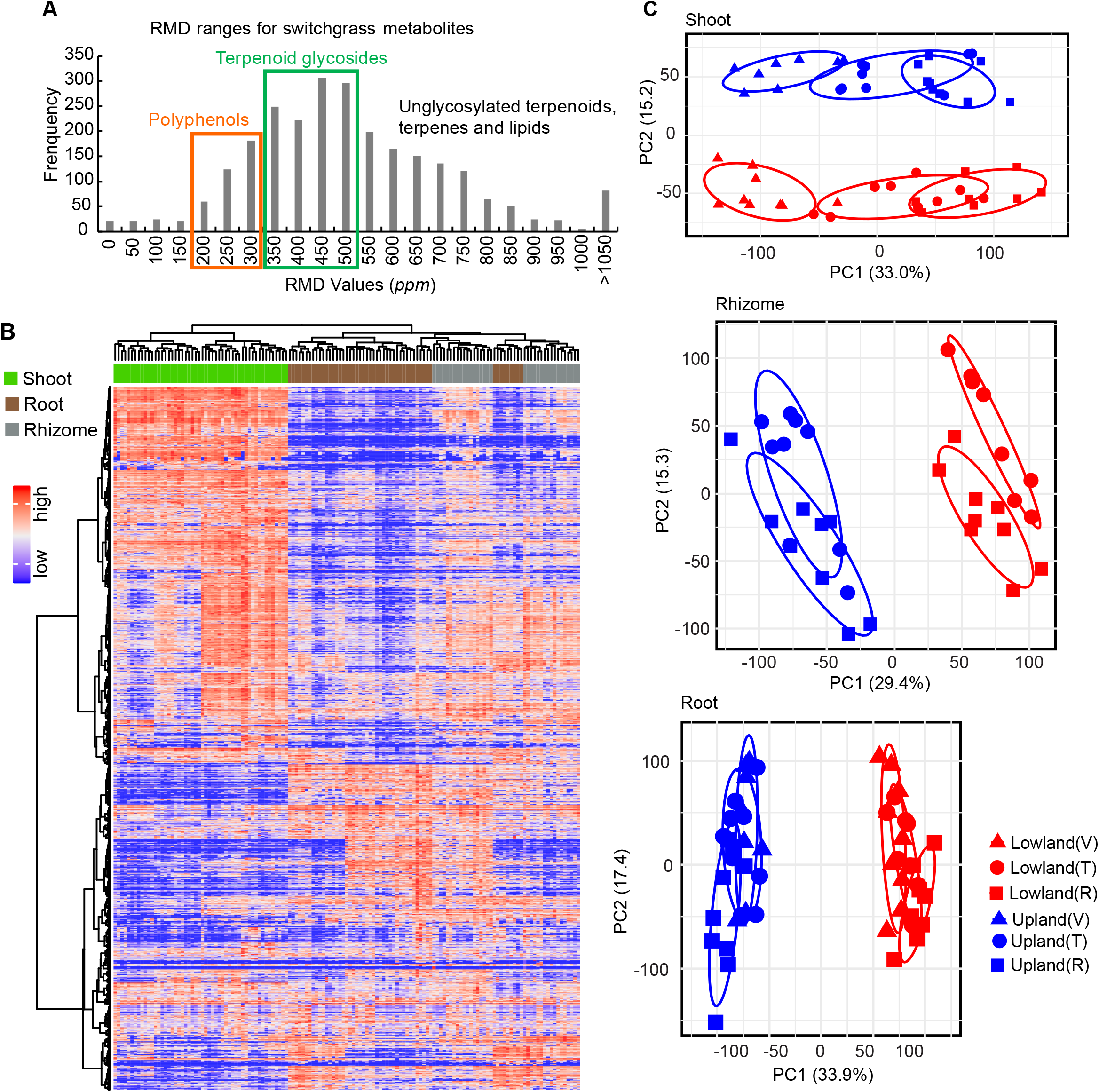
Untargeted metabolome profiling for switchgrass. (A) Histogram of RMD values for the total 2586 features detected in this study by LC-MS in positive ion mode. The green and orange rectangles highlight regions corresponding to the ranges of the RMD values anticipated for terpenoid glycosides and polyphenols, respectively. (B) The metabolome of the six switchgrass cultivars, three tissue types and three developmental stages shown by a heatmap with HCA. The row and column clusters symbolize the 2586 features and 48 sample groups (containing 139 individual samples), respectively. The values representing the metabolite abundances that were used to make the heatmap were scaled to a range from 0 (the lowest abundance) to 1 (the highest abundance). (C) PCA-score plots for the switchgrass shoot, rhizome and root metabolite profiles (n=8 for ‘Upland Vegetative’ and ‘Lowland Reproductive’; n=9 for all the other groups). The percentage of explained variation is shown on the x- and y-axes. V, vegetative phase; T, transition phase; R, reproductive phase.

The untargeted metabolome data provide a comprehensive view of specialized metabolite variation among the samples, and broad patterns of variation were revealed using hierarchical clustering analysis (HCA, **Fig 1B)**. The aerial (shoot) and subterranean tissue (root and rhizome) metabolites differed noticeably, consistent with the hypothesis that there are fundamental dissimilarities between the above- and below-ground tissue metabolomes. In contrast, the rhizome and root tissues were more similar to each other. Differences in metabolomes of each of the three tissue types across switchgrass cultivars (genotypes) were investigated using principal component analysis (PCA). The shoot metabolite profiles clustered into distinct groups in the PCA scores plot, corresponding with the upland (blue) and lowland (red) switchgrass ecotypes (**Fig 1C, top panel**). Separation of the metabolite profiles was especially clear for different developmental stages (developmental stages are differentiated by symbol shape and ‘V, T and R’, standing for ‘vegetative, transition and reproductive phase’ respectively in **Fig 1C, top panel**). The PCAs also showed clear-cut differences in metabolite profiles between the upland and lowland genotypes in both the rhizomes and roots (**Fig 1C, middle and bottom panels, respectively**). The developmental stage-associated variance in metabolite profiles of these two subterranean tissues was less apparent compared to that in the aerial tissue. Taken together, the metabolite PCA analysis revealed distinct patterns between upland and lowland switchgrass cultivars of the three tissues across the three developmental stages.

### Upland and lowland ecotypes have strikingly distinct metabolomes

We next focused on each *tissue type × developmental stage* combination and identified the metabolite features that differentially accumulated in either upland or lowland ecotypes. Surprisingly, 25% (256 of 1035, **Table 1**) of the features detected in extracts of the *vegetative-stage tillers* predominantly accumulated in one or the other switchgrass ecotype. Such features were termed as ecotype ‘differentially accumulated features’ (DAFs). Specifically, there are 126 upland enriched and 130 lowland enriched DAFs in extracts of the *vegetative-stage tillers* (**Fig 2A and Table 1**). Analysis of *vegetative-stage roots* further revealed a total of 879 features with 35% (310, **Table 1**) meeting the ecotype DAF statistical threshold. Of these 310 DAFs, similar numbers of features were found to be either upland- (149) or lowland- (161) enriched (**Fig 2B and Table 1**). DAFs were also identified for the other *developmental stage × tissue type* combinations (**Fig S2 and Table 1)**.

**Figure 2.**
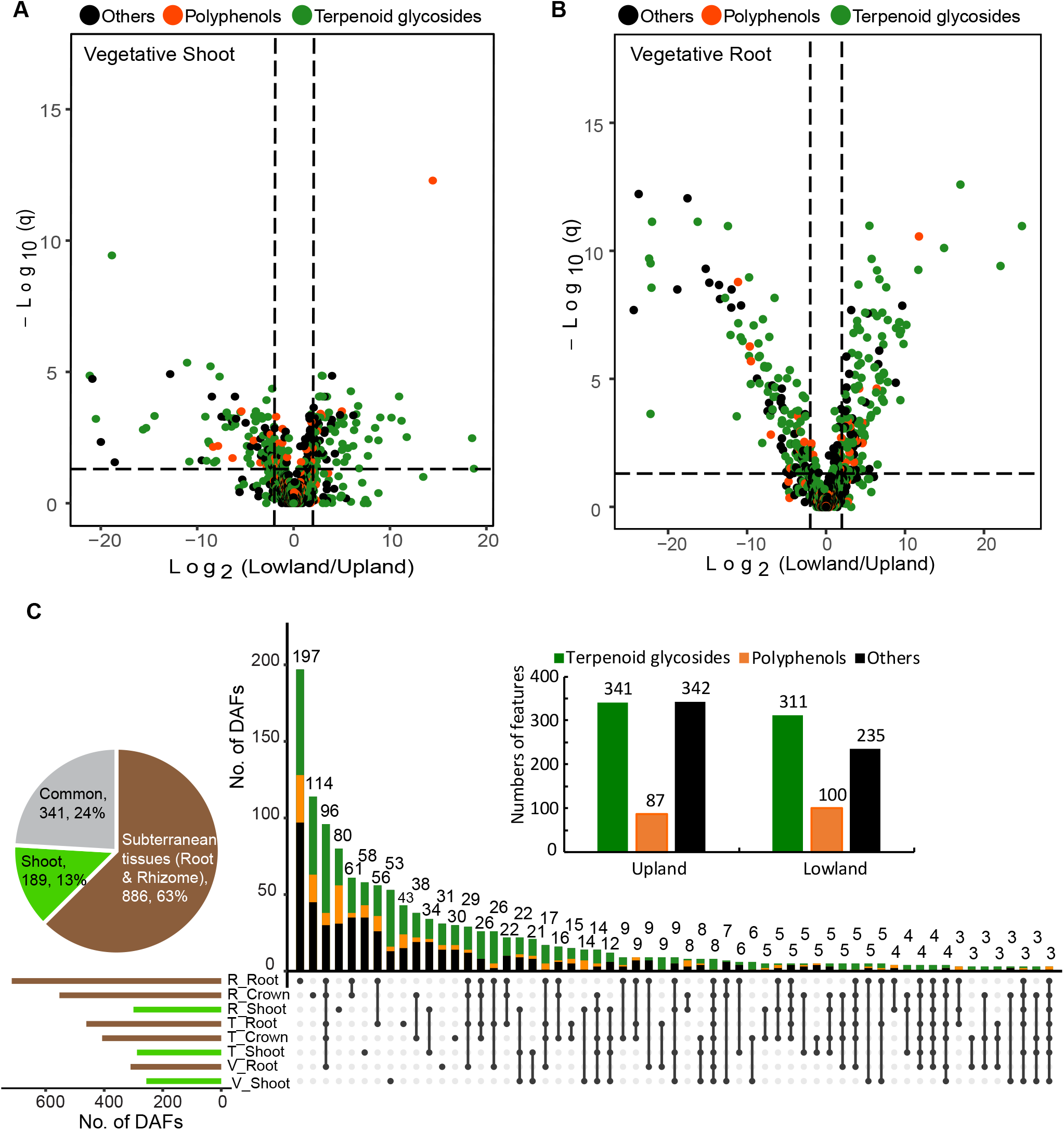
Differentially accumulated features (DAFs) were identified between the upland and lowland ecotypes. Significance analysis (cutoff threshold: FDR adjusted *p* ≤ 0.05; fold changes ≥ 2) was performed to screen for the DAFs between the upland and lowland switchgrass ecotypes (n = 8 or 9) in various *developmental stage × tissue type* samples. Results of the analyses for (A) *vegetative-stage shoots* and (B) *vegetative-stage roots* are shown here using volcano plots. Putative terpenoid glycosides, polyphenols and metabolites from the other categories were classified using RMD filtering and the results color coded. (C) In total, 1416 unique (non-overlapping) ecotype DAFs were identified for the eight *developmental stage × tissue type* combinations. The inserted barplot shows that upland and lowland ecotypes accumulated similar numbers of the predominant DAFs likely terpenoid glycosides (green) and polyphenols (orange). The inserted pie chart indicates percentages of the DAFs contributed by aerial (shoot) vs. subterranean (root/rhizome) tissues. V, vegetative phase; T, transition phase; R, reproductive phase.

**Table 1.** Numbers of the total detected features and differentially accumulated features (DAFs) in each of the eight *developmental stage × tissue type* sample classes.

| *Developmental stage x Tissue type* | No. of total detected features <sup>a</sup> | No. of DAFs<sup>b</sup> |  |  |
| --- | --- | --- | --- | --- |
|  |  | Upland enriched | Lowland enriched | Total |
| *Vegetative-stage x shoot* | 1035 | 126 | 130 | 256 |
| *Vegetative-stage x root* | 879 | 149 | 161 | 310 |
| *Transition-stage x shoot* | 1037 | 155 | 133 | 288 |
| *Transition-stage x rhizome* | 1114 | 268 | 139 | 407 |
| *Transition-stage x root* | 965 | 226 | 235 | 461 |
| *Reproductive-stage x shoot* | 1612 | 100 | 200 | 300 |
| *Reproductive-stage x rhizome* | 1378 | 324 | 229 | 553 |
| *Reproductive-stage x root* | 1327 | 408 | 309 | 717 |

<sup>a</sup> Sum of the numbers in this column is larger than the numbers of the total detected features (2586) as some features were counted multiple times in several sample classes.
<sup>b</sup> Sum of the numbers in the column of Total is larger than the total detected unique DAFs (1416) as some DAFs were counted multiple times. The detailed information regarding these DAFs can be found in **Table S2 – 9**. Criteria for the DAFs: FDR adjusted $p \leq 0.05$ ; fold changes $\geq 2$ .

Altogether, 1416 unique ecotype DAFs were identified for the eight *tissue type × developmental stage* combinations included in this study (**Fig 2C and Table S2 – 9)**, accounting for approximately half of the features in the full dataset. Based on RMD filtering, 46% and 13% of the DAFs were predicted to be terpenoid glycosides and polyphenol-derived metabolites, respectively (**green and orange dots on Fig 2 A-B and Fig S2)**. The numbers of the DAFs preferentiality accumulated in upland and lowland ecotypes are equivalent to each other (**Fig 2C inset barplot**). Furthermore, 63% of the DAFs were found in subterranean tissues while only 13% were unique to the aerial tissues. The remaining 24% were detected in both above- and below-ground tissues (**Fig 2C inset pie chart)**. These results reveal that switchgrass subterranean tissues are the major sources of the observed specialized metabolic ecotypic diversity.

### Detailed analysis of metabolite diversity using MS/MS

The observation that terpenoid and polyphenol metabolite classes are highly represented in DAFs led us to use high resolution LC-MS/MS to characterize these features in more detail. In total, we annotated 72 saponins, 10 diterpenoids, four sesquiterpenoids and seven flavonoid glycosides (**Fig 3A and Table S11 – 14)** from the six switchgrass cultivars. The identified saponins (**Table S14**) vary in their precursor masses and RTs due to differences in both aglycones and sugar moieties. Assignments of multiple fragment ions generated by collision induced dissociation (CID) resulting from the losses of sugar units allowed for annotation of the conjugated monosaccharides as well as the aglycones. For 44 (out of the 72) saponins, the presence of signal at *m/z* 415 was diagnostic of the diosgenin aglycone ^28^. Fragmentations of 20 saponins yielded an aglycone fragment ion at *m/z* 431, while further loss of 18 Da (H_2_O) resulted in *m/z* 413; this suggests that this core has an additional oxygen atom compared to diosgenin. For three saponins, the presence of signal at *m/z* 417 was indicative of a tigogenin aglycone ^29^. Finally, larger aglycone fragments at *m/z* 457 and 473 were observed for six and nine saponins, respectively. Elemental composition suggested that these two aglycones both have 29 carbons and differed by one oxygen, as predicted for nortriterpenoids. Moreover, the LC-MS spectra for about two thirds of the 72 saponins displayed an abundant [M+H-H_2_O]^+^ ion, characteristic of furostanol saponins, which contain a C-22 labile hydroxyl group due to the hemiketal structure. The furostanol saponins are often glycosylated at their sidechains (also shown by our NMR results below, **Fig 4 B – H**). Therefore, based on type of aglycone and sidechain glycosylation, we grouped the 72 identified saponins into seven classes: D-415-SCG, D-415, D-431-SCG, D-431, D-417-SCG, D-457 and D-473 (‘D’ indicates a diosgenin derived aglycone; numerical number reflects *m/z* value of aglycone fragment ion detected by positive mode MS; ‘SCG’ indicates ‘sidechain glycosylation’).

**Figure 3.**
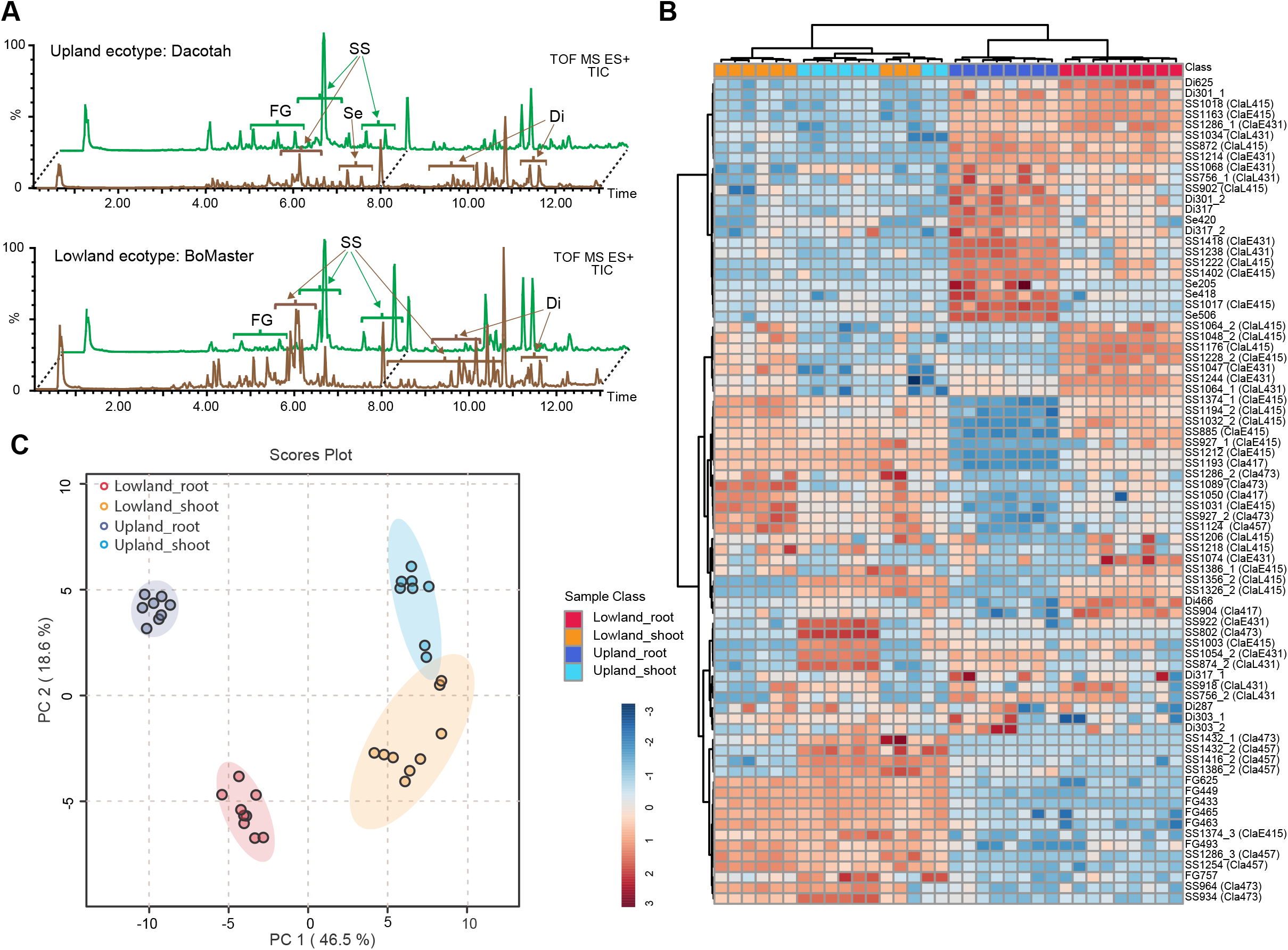
Upland and lowland ecotypes are distinct in specialized metabolite-profile. (A) Total ion chromatograms (TIC) for upland Dacotah and lowland BoMaster shoot (green) and root (brown) extracts. Areas where the specialized metabolites were identified are labeled by the short names. FG, flavonoid glycoside; ESS, early eluting steroidal saponin; LSS, late eluting steroidal saponin; TS, triterpenoid saponin; Di, diterpenoid; Se, sesquiterpenoid. (B) Heatmap showing relative abundances of the specialized metabolites (vertical axis) across the biological samples (horizontal axis). Classifications of the saponins were shown in the parentheses. Relative metabolite abundances were log10 scaled to a range between −3 (lowest) and 3 (highest). Clustering method/distance: Ward/Eucidean. (C) PCA scores plot showing distinct separations of the specialized metabolite-profiles among upland shoot, upland root, lowland shoot and lowland root.

**Figure 4.**
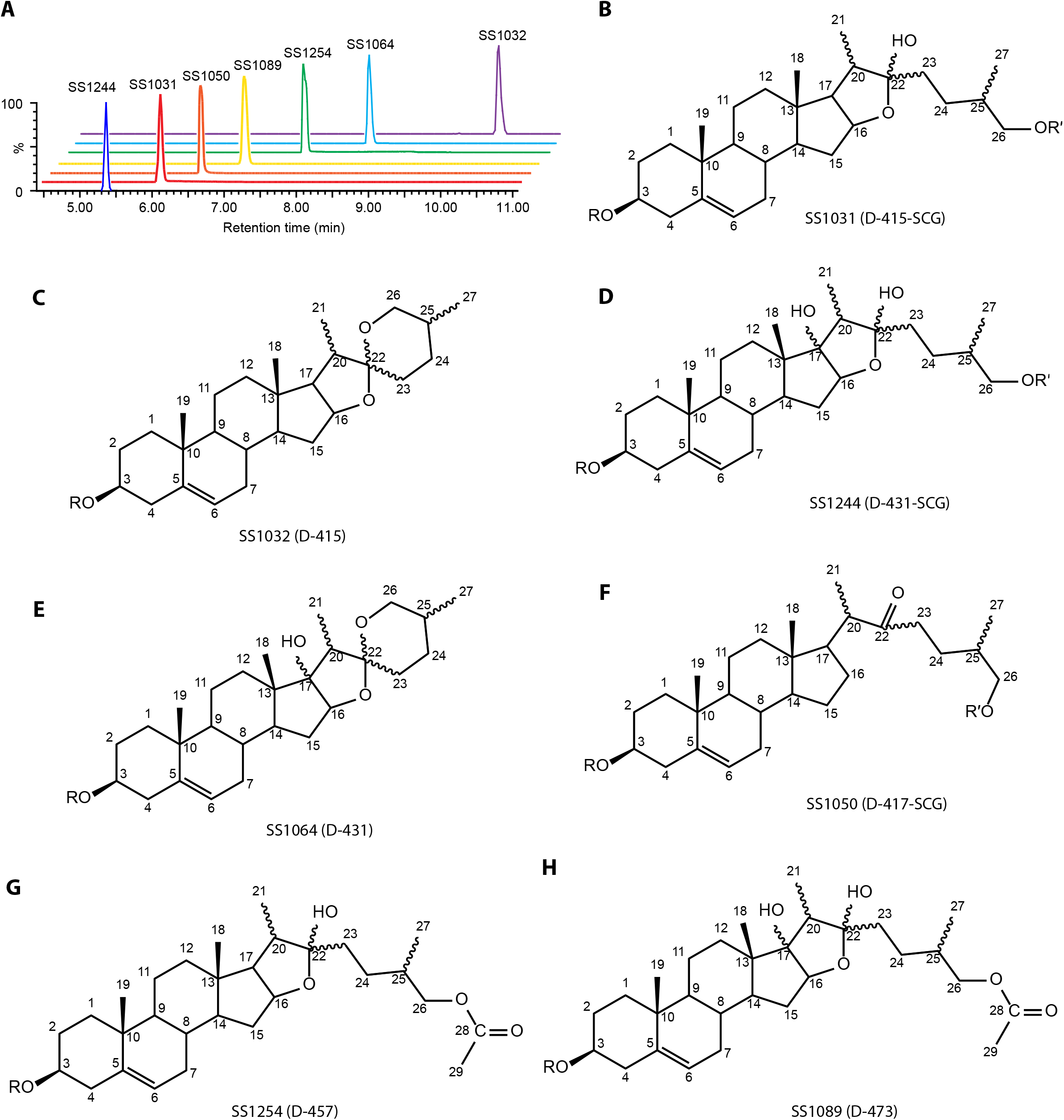
Chemical structures of the saponins identified in switchgrass. (A) The seven purified saponins, representing the seven switchgrass saponin classes, have distinct RTs between 5 and 11 min. (B) – (H) Structures and numbering of the aglycones for the saponins SS1031, SS1032, SS1244, SS1064, SS1050, SS1254 and SS1089. Classifications of the saponins were shown in the parentheses. R and R′ indicate position of sugar moiety at C-3 and C-26 (on the side chain), respectively.

PCA analysis (**Fig 3C**) using all the identified specialized metabolite features as the loadings (**Fig 3B and Fig S5**) revealed that ecotype plays an important role in the separation of the metabolite profiles even when multiple tissue types are included. Specifically, flavonoid glycosides predominately accumulated in shoot while diterpenoids and sesquiterpenoids preferentially accumulated in root (**Fig 3B and Fig S5A**); together these features separated the tissue types on the PCA (**Fig 3C**). Notably, all four sesquiterpenoids (**Table S12**) were exclusively detected in upland ecotype root, >1000-fold higher than they were found in lowland root (**Table S10**). In contrast, a diterpenoid glycoside, Di466 (**Table S13**), was abundant in lowland root but nearly undetectable in upland root samples (also a >1000-fold accumulation difference, **Table S10**). Hence, such features also contribute to separation of the ecotypes on the PCA (**Fig 3C**).

Accumulation patterns of the saponins **(Fig 3B and Fig S5B**) were relatively complex compared with the C15 and C20 terpenes. Five general patterns were observed: 1) D-415 saponins preferentially accumulated in lowland ecotype root; 2) D-431-SCG and D-431 saponins preferentially accumulated in root tissues but accumulation was not ecotype specific; 3) D-457 and D-473 saponins predominately accumulated in shoot with no apparent ecotype specificity; 4) D-417-SCG saponins preferentially were found in lowland shoot; 5) D-415-SCG saponins showed neither strong tissue nor ecotype specific accumulating.

### NMR characterization of saponins

To unequivocally determine the structures for the switchgrass saponins we selected seven saponins that represent unique classes (**Fig 4A**) for HPLC-purification: SS1031, SS1032, SS1244, SS1064, SS1050, SS1254 and SS1089 (‘SS’ stands for steroidal saponin). Their molecular formulas were proposed based on positive mode high resolution LC-MS/MS analysis (**Fig S6 – 12**). NMR spectra (^1^H, DEPTQ, HSQC, COSY, HMBC and TOCSY) were generated for them (**Table S15 – 21 and Fig S13 – 61**). The proton resonances determined from ^1^H, COSY and TOCSY spectra in all samples fell into three distinct chemical shift regions: 0.8 – 2.6 *ppm* from aglycone backbone hydrogens; 3.2 – 4.2 *ppm* from sugar hydrogens; as well as sugar ring anomeric and aglycone olefinic hydrogens from 4.2 – 5.6 *ppm*. The HSQC, HMBC and DEPTQ spectra were used to assign chemical shifts of the saponin core carbons, beginning with those downfield shifted carbons, due to direct connections with oxygens or double bonds. When combined with ^1^H – ^1^H couplings established from COSY and TOCSY spectra, we could assign each aglycone position. Positions of sugar moiety substitutions were determined from HMBC spectra, based on ^1^H – ^13^C correlations separated by two or three bonds. All seven saponins were glycosylated at the C-3 hydroxyl group, while three were also glycosylated at the C-26 position (side-chain glycosylation). We also observed an olefinic hydrogen from HSQC and ^1^H, indicating a double bond between the C-5 and C-6, for all seven saponins; thus, it rules out the possibility that SS1050 contained a tigogenin core. Collectively these data identified SS1031 (**Fig 4B**) as the previously characterized switchgrass saponin, protodioscin, with sidechain glycosylation ^18^ and SS1032 (**Fig 4C**) as a steroidal saponin derived from a diosgenin with the characteristic core spiroketal moiety but no sidechain glycosylation. These two saponins are likely biosynthetically related as the final six-member heterocyclic ring closure forming the spiroketal structure in diosgenin is spontaneous *in planta* ^19^. Similarly, SS1244 (**Fig 4D**) and SS1064 (**Fig 4E**) are related saponins that are derived from diosgenin: the former with side chain glycosylation and the latter without. Both share an additional tertiary hydroxyl group on the C-17. The relatively low-abundance SS1050 (**Fig 4F**) has a cholesterol-like C_27_ aglycone with only four rings, side chain glycosylation and a C-22 ketone group (characterized by a chemical shift of 215.6 *ppm* for the carbonyl carbon).

A unique feature of the diosgenin-derived saponins SS1254 (**Fig 4G**) and SS1089 (**Fig 4H**) is that they appear to be acetylated -- rather than glycosylated -- at C-26. This is consistent with the observed C_29_ aglycone mass of 456 Da (42 Da larger than that of diosgenin, which is 414 Da) and 472 Da (42 Da larger than the 430 Da of oxydiosgenin) respectively. This structural feature was revealed by HMBC data where the C-28 carbonyl carbon at 171.0 *ppm* is only correlated with the C-26 methylene and C-29 methyl protons. SS1089 has an additional tertiary hydroxyl group on the C-17 position compared to SS1254.

### Differential saponin accumulation between upland and lowland revealed by LC- and GC-MS analysis

As an approach for quantifying differential accumulation of the saponins that is orthogonal to PCA loading plots (**Fig S5B**) and HCA (**Fig 3B**), we compared cumulative ion counts for each saponin class between the ecotypes across different *tissue × developmental stage* sample groups. The D-415-SCG and D-415 were the dominant saponin forms in terms of their accumulative ion counts and detectable, and were approximately one order of magnitude higher than the other classes. The lowland ecotypes accumulated much higher levels of these saponins than did upland in root tissues, and this was especially notable during the reproductive stage (**Fig 5 A and B**). The two C-17 hydroxylated saponin classes, D-431-SCG and D-431, had higher accumulations in below-ground tissues, without apparent accumulation differences between the two ecotypes (**Fig 5 C and D**). In contrast, the two acetylated sidechain saponin forms (D-457 and D-473) were found to have much higher accumulations in above-ground tissue than below-ground tissues (**Fig 5 F and G**). The only saponin form not oxidized at the C-16 position (D-417-SCG, **Fig 5E**) showed a tendency towards higher accumulations in lowland shoot at later developmental stage. For the total (summed) saponins, lowland root accumulated more than upland root at all three developmental stages. In contrast, shoots of the two ecotypes accumulated comparable amounts of total saponins across development stages (**Fig 5H upper panel**). The saponins represented about 20 to 35% vs. 5 to 10% of the total ion counts in lowland and upland root, respectively (**Fig 5H lower panel**), indicating the contribution of these specialized metabolites in defining distinct phenotypes between ecotypes.

**Figure 5.**
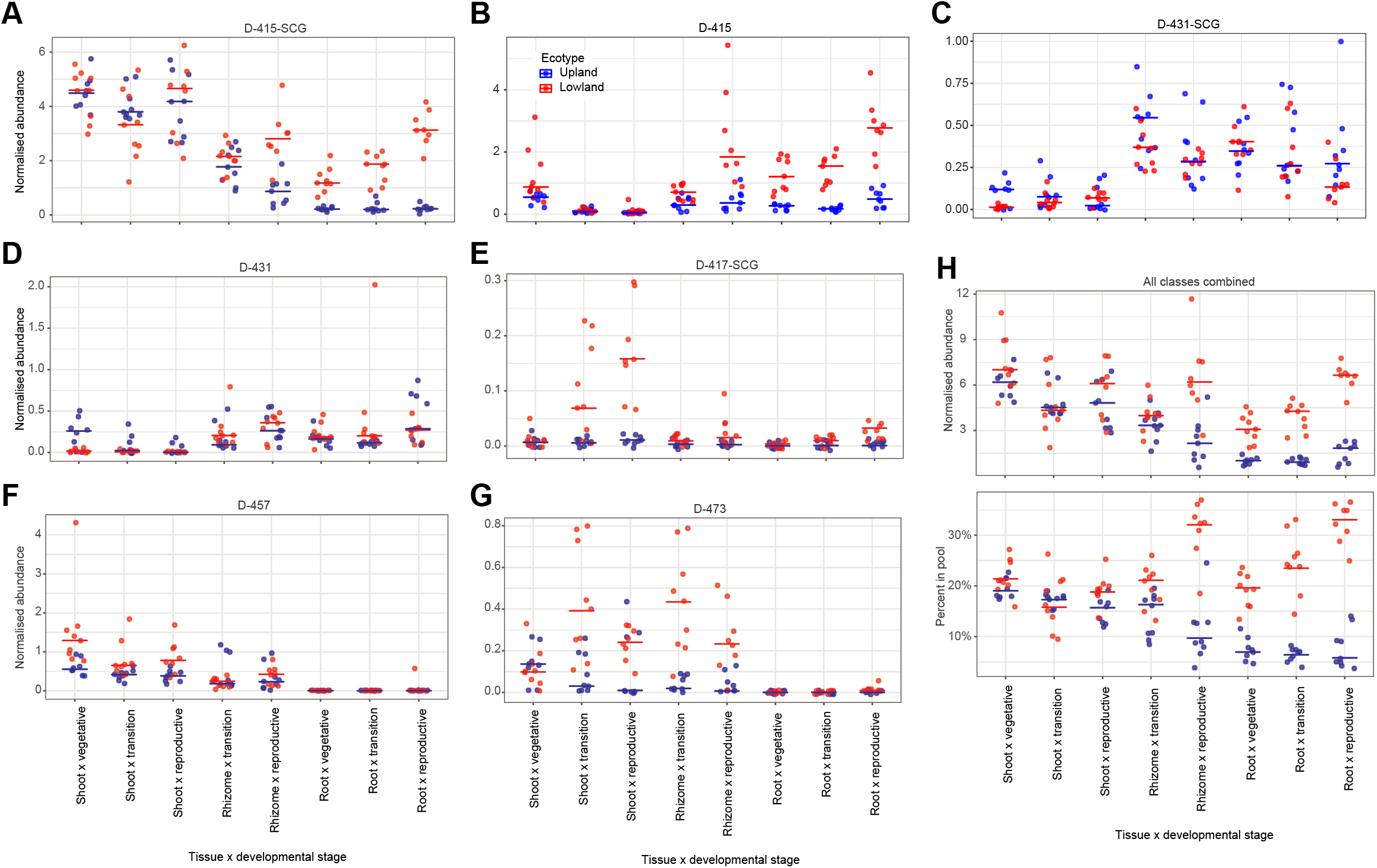
Relative quantification of individual saponin classes and total saponins. (A) – (G) Accumulative ion counts for each individual saponin classes and (H, lower panel) the total saponins (all classes combined) from the eight *tissue type × developmental stage* sample classes measured by positive mode LC-MS. (H, lower panel) Percentages of the saponins in the total ion pools. Normalized abundances (y-axis) were calculated as (*ion intensity of the feature / ion intensity of the internal standard*). For all panels, the horizontal bars represent median values (n = 8 or 9). Red, lowland ecotype; Blue, upland ecotype.

While this LC-MS approach is excellent for documenting the diversity of saponin types, it is not ideal for quantification because of the uneven ionization efficiencies of the early- vs. late-eluting analytes (caused by the changing ratio of water and organic mobile-phase during chromatography). To more accurately determine total saponin concentrations in different tissues and cultivars, a GC-MS based quantification method was developed to quantify the sapogenins after hydrolytic removal of sugars through the comparison with an authentic diosgenin standard (**Materials and Methods**). As a result, we identified six diosgenin-derived steroidal sapogenin peaks and two peaks annotated as triterpenes (**Fig 6A and Table 2**). The triterpene peaks were only present in leaf *blade* samples and might arise from acetylated steroids if the hydrolytic removal of acetates was incomplete.

**Figure 6.**
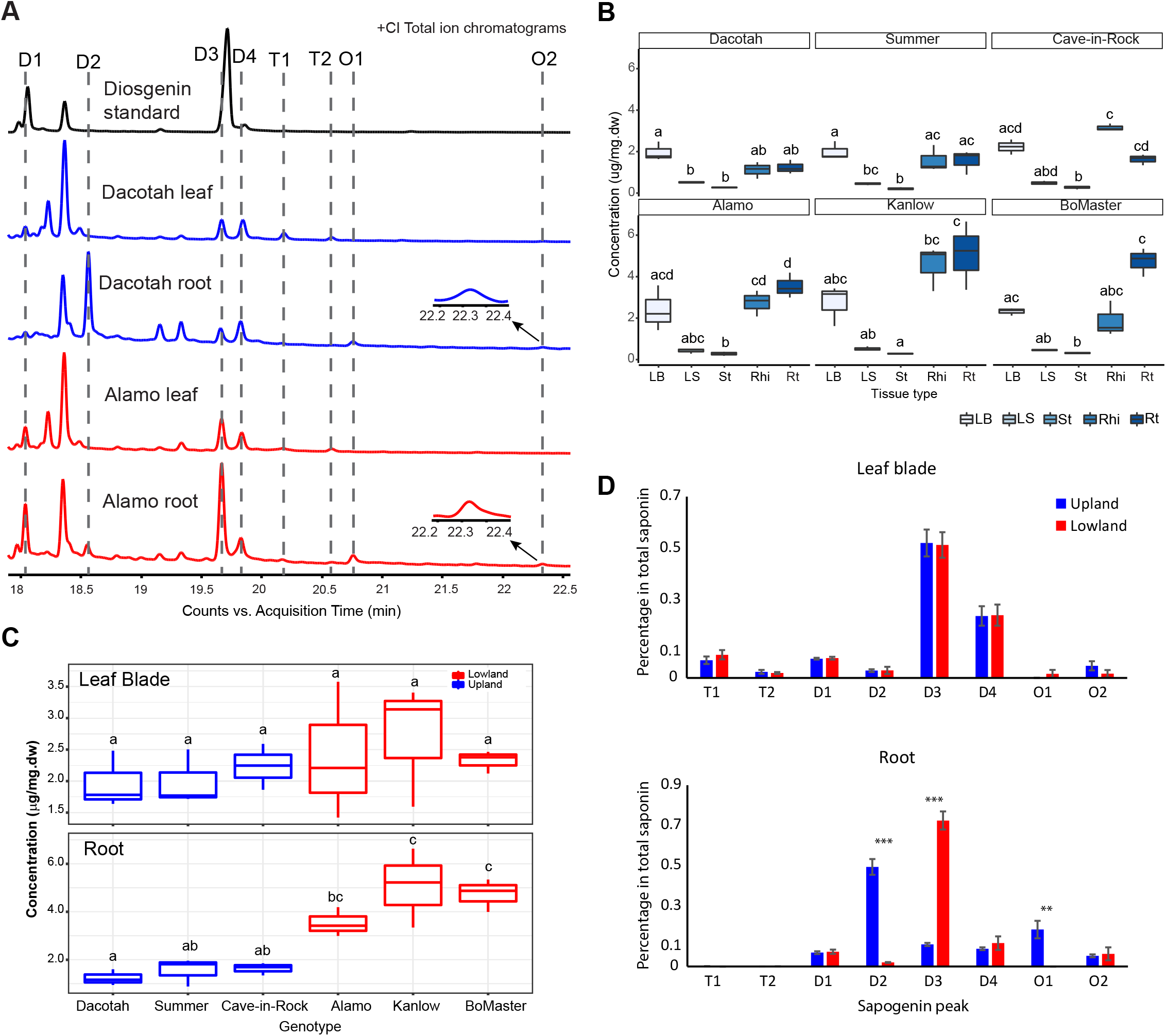
Sapogenin aglycone peaks identified in switchgrass extracts were quantified by GC-MS. (A) +(CI) GC-MS total ion chromatograms (TICs) of diosgenin standard (black), Dacotah leaf and root (blue), Alamo leaf root (red). The identified sapogenin aglycone peaks in the switchgrass and standard samples are indicated and aligned by the dashed lines. Zoomed-in views for the peak O2 in Dacotah root and Alamo root are indicated by the arrows. (B) Comparison of the total sapogenin concentrations among the five tissue types for each switchgrass cultivar (Kruskal-Wallis test: *p* = 0.012, 0.026, 0.009, 0.024, 0.017 and 0.011 for Dacotach, Summer, Cave-in-Rock, Alamo, Kanlow and BoMaster respectively). LB, leaf blade; LS, leaf sheath, St, stem; Rhi, rhizome; Rt, root. (C) Comparison of the total sapogenin among the six switchgrass cultivars in leaf blade (Kruskal-Wallis test: *p* = 0.766) and root (Kruskal-Wallis test, *p* = 0.016). Different lower-case letters on top of the boxes designate statistically different means (Post-Hoc test: Dunn’s test). (D) Ratio of the individual sapogenins in leaf blade (upper) and root (lower) of upland and lowland ecotypes. Heights of the bars reflect the means of the nine replicates (three cultivars × three replicates) for each ecotype; error bars show the standard error of the mean; ** standards for 0.001 ≤ *p* ≤ 0.01, *** standards for *p* < 0.001 (one tailed t-test).

**Table 2.** TMS derivatized sapogenin aglycones identified in switchgrass extracts by chemical ionization (CI) GC-MS analysis. The TMS-derivatized molecules are considered as the molecular ions in each case here. TMS, trimethylsilyl group [Si(CH_3_)_3_].

| Peak | Retention index (RI)<sup>a</sup> | Key ion annotations (*m/z*)<sup>b</sup> | Abundant fragment ion *m/z*<sup>c</sup> | Formula of the neutral molecule | Mass of the neutral molecule (Da) | Annotation of the neutral molecule<sup>d</sup> |
| --- | --- | --- | --- | --- | --- | --- |
| D1 | 3114 | 397 [M+H]<sup>+</sup> | 139, 282 | C<sub>27</sub>H<sub>40</sub>O<sub>2</sub> | 396 | Anhydro-diosgenin |
| D2 | 3194 | 397 [M+H-TMS-OH]<sup>+</sup>, 471 [M-CH<sub>3</sub>]<sup>+</sup>, 485 [M-H]<sup>+</sup>, 487 [M+H]<sup>+</sup> | 139, 282 | C<sub>30</sub>H<sub>50</sub>O<sub>3</sub>Si | 486 | Diosgenin (TMS derivatized) |
| D3 | 3339 | 397 [M+H-TMS-OH]<sup>+</sup>, 471 [M-CH<sub>3</sub>]<sup>+</sup>, 485 [M-H]<sup>+</sup>, 487 [M+H]<sup>+</sup> | 139, 187, 282 | C<sub>30</sub>H<sub>50</sub>O<sub>3</sub>Si | 486 | Diosgenin (TMS derivatized) |
| D4 | 3361 | 397 [M+H-TMS-OH]<sup>+</sup>, 471 [M-CH<sub>3</sub>]<sup>+</sup>, 485 [M-H]<sup>+</sup>, 487 [M+H]<sup>+</sup> | 139, 187, 282 | C<sub>30</sub>H<sub>50</sub>O<sub>3</sub>Si | 486 | Diosgenin (TMS derivatized) |
| T1 | 3399 | 409 [M+H-TMS-OH]<sup>+</sup>, 483 [M-CH<sub>3</sub>]<sup>+</sup>, 497 [M-H]<sup>+</sup> | 218, 203, 190 | C<sub>33</sub>H<sub>58</sub>OSi | 498 | β-Amyrin (TMS derivatized) |
| T2 | 3439 | 409 [M+H-TMS-OH]<sup>+</sup>, 483 [M-CH<sub>3</sub>]<sup>+</sup>, 497 [M-H]<sup>+</sup> | 218, 203, 190 | C<sub>33</sub>H<sub>58</sub>OSi | 498 | β-Amyrin (TMS derivatized) |
| O1 | 3462 | 395 [M+H-TMS-OH-H<sub>2</sub>O]<sup>+</sup>, 413[M+H-TMS-H<sub>2</sub>O]<sup>+</sup>, 487 [M-CH<sub>3</sub>]<sup>+</sup>, 501 [M-H]<sup>+</sup>, 503 [M+H]<sup>+</sup> | 139, 187, 298 | C<sub>30</sub>H<sub>50</sub>O<sub>4</sub>Si | 502 | Oxydiosgenin (TMS derivatized) |
| O2 | 3620 | 395 [M+H-TMS-2H<sub>2</sub>O]<sup>+</sup>, 413[M+H-TMS-OH]<sup>+</sup>, 487 [M-CH<sub>3</sub>]<sup>+</sup>, 501 [M-H]<sup>+</sup>, 503 [M+H]<sup>+</sup> | 139, 187, 298 | C<sub>30</sub>H<sub>50</sub>O<sub>4</sub>Si | 502 | Oxydiosgenin (TMS derivatized) |

<sup>a</sup> The RIs were calculated using a homologous series of n-alkane standards with the same GC-MS method as the samples (**Fig. S4A**).
<sup>b</sup> The *m/z* information for the key ion annotations was obtained by (CI) GC-MS (**Fig. S62**).
<sup>c</sup> The *m/z* information for the fragment ions was obtained by (EI) GC-MS (**Fig. S4B**).
<sup>d</sup> The annotations were made based on the pseudomolecular ions and fragment ion information from electron ionization (EI) MS (**Fig S4B**) by comparing with the diosgenin standard and/or searching against National Institute of Standards and Technology (NIST) NIST17 GC-MS library (www.chemdata.nist.gov) for the best matching compounds.

The total saponin (all eight peaks summed) concentrations were first compared across tissue types for each switchgrass cultivar (**Fig 6B and Table S22**). For every cultivar, saponins were detectable in all five analyzed tissue types and higher in leaf *blade*, rhizome and root than they were in leaf *sheath* and shaved stem. The two upland cultivars Dacotah and Summer have the highest saponin concentration in leaf blade, whereas all three lowland cultivars accumulate the highest saponin level in root. The third upland cultivar, Cave-in-Rock, however, accumulates the highest total saponin in rhizome. We then compared the total saponins in leaf *blade* and root across the six cultivars. The total saponin concentrations in lowland roots are uniformly higher than those in roots of the three upland cultivars, with Kanlow and BoMaster showing statistical significance (*p* < 0.05, Kruskal-Wallis test). In comparison, leaf blade total saponins revealed no ecotype-related statistical difference (**Fig 6C**). Moreover, different sapogenin compositions were also found between upland and lowland roots but not leaf blades (**Fig 6D**). This was due to the differentiated accumulations of two diosgenin isomers, D2 and D3, and one oxydiosgenin isomer, O1 identified only in root. Taken together, quantitative analysis of sugar-free sapogenins supports the results of LC-MS analysis showing strong genetic difference in root saponin accumulation. Considering the high abundances of the saponins, water solubilities of these molecules and the documented bioactivities to the microbes ^30^, the differential accumulation in root might play a role in shaping the ecotype specific rhizosphere microbiomes in switchgrass.

## Discussion

Plants produce a plethora of structurally diverse specialized metabolites that serve roles ranging from attracting beneficial organisms to combating deleterious biotic and abiotic agents. However, domestication and improvement of crops typically leads to reduced amounts and types of these advantageous metabolites. While restoring these beneficial traits to existing crops is generally not feasible, development of new food, fuel and fiber crops can be done in such a way to maintain or enhance existing metabolic variation, potentially limiting the need for toxic pesticides.

Switchgrass is a compelling example of a low-fertilizer and low-pesticide cellulosic bioenergy crop in the USA ^11^. Switchgrass populations (ecotypes) are found from the northern Midwest to Texas, and along the east coast; unlike the relatively narrow genetic and phenotypic variation of traditional crops, there is abundant genetic variation across these populations. Striking differences in numerous plant traits were documented between populations of the upland and lowland ecotypes ^31^, including biomass production ^32^, rust pathogen (*Uromyces graminicola*) resistance ^33^ as well as tolerance to low nitrogen, drought and freezing conditions ^14,34,35^. These documented phenotypic and physiological differences encouraged us to compare the upland and lowland switchgrass metabolomes and catalog the ecotype specific specialized metabolites. In addition to being of interest for pathway discovery, and as tools to understand the genetic architecture of switchgrass, such information should be valuable for breeding switchgrass varieties that highly productive with no or low pesticides inputs.

Analysis of six switchgrass accessions representing both upland and lowland ecotypes of eight specific *tissue type × developmental stage* sample classes identified a remarkable amount of metabolite variation. In fact, 1416 ecotype DAFs (**Table 1 and Table S2 – 9**) account for half of the metabolite features detected in this study. Many of these differences were quite large: there were 157 DAFs showing >1000-fold accumulation difference between the two ecotypes in at least one of the eight sample classes (**Table S10)**. In contrast, published metabolomics analysis for maize showed that metabolite profiles of the six genetically defined GWAS populations failed to separate in PCA, even when the metabolite data were independently analyzed within the same tissue type ^36^.

Based on RMD filtering ^27^, 46% and 13% of the switchgrass ecotype specific DAFs are proposed to be terpenoid glycosides and polyphenol derived metabolites respectively. Relatively few switchgrass terpenoid and polyphenol specialized metabolites were identified in the past, including leaf steroidal saponins ^18,37^, diterpenoid derived antimicrobial phytoalexins ^38^ and quercetin based flavonoids ^23^. Our identification of more than 1,000 unannotated DAFs suggests the possibility of many more switchgrass terpenoids and polyphenols to be characterized. By generating MS/MS spectra beginning with the most abundant metabolites, we identified close to 100 specialized metabolites. In contrast to the flavonoid glycosides (**Table S11**), the diterpenoids (**Table S12**), sesquiterpenoids (**Table S13**) and most of the steroidal saponins (**Table S14**) did not match known compounds in MS/MS databases (**Material and Methods**): this led us to subject representative saponins to NMR analysis.

Saponins stand out in this study as highly abundant and differentially accumulating metabolites, exhibiting diversity in their cores, as well as glycosylation positions and types. This structural diversity indicates tissue and/or genotype specific activities of as yet uncharacterized CYP450s, UDP-glycosyltransferases (UGTs) and other tailoring enzymes (e.g., acyltransferases). This conclusion is supported by the seven distinct sapogenin aglycone structures elucidated by LC-MS and NMR results and their accumulation patterns in switchgrass. These include diosgenin cores with the characteristic 5,6-spiroketal moiety (**Fig 4C**), as well as a diosgenin core that presumably is derived from a metabolic intermediate ^19^, in which the sidechain is stabilized by glycosylation and thus prevented from the spontaneous cyclization to form the final pyranosidic ring **(Fig 4B**); this metabolite was previously identified in switchgrass aerial tissues ^18^. Besides glycosylation, the sidechain also can be stabilized by acetylation resulting in the C_29_ aglycones (**Fig 4 G and H**). Predominant accumulation of the sidechain acetylated saponins in shoot implies involvement of one or more tissue specific acyltransferase activities in switchgrass saponin biosynthesis. Likewise, diosgenin C-17 hydroxylation (**Fig 4 D, E and H**) was found for the saponins preferentially accumulated in root tissues, suggesting a root specific CYP450 activity. We also characterized a saponin core that is not oxidized at the C-16 position (**Fig 5E**), which might derive from a precursor or side product of the diosgenin biosynthetic pathway ^19^. Saponins with this core were seen across the cultivars analyzed but were more abundant in lowland shoot especially at a later developmental stage. This indicates the switchgrass CYP450 responsible for the cholesterol C-16 oxidation is possibly regulated in an *ecotype × tissue × development* manner in switchgrass.

The switchgrass saponins we characterized were either singly glycosylated at C-3 (**Fig 4 C, E, G and H**) or glycosylated at both C-3 and on the C-26 sidechain (**Fig 4 B, D and F**). Glycosylation diversity also comes from variation in the conjugating saccharide types, with one to six monosaccharides observed (**Table S14**). MS neutral mass losses corresponding to anhydrous glucose/galactose, rhamnose and xylose were observed for these saponins, and some monosaccharides were also acetylated (**Table S14**). Published studies indicated that the conjugated sugar moieties impact saponin bioactivities; for example, the oat avenacoside steroidal saponins are activated by deglycosylation upon leaf damage or pathogen attack ^39,40^, while antimicrobial activities of other saponins seem to rely on glycosylation ^30^. The purified switchgrass saponins with distinct glycosylation patterns provide good opportunities to perform structure-function analysis using an *in vitro* microbial bioassays.

The total saponin levels in roots of three lowland switchgrass cultivars, Alamo, Kanlow and BoMaster, are 3.5, 5.1 and 4.7 μg/mg dw respectively, which are higher than the three upland cultivars, Dacotah, Summer and Cave-in-Rock, at 1.2, 1.5 and 1.6 μg/mg dw (**Fig 6C lower panel and Table S22**). The root saponin contents in the lowland cultivars are close to those observed in legume roots: for example the *Medicago truncatula* ^41^ and two *Medicago sativa* cultivars, Radius ^42^ and Kleszczewska ^43^, contain 5.9, 5.0 and 9.3 μg/mg dw saponins in their roots, respectively.

Information about accession and ecotypic differences in saponin content could provide tools for improvement of switchgrass biomass. For example, avenacins have well documented protective roles protecting oat roots from the fungal ‘take-all’ disease ^6–8^. Saponin root differential accumulation among switchgrass ecotypes suggests that they might be attractive targets for breeding cultivars with an increased ability to improve yield by modulating microbiome structure and function ^44–46^. In contrast, breeding for low leaf saponins might produce varieties with biomass that is efficiently converted into fuel by avoiding accumulation of toxins that interfere with growth of processing microbes. Taken together, our results provide opportunities to identify targets for producing switchgrass varieties with improved plant/microbiome traits, increased biomass yield or biofuel conversion at lower economic and environmental costs.

## Materials and Methods

### Plant material

The six switchgrass cultivars used in this study were the upland ecotypes Dacotah, Summer and Cave-in-Rock and lowland ecotypes Alamo, Kanlow and BoMaster. The seeds were ordered from Native Connections (http://nativeconnections.net, Three Rivers, MI). The plants were grown under controlled growth conditions: temperature set at 27 °C with 16 h light (500 μE m^−2^s^−1^) per day and relative humidity set to 53%. Seeds were sown directly in a 1:1 mixture of sand and vermiculite, watered twice a week with deionized water and fertilized once every two weeks using half-strength Hoagland’s solution ^47^.

For untargeted LC-MS analysis, plant tissues were harvested at one-, two- and three-months after imbibition, corresponding to the vegetative, transition and early reproductive developmental stages, respectively ^26^. Roots, rhizomes and shoot (**Fig S1 A**) were collected separately for all the switchgrass cultivars. For example, one sample represented a specific *cultivar (genotype) × developmental stage × tissue type* combination (**Fig S1 B**). There were three biological replicates from three independent plants with two exceptions: two samples each from two independent plants for vegetative phase Cave-in-Rock samples and early reproductive phase Alamo samples (due to sample loss). For the GC-MS quantification of sapogenins, samples were only collected from the 3-month-old (early reproductive phase) plants. All samples were immediately frozen in liquid nitrogen and stored at −80 °C until extraction.

### Metabolite extraction

All chemicals were obtained from Sigma Aldrich (St. Louis, MO) unless otherwise specified. The samples were frozen in liquid nitrogen and powdered using 15mL polycarbonate grind vial sets (OPS Diagnostics, Lebanon, NJ) on a Mini G high throughput homogenizer (SPEX SamplePrep, Metuchen, NJ). 500 mg of each sample was extracted at 4 °C overnight (14 – 16 hours) in 5 mL of 80% methanol containing 1 μM telmisartan internal standard. Extracts were centrifuged at 4000 g for 20 min at room temperature to remove solids. Supernatant from each sample was transferred to an HPLC vial and stored at −80 °C prior to LC-MS analysis. For GC-MS, 50 mg of lyophilized sample was extracted in 1 mL of 80% methanol following the workflow described for LC-MS sample preparation above, as described by Tzin et al. ^48^.

### UPLC-ESI-QToF-MS analysis

Reversed-phase Ultra Performance Liquid Chromatography – Positive Mode Electrospray Ionization – Quadrupole Time-of-Flight MS (UPLC-(+)ESI-QToF-MS) analyses were performed with a Waters Acquity UPLC system coupled to a Waters Xevo G2-XS quadrupole time-of-flight (QToF) mass spectrometer (Waters, Milford, MA). The chromatographic separations were performed using a reversed-phase, UPLC BEH C18, 2.1 mm × 150 mm, 1.7 μm column (Waters) with a flow rate of 0.4 mL/min. The mobile phase consisted of solvent A (10 mM ammonium formate/water) and solvent B (100% acetonitrile). The column oven was maintained at 40 °C. Separations were achieved utilizing a 20-min method, injecting 10 μL of extract and using the following method (%A/%B): 0-1.0 min hold (99/1), linear gradient to 15 min (1/99), hold (1/99) until 18 min, returning at 18.01 min (99/1) and holding until 20 min. The Xevo G2-XS QToF was operated using the following static instrument parameters: desolvation temperature of 350 °C; desolvation gas flow rate at 600 L/h; capillary voltage of 3.0 kV; cone voltage of 30 V. Mass spectra were acquired in continuum mode over *m/z* 50 to 1500 using data-independent acquisition (DIA, MS^E^) or data-dependent MS/MS acquisition (DDA), with collision potential scanned between 20 – 80 V for the higher-energy function for DIA (and 20 – 60 V for DDA). The DDA mode automatically selected the three most abundant molecular ions to pass through the mass filter for fragmentation analysis at each scan. The MS system was calibrated using sodium formate, and leucine enkephalin was used as the lock mass compound but automated mass correction was not applied during DIA data acquisition. QC and reference samples were analyzed every 20 injections to evaluate the stability of the LC-MS system.

### Data processing and metabolite mining for the untargeted metabolomics analysis

Acquired raw MS data were processed using the Progenesis QI software package (v.3.0, Waters, Milford, MA) using retention time (RT) alignment, lock mass correction, peak detection, adduct grouping and deconvolution. The identified compounds were defined by the RT and *m/z* information and we also refer to these as *features*. The parameters used with Progenesis processing were as follows: sensitivity for peak picking, default; minimum chromatographic peak width, 0.15 min and RT range, 0.3 to 15.5 min. Intensities (ion abundances) of all the detected features were normalized to the internal standard, telmisartan, before downstream statistical analyses. Online databases – including KEGG, MassBank, PubChem and MetaboLights – were used to provide annotations to the features based on 10 *ppm* precursor tolerance, 95% isotope similarity and 10 *ppm* theoretical fragmentation pattern matching with fragment tolerance.

The complementary method Relative Mass Defect (RMD) filtering ^27^ was used to assign chemical classes for the features. Briefly, an RMD value of each feature was calculated in *ppm* as (mass defect/measured monoisotopic mass) × 10^6^. This value reflects the fractional hydrogen content of a feature and provides an estimate of the relative reduced states of carbons in the metabolite precursor of that feature. For example, in this study, we defined terpenoid glycosides (RMD of 350-550) or phenolics (RMD of 200-350) using this method. Features with RMD > 1200 are likely contaminants (e.g. inorganic salts) in the MS system.

The DDA was carried out for a set of pooled samples to generate positive mode MS/MS spectra for the abundant ions. Specialized metabolite discovery was performed by mining the DDA data and beginning with the most abundant metabolites. Characteristic precursor/fragment ions and RMD were used to assign metabolites to a particular chemical class (e.g. flavonoid glycoside, diterpenoid, sesquiterpenoid and saponin).

### Saponin purification

The switchgrass (Kanlow) plants were grown in a growth chamber using the conditions described in the section about Plant material. About 150 – 200 g fresh root or shoot tissues from fully matured plants (3 months post-germination) were harvested. The tissues were ground into powders with liquid nitrogen and placed in a 2 L beaker. 1.5 – 2 L of 80% methanol (in water) was added, and the mixture was incubated for 48 hours at 4°C. The mixture was centrifuged at 4000*g* for 15 min to remove insoluble debris. The supernatant volume reduced under vacuum using a rotary evaporator, followed by evaporation to dryness using a SpeedVac vacuum concentrator.

The residue was redissolved in 100 mL water. The liquid – liquid phase partitioning was carried against firstly hexane and then ethyl acetate, with a 1: 1 ratio, to remove the non-polar interfering metabolites. The resultant water phase was then loaded to a 35 cc C18 SPE cartridge (Waters). The cartridge was washed using 3 times of each 0%, 10%, 20% and 50% methanol (in water) to remove the polar interfering metabolites. The cartridge was eluted using 3 times of each 70%, 80% and 90% methanol (in water) to obtain the saponin-enriched fractions (eluates). Solvent was evaporated to dryness under vacuum using SpeedVac, and residue was redissolved in 8 mL of 80% methanol (in water). The insoluble was removed by centrifugation at 4000*g* for 5 mins at 25°C. Supernatants were transferred to autosampler vials.

Purification was carried out using Waters 2795 pump/autosampler connected with LKB Superrac 2211 fraction collectorand a Waters Symmetry C18 HPLC column (100 Å, 5μm, 4.6 mm × 150 mm). The mobile phase consisted of 0.15% formic acid in water, pH 2.8 (Solvent A) and acetonitrile (Solvent B). The linear gradient elution used to purify the saponin SS1244, SS1031, SS1064 and SS1032 from root tissues was 1% B at 0 min, 30% B at 1.01 min and linear-increased to 40% B at 7 min, 50% B at 7.01 min and linear-increased to 70% B at 15 min, 99% B at 15.01 min and held at 99% B between 15.01 and 18 min. A slightly modified linear gradient elution was used to purify the saponins SS1050, SS1098 and SS1254 from shoot tissues, 1% B at 0 min, 30% B at 1.01 min and linear-increased to 40% B at 8 min, 50% B at 8.01 min and linear-increased to 60% B at 15 min, 99% B at 15.01 min and held at 99% B between 15.01 and 18 min. The solvent flow rate was 1.5 mL/min and the column temperature was 40°C. Eluate was collected for every 10 sec as one fraction for each injection, using an injection volume of 100 μL. The HPLC fractions containing the seven targeted saponins were estimated to be > 75% pure based on LC-MS analysis results. Fractions of adequate purity for the same saponins were pooled.

### NMR Spectroscopy

NMR spectra of purified saponin samples were acquired using a Bruker Ascend 600 MHz spectrometer (Bruker Biospin, Germany) operating at 600.13 MHz for proton and equipped with an inverse 1.7 mm TCI micro Cryoprobe and a SampleJet auto-sampler unit. For ^1^H NMR spectra, solvent suppression with shaped pulse program (wetdc) was used with a scan number of 16 at temperature 298K with pulse width 10.75 μs and power 0.2 W. Acquisition time for each scan was 0.681 s with a delay time 3 s for a spectral width of 20 ppm. For locking the magnetic field CD_3_OD was used as the solvent and ^1^H spectra were calibrated using the residual solvent peaks. Baseline and phase correction were performed manually using Bruker Topspin 3.5.6 software. Subsequently, 2D NMR spectra were collected using default Bruker pulse program cosygpmfppqf for COSY (ns=16, sw= 13 for F1 & F2), hsqcedetgpsp.3 for HSQC (ns=64, sw= 13 for F2 and 220 for F1), hmbcgpndqf for HMBC (ns=128, sw=13 for F1 & F2) and mlevphpr.2 for TOCSY (ns=64, sw= 13 for F1 & F2) with pulse width 10.75 μs and power 0.2 W for 1H and pulse width 12 μs and power 68W for 13C. DEPT-Q spectra were collected using a Bruker Avance III 800 MHz spectrometer equipped with a 5 mm TCI cryoprobe. Data were obtained using deptsp135 pulse program with 3072 scan for a sw 222 ppm and 0.734 s acquisition time for each scan with pulse width 14 μs and power 107.15 W for 13C. Finally, the data were visualized with both Topspin 3.5.6 and MestReNova software for peak assignments.

### Analysis of switchgrass sapogenins by acid hydrolysis, derivatization and GC-MS

To analyze sapogenins, acid hydrolysis was carried out according to a previously published protocol ^49^. In brief, 300 μL of switchgrass extract, 200 μL of distilled water and 100 μL of 12 M hydrochloric acid (MilliporeSigma, Burlington, MA) were mixed in a polypropylene microcentrifuge tube and incubated at 85°C for 2 h. The samples were cooled and evaporated to dryness under vacuum with the temperature ≤ 40 °C. The resultant pellet was dissolved in 500 μL distilled water and extracted with 500 μL ethyl acetate for phase partition. After this, 300 μL of the ethyl acetate layer was transferred to a new microcentrifuge tube and evaporated to dryness under vacuum at room temperature. The dry residue was dissolved in 100 μL *N*-methyl-*N*-(trimethylsilyl)trifluoroacetamide (MSTFA), derivatized overnight at 60 °C and analysed using a 30 m VF5 column (Agilent Technologies, Santa Clara, CA; 0.25 mm ID, 0.25 μm film thickness) coupled to an Agilent 5975 single quadrupole MS (Agilent Technologies, Santa Clara, CA) operated using 70 eV electron ionization (EI). Then, the same set of samples were analysed on an Agilent 7010B triple quadrupole MS using the same column for chemical ionization (CI). The MS scanning range was *m/z* 80 – 800. Splitless sample injection was used, with helium as carrier gas at constant flow of 1 mL/min and the inlet and transfer line held at 280 °C. The GC temperature program was as follows: held at 50 °C for 1 min; ramped at 30 °C /min to 200 °C; ramped at 10 °C /min to 320 °C and held for 10 min.

Absolute and relative quantification of the sapogenins were performed using commercially available diosgenin (~95%, Sigma Aldrich) as an external standard. Serially diluted standards (6 – 192 μg/mL dissolved in 80% ethanol) were pre-treated in the same way as the other samples: after hydrochloric acid hydrolysis, four target peaks were identified that are derived from the commercial diosgenin. They were termed as diosgenin standard (DS) 1 – 4. The DS4, eluting at 21.3 minutes, could be detected by EI GC-MS at high standard concentrations (**Fig S3A**). All target peaks were combined when plotted against the standard’s concentrations to generate six-point response curve. Duplicate technical replicate analyses were done for each standard sample used to generate the standard response curve, which was linear (r^2^ > 0.97, **Fig S3B**), and was used to calculate relative concentrations for sapogenins detected in switchgrass extracts. Quantifications were based on the peak areas calculated from total ion chromatograms for standards, diosgenin-type sapogenins and the unknown sapogenins with chemical structures similar to diosgenin. Six individual plants were harvested for each switchgrass genome type and pooled into three groups of two individual plants. Pooling permitted collection of enough tissue to perform separate analysis of leaf blade, leaf sheath, stem, rhizome and root to overcome the issue of limited amount of plant tissue.

### Statistical analysis

To visualize the metabolomic variation in tissue types, developmental stages and ecotypes of switchgrass, Hierarchical Clustering Analysis (HCA) and Principal Component Analysis (PCA) were performed using R (v. 3.5.1) and MetaboAnalyst 5.0 online tool platform ^50^. Signals were normalized to internal standard area and tissue mass, log-transformed and scaled by Pareto scaling prior to these analyses. To assess the relationship among samples and among features, hierarchal clustering with Euclidean distance as the similarity measure and Ward.D2 as the clustering algorithm was used. The relationship results were visualized in the form of dendrograms on the heatmap. Significance analyses were carried out using the Progenesis QI software (Waters) to identify the differentially accumulated features (DAFs) between the upland and lowland ecotypes. The cutoff threshold of the significance analyses was FDR adjusted Student’s t-test *p* ≤ 0.05 and fold-change ≥ 2. Results of the analyses were visualized by volcano plots. To examine statistical differences in the sapogenin concentrations among samples, Kruskal-Wallis test and Post-Hoc Dunn’s tests were performed in R. *p* ≤ 0.05 was considered statistically significant.

## ACKNOWLEDGMENTS

We thank Gregory Bonito (Michigan State University) for providing the switchgrass seeds used in this study and helpful advice on the manuscript; Anthony Schilmiller and Cassandra Johnny (MSU Mass Spectrometry and Metabolomics Core) for LC- and GC-MS analysis related technical support. This material is based upon work supported by the Great Lakes Bioenergy Research Center, U.S. Department of Energy, Office of Science, Office of Biological and Environmental Research under Award Number DE-SC0018409.

## Notes

### Competing Interest Statement

The authors have declared no competing interest.

### Summary of Updates

We added NMR results and re-annotation of LC-MS data.

