## Supplementary Figures for "Switchgrass metabolomics reveals striking genotypic and developmental differences in specialized metabolic phenotypes"

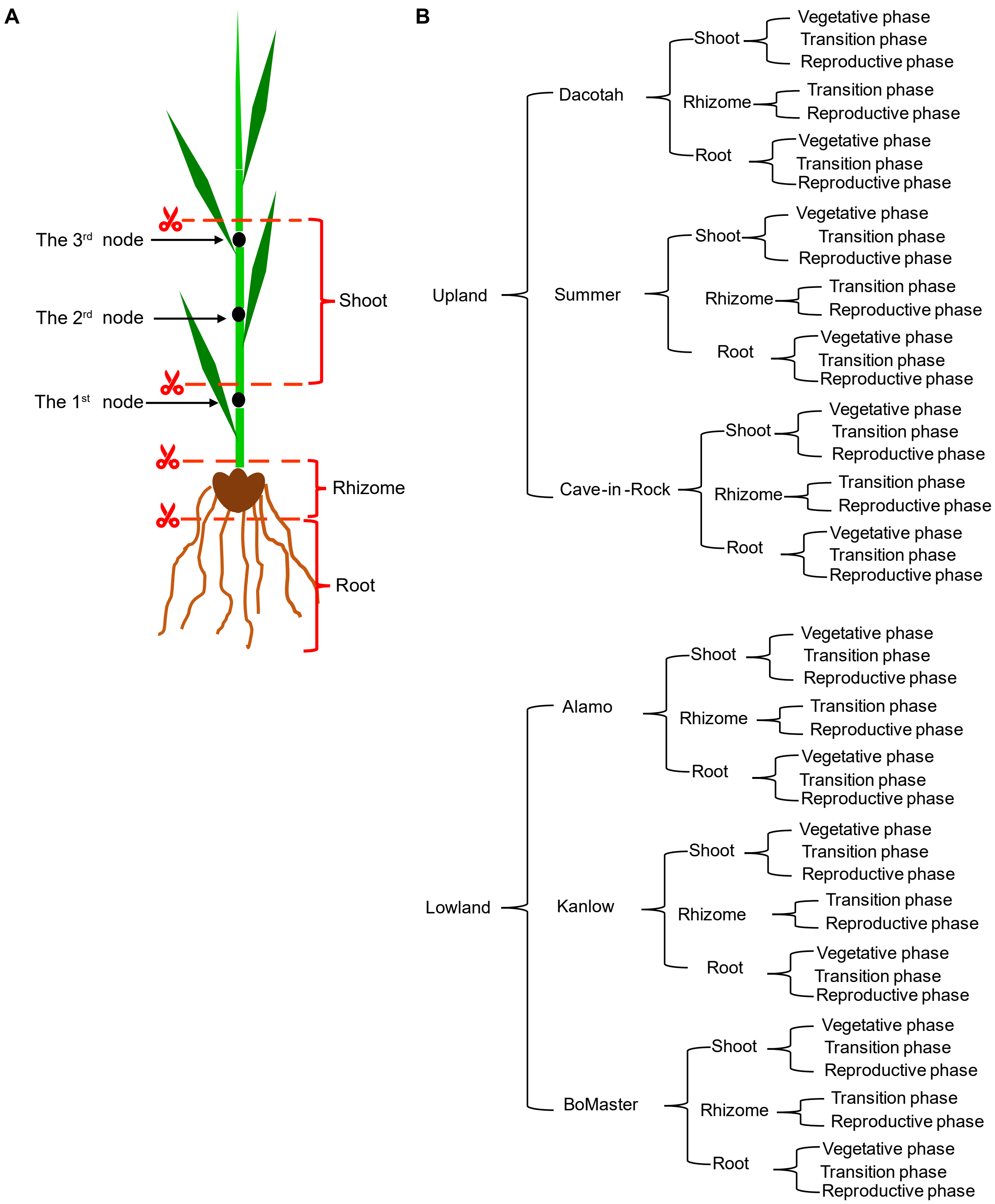


**Figure S1. The method for sample collection and sample panel involved in this study.** (A) Developmental stage identifications and sample collections were performed according to Hardin et al. (2013). The diagram shows how the shoot, rhizome and root tissues were collected from the plants at the transition- or reproductive-phase. The entire above- or below-ground parts were collected for the plants at the vegetative-phase. (B) The sample panel in this study contains six switchgrass cultivars, three tissue types and three developmental stages. Rhizomes were not collected for the plants at the vegetative-phase, due to the small size. In total, there are 48 samples groups in this study.

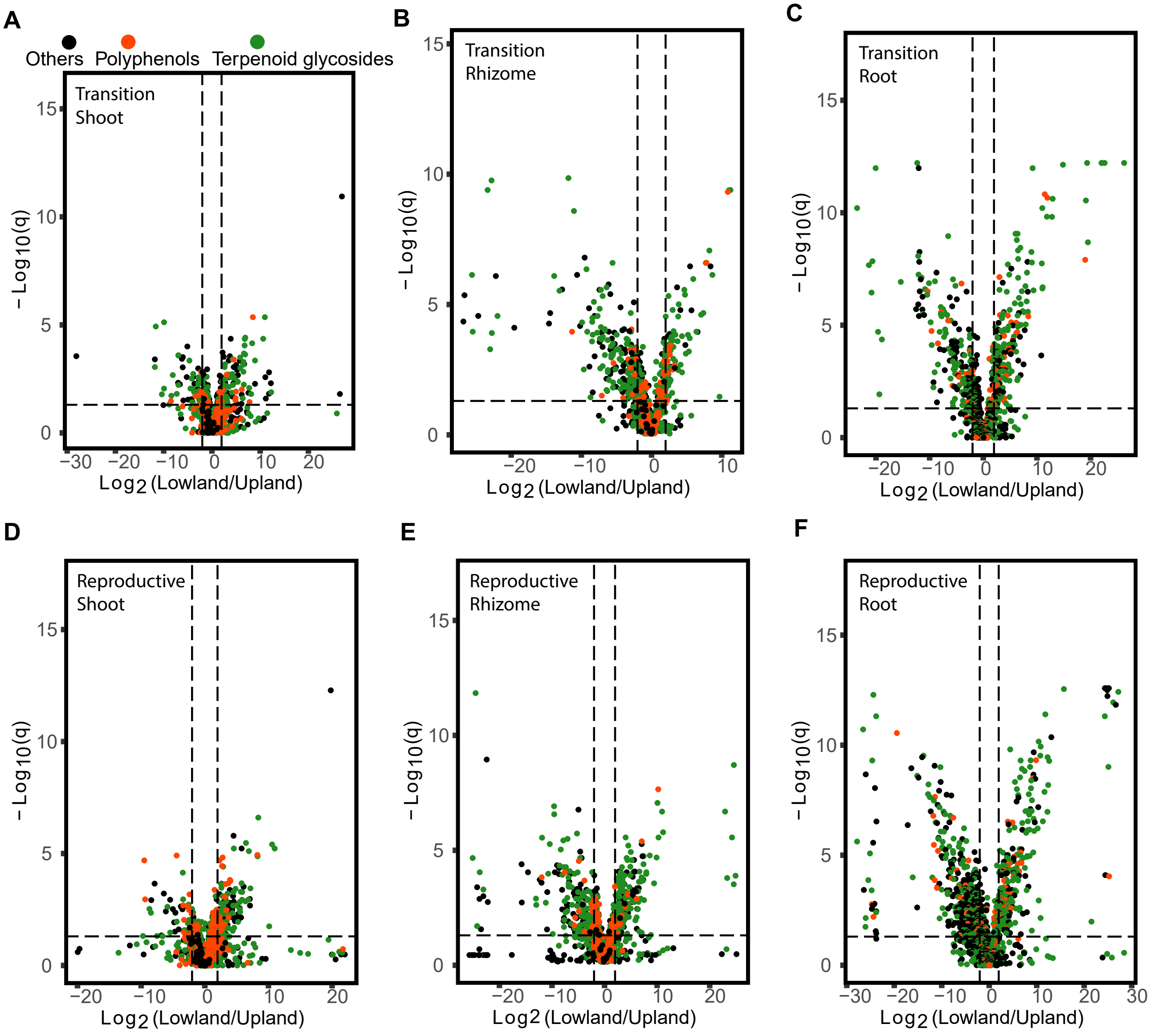


**Figure S2. Significance analysis for the differentially accumulated features (DAFs) between the upland and lowland switchgrass ecotypes for each given *tissue type x development stage* combination.** (A) *Shoot x Transition phase*. (B) *Rhizome x Transition phase*. (C) *Root x Transition phase*. (D) *Shoot x Reproductive phase*. (E) *Rhizome x Reproductive phase*. (F) *Root x Reproductive*. The horizontal dash line stands for the adjusted *p*-value = 0.05. Vertical dash lines stand for the $\log_2$ value of the fold-change = 2. Metabolite symbols are color-coded based on relative mass defect (RMD) analysis.

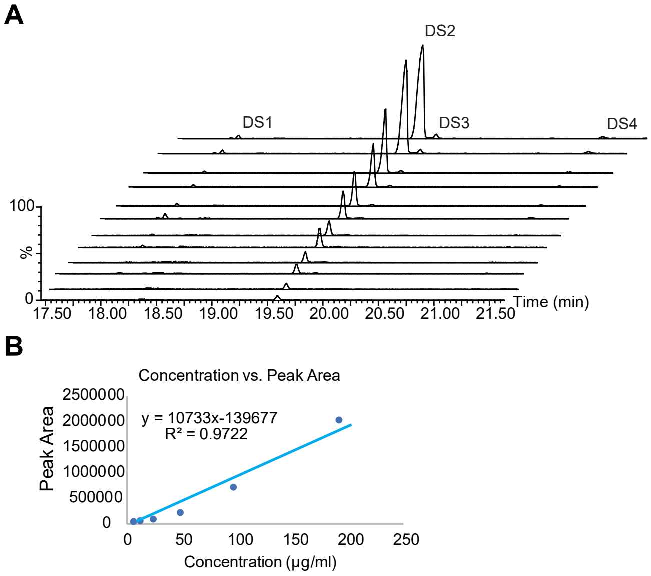


**Figure S3. The six-point instrument response curve over a range of 6 – 192 mg/ml by serial dilution of the diosgenin standard.** (A) Extracted ion (*m/z* 139) chromatograms of duplicate standard injections from the standard concentration series (6, 12, 24, 48, 96 and 192 g/ml). One major (DS2) and three minor (DS1, DS3 and DS4) diosgenin standard derived peaks (due to the sample treatment, **Materials and Methods)** were identified. Areas for the four peaks were combined when plotted against the concentration. (B) Reproducibility and linearity (B, $r^2 > 0.97$) of the method.

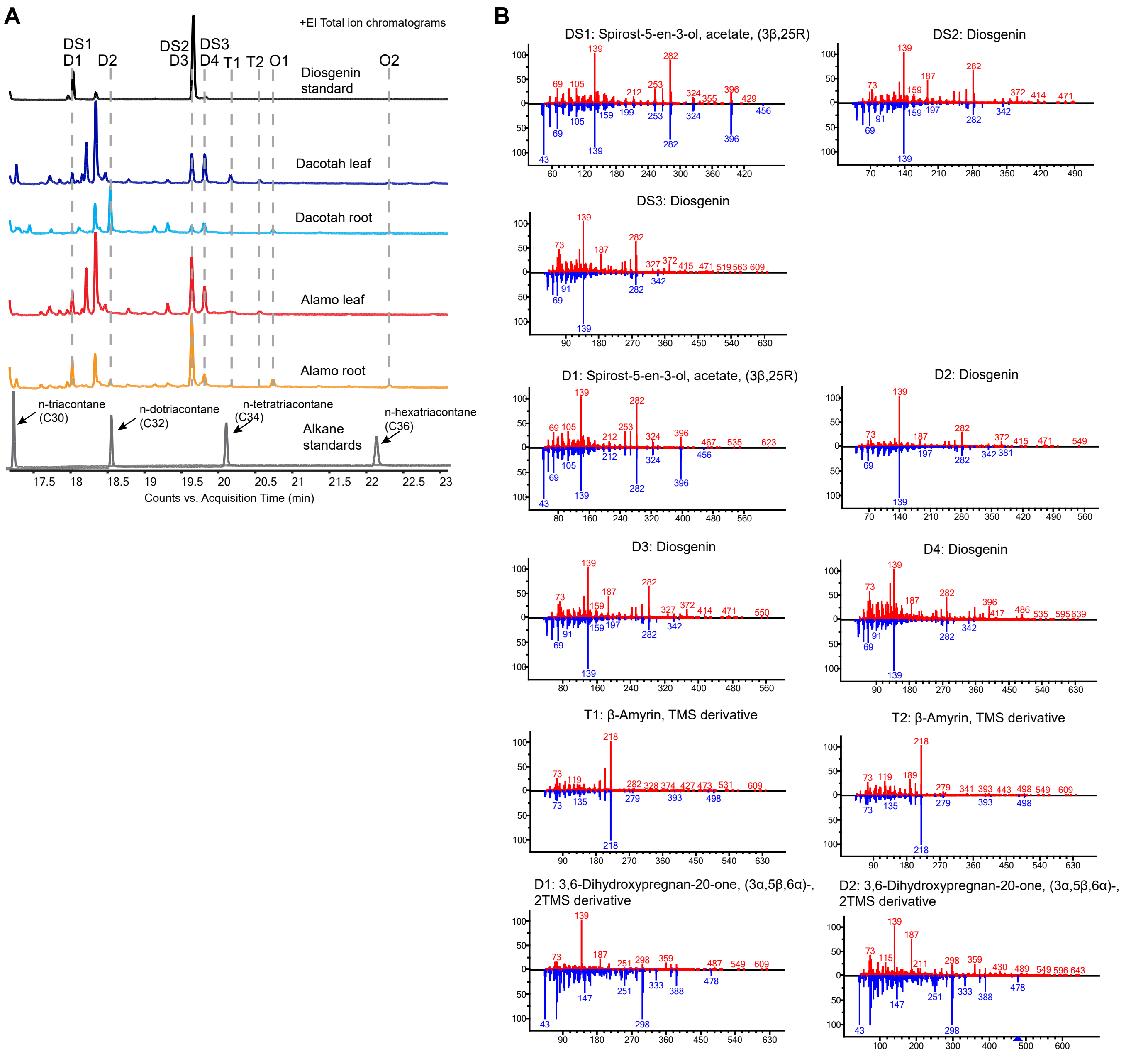


**Figure S4. Sapogenin aglycone peaks identified in switchgrass extracts by (EI) GC-MS**. (A) +(EI) GC-MS total ion chromatograms (TICs) of the extracts of Dacotah leaf (dark blue), Dacotah root (blue), Alamo leaf (red), Alamo root (orange) as well as a diosgenin standard sample (black) and an n-alkane standard mixture (gray) containing n-triacontane (C30), n-dotriacontane (C32) , n-tetratriacontane (C34) and n-hexatriacontane (C36) that was used for calculating the RIs. The identified potential sapogenin peaks in the switchgrass and standard samples are indicated and aligned by the dashed lines. Note that the D1, D3 and D4 peaks detected in switchgrass samples have slightly different RTs from the DS1, DS2 and DS3 peaks detected in the diosgenin standard sample (black). (B) The (EI) GC-MS spectral pattern (upper/red) for each potential sapogenin peak. The abundant fragment ions for each peak are listed in **Table 2**. The best NIST library matches (lower/blue) for the EI spectral patterns of the detected sapogenin peaks, based on the highest reverse match scores, are shown.

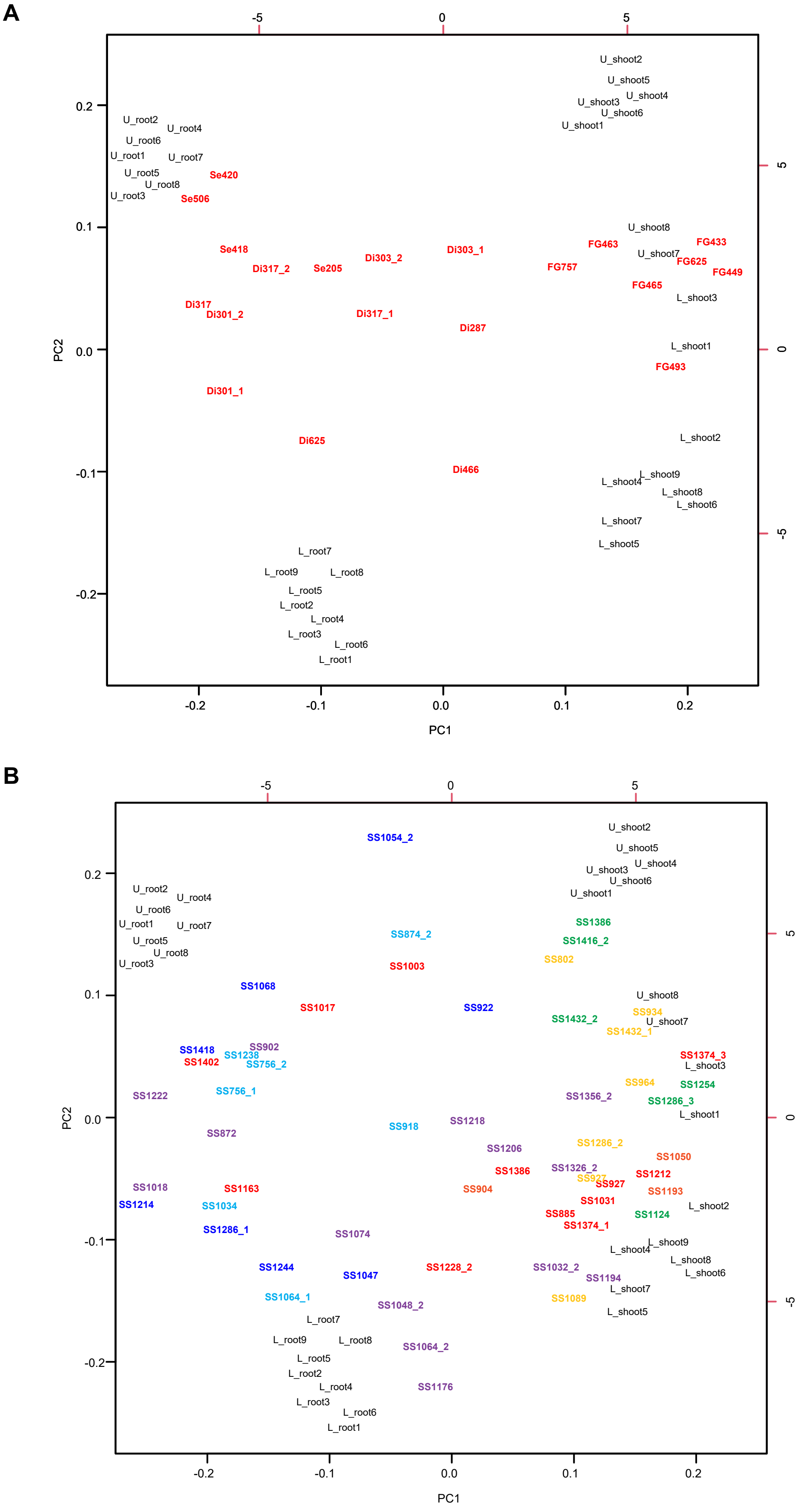
A
-5
0
5
U_shoot2
U_shoot5
U_shoot4
U_shoot3
U_shoot6
U_shoot1
U_root2
U_root4
U_root6
U_root1
U_root7
U_root5
U_root8
U_root3
Se420
Se506
U_shoot8
FG433
FG463
U_shoot7
FG625
FG449
Se418
Di303_1
Di317_2
Se205
Di303_2
FG757
FG465
L_shoot3
Di317
Di301_2
Di317_1
Di287
L_shoot1
FG493
Di301_1
Di625
L_shoot2
Di466
L_shoot9
L_shoot4
L_shoot8
L_shoot6
L_shoot7
L_shoot5
L_root7
L_root9
L_root8
L_root5
L_root2
L_root4
L_root3
L_root6
L_root1
0.2
0.1
0.0
-0.1
-0.2
PC2
-0.2
-0.1
0.0
0.1
0.2
PC1
B
SS1054_2
SS874_2
SS1003
SS1386
SS1416_2
SS802
SS1068
SS1017
SS922
SS934
SS1432_2
SS1432_1
SS1418
SS902
SS1238
SS1402
SS756_2
SS1374_3
SS1222
SS756_1
SS964
SS1254
SS1356_2
SS1286_3
SS918
SS1218
SS872
SS1206
SS1286_2
SS1050
SS1386
SS1326_2
SS1212
SS927
SS927
SS1193
SS1018
SS1163
SS904
SS1214
SS1034
SS885
SS1031
SS1124
SS1374_1
SS1286_1
SS1074
SS1244
SS1047
SS1228_2
SS1032_2
SS1194
SS1064_1
SS1048_2
SS1089
SS1064_2
SS1176

**Figure S5. Biplot of the PCA in Fig. 3C showing differential accumulations of the specialized metabolites among the four sample groups (upland shoot, upland root, lowland shoot and lowland root).** For an easier visualization, the biplot was separated into an upper panel for sesquiterpenoids, diterpenoids and flavonoid glycosides (A) and a lower panel for saponins (B). The saponins from different classes were colored to match with the color schemes in **Fig 4A** and **Table S21** (Red, D-415-SCG; Purple, D-415; Dark blue, D-431-SCG; Blue, D-431; Orange, D-417-SCG; Green, D-457; Yellow, D-473) For the class names, the letter 'D' stands for 'diosgenin'; numerical number reflects *m/z* value of aglycone fragment ion detected by positive mode MS; 'SCG' stands for 'side chain glycosylation'. The metabolites further away from the center are more preferentially accumulated in certain tissue types and/or ecotypes.

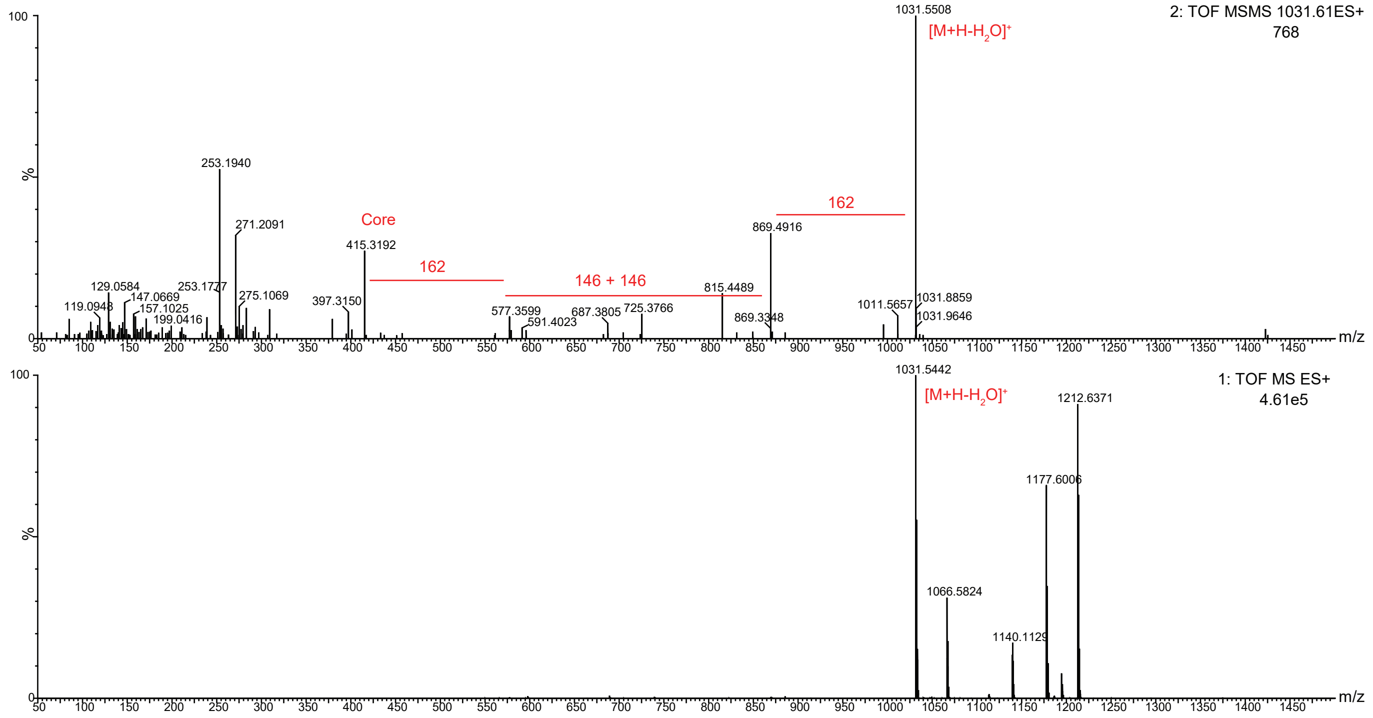


**Figure S6. Mass spectra of the switchgrass saponin SS1031 analyzed by high resolution ES+ LC-MS using DDA mode.** MS function: 1. survey scan; 2. DDA scan (MS/MS). The integer values of the MS neutral mass losses are labeled (18, loss of one molecule of water; 146, loss of one molecule of deoxyhexose; 162, loss of one molecule of hexose). See **Table S14** for additional details.

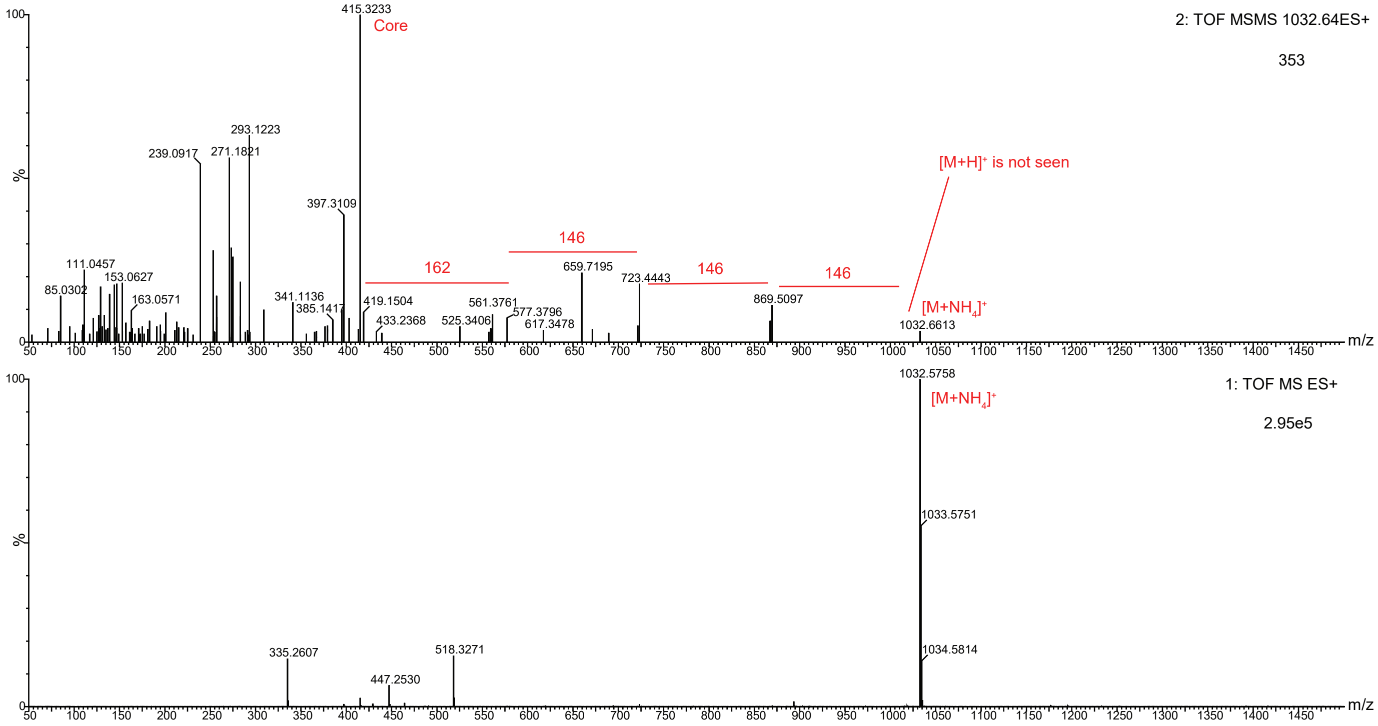


**Figure S7. Mass spectra of the switchgrass saponin SS1032 analyzed by high resolution ES+ LC-MS using DDA mode.** MS function: 1. survey scan; 2. DDA scan (MS/MS). The integer values of the MS neutral mass losses are labeled (18, loss of one molecule of water; 146, loss of one molecule of deoxyhexose; 162, loss of one molecule of hexose). See **Table S14** for additional details.

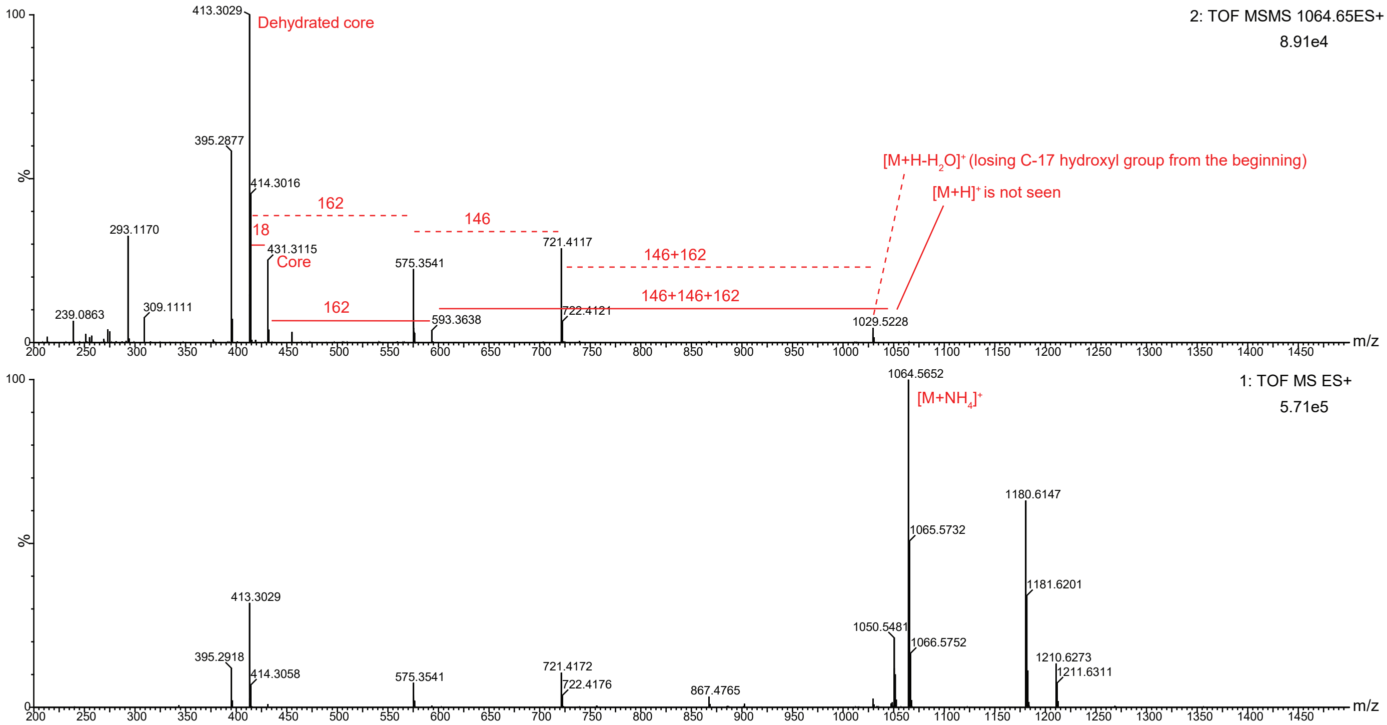


**Figure S8. Mass spectra of the switchgrass saponin SS1064 analyzed by high resolution ES+ LC-MS using DDA mode.** MS function: 1. survey scan; 2. DDA scan (MS/MS). The integer values of the MS neutral mass losses are labeled (18, loss of one molecule of water; 146, loss of one molecule of deoxyhexose; 162, loss of one molecule of hexose). See **Table S14** for additional details.

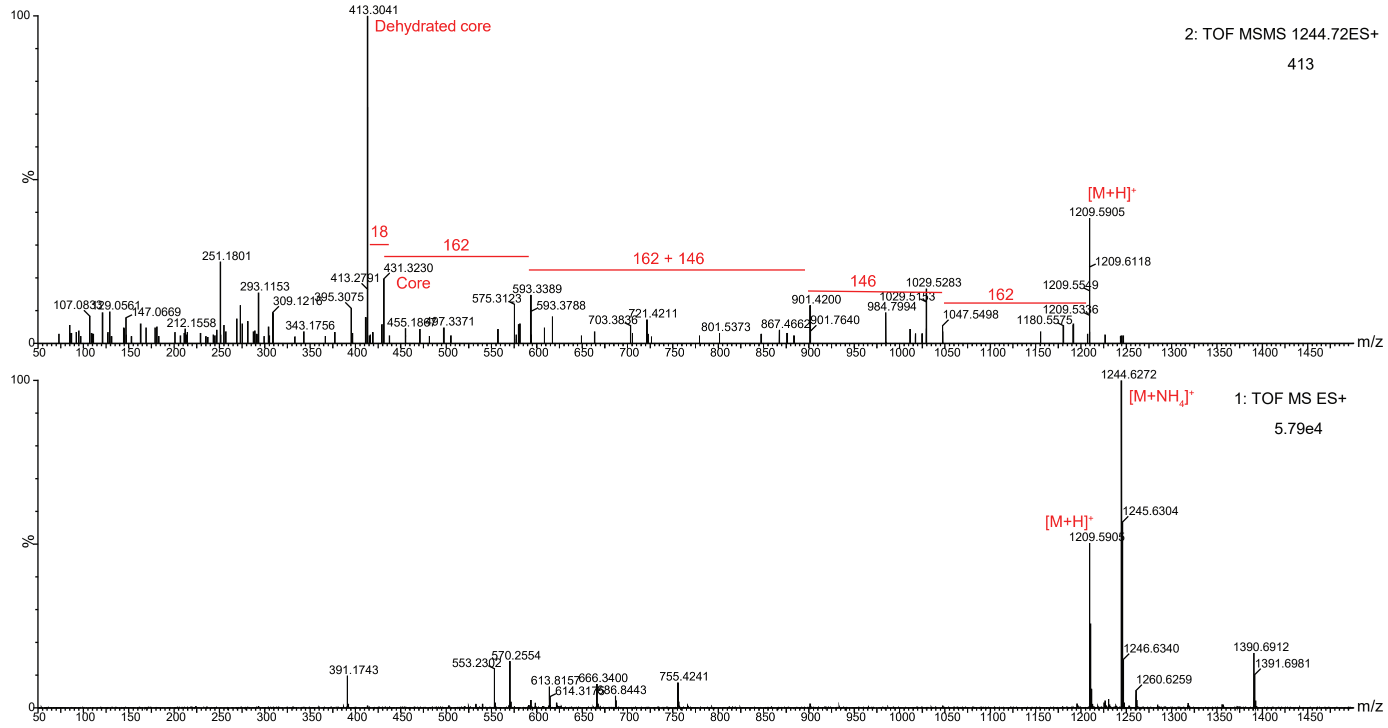


**Figure S9. Mass spectra of the switchgrass saponin SS10244 analyzed by high resolution ES+ LC-MS using DDA mode.** MS function: 1. survey scan; 2. DDA scan (MS/MS). The integer values of the MS neutral mass losses are labeled (18, loss of one molecule of water; 146, loss of one molecule of deoxyhexose; 162, loss of one molecule of hexose). See **Table S14** for additional details.

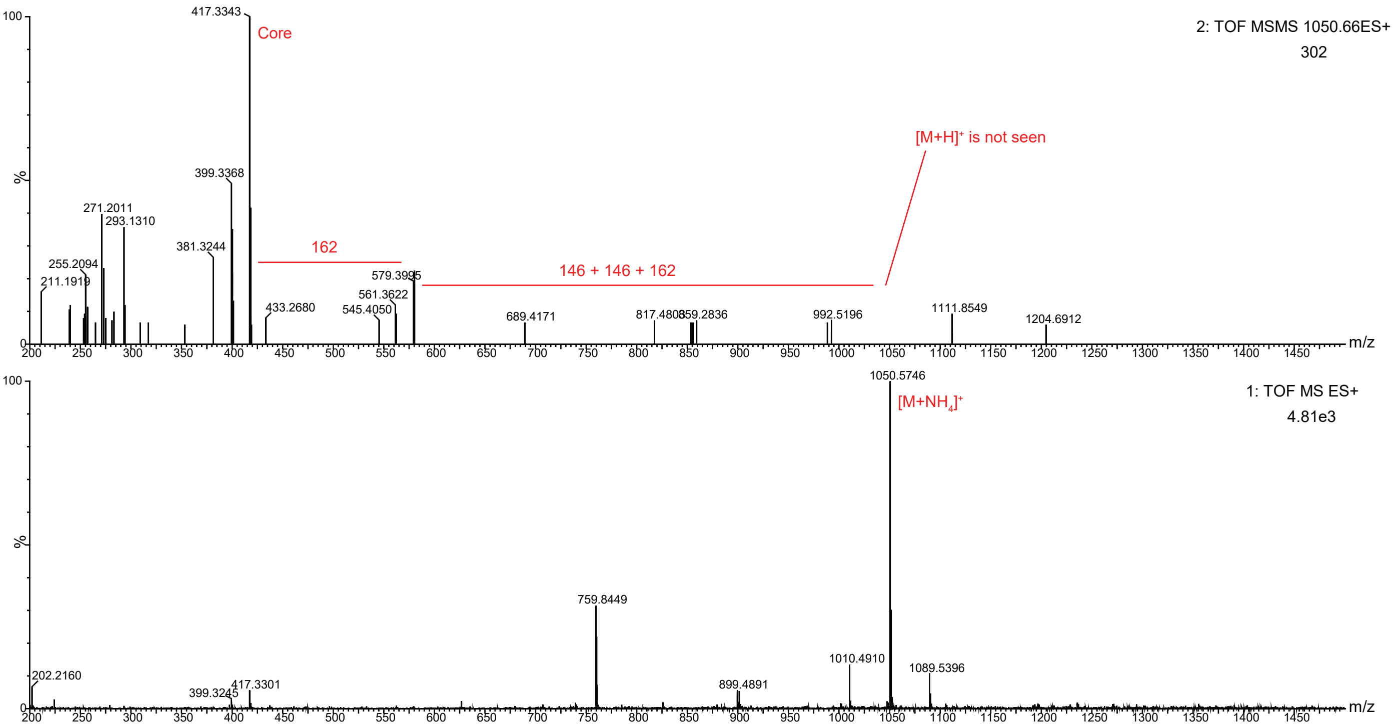


**Figure S10. Mass spectra of the switchgrass saponin SS1050 analyzed by high resolution ES+ LC-MS using DDA mode.** MS function: 1. survey scan; 2. DDA scan (MS/MS). The integer values of the MS neutral mass losses are labeled (18, loss of one molecule of water; 146, loss of one molecule of deoxyhexose; 162, loss of one molecule of hexose). See **Table S14** for additional details.

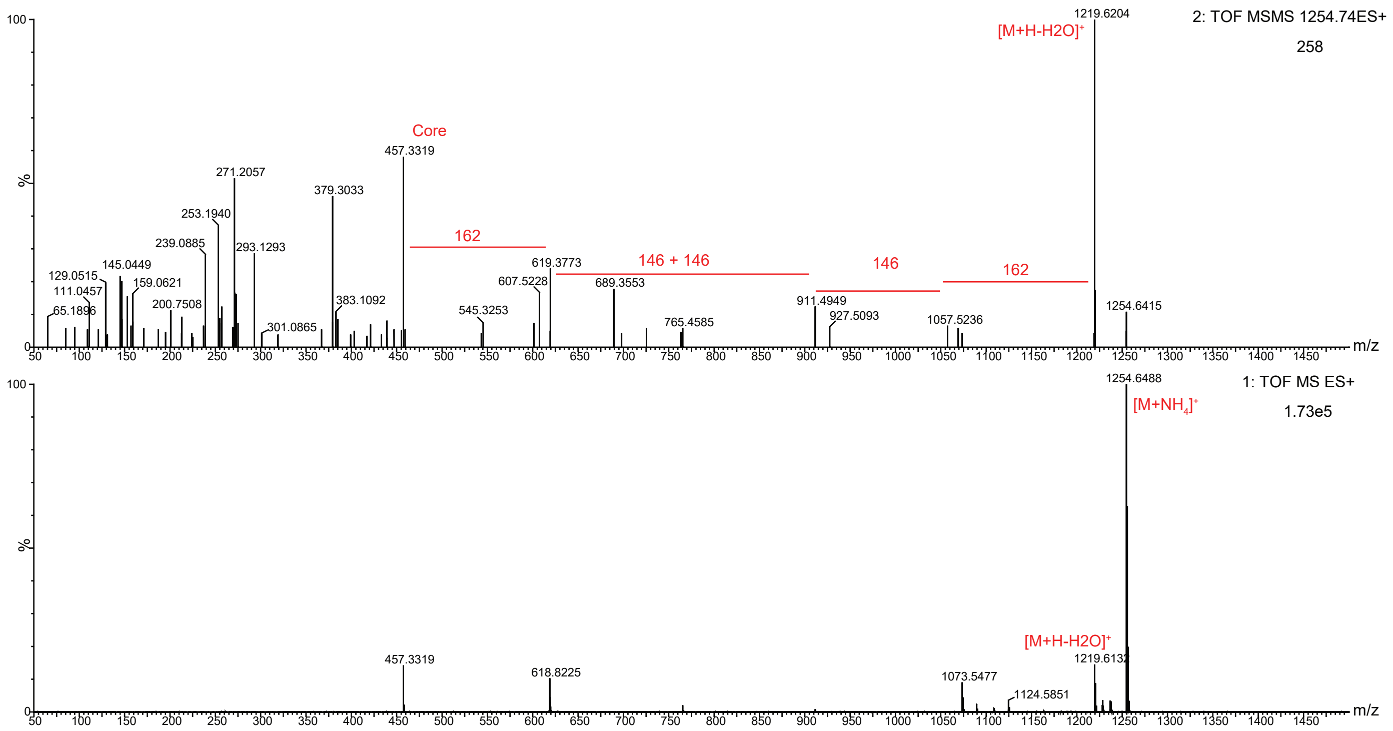


**Figure S11. Mass spectra of the switchgrass saponin SS1254 analyzed by high resolution ES+ LC-MS using DDA mode.** MS function: 1. survey scan; 2. DDA scan (MS/MS). The integer values of the MS neutral mass losses are labeled (18, loss of one molecule of water; 146, loss of one molecule of deoxyhexose; 162, loss of one molecule of hexose). See **Table S14** for additional details.

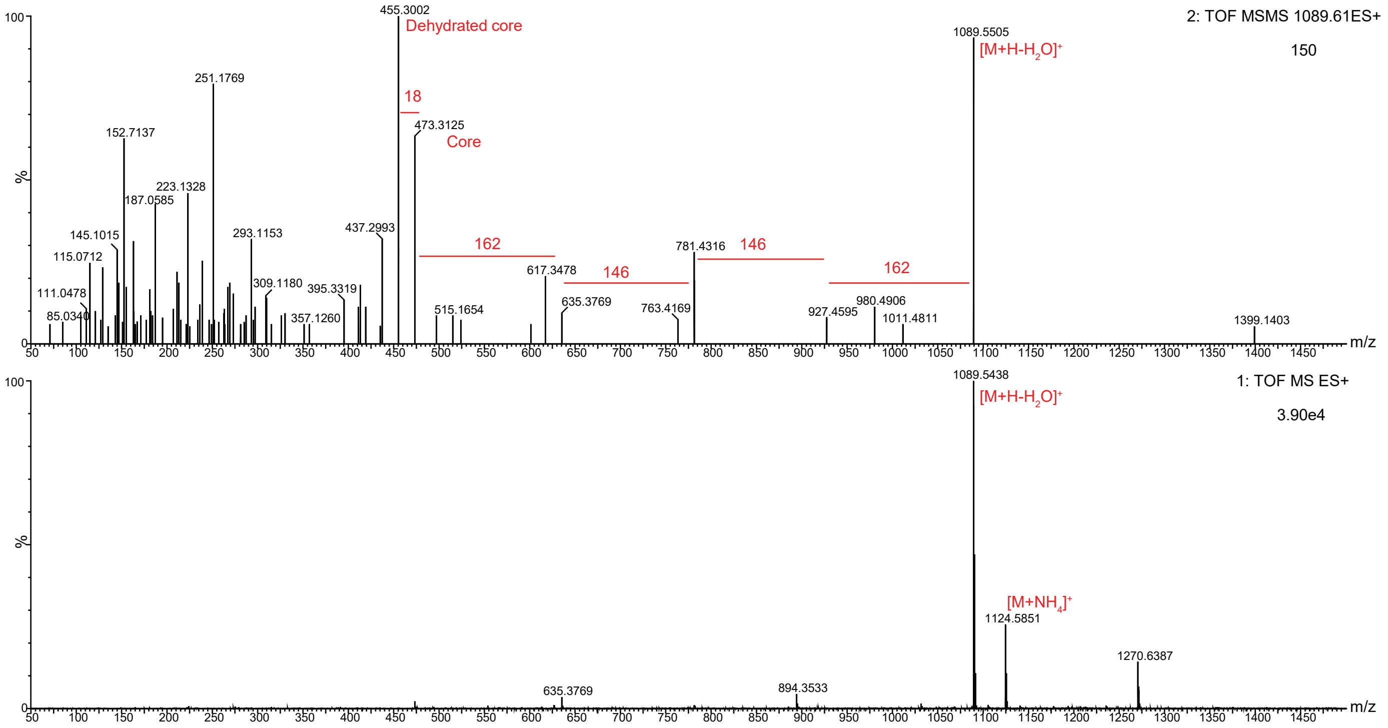


**Figure S12. Mass spectra of the switchgrass saponin SS1089 analyzed by high resolution ES+ LC-MS using DDA mode.** MS function: 1. survey scan; 2. DDA scan (MS/MS). The integer values of the MS neutral mass losses are labeled (18, loss of one molecule of water; 146, loss of one molecule of deoxyhexose; 162, loss of one molecule of hexose). See **Table S14** for additional details.

**Table S15. Chemical shift assignments of the aglycone of the switchgrass saponin SS1031. Chemical formular: $C_{27}H_{44}O_4$**

| Position | $^{1}H$ δ (ppm) | $^{13}C$ δ (ppm) | Annotation |
|---|---|---|---|
| 1 | 1.88/1.09 | 36.65 | CH2 |
| 2 | 1.92/1.64 | 29.35 | CH2 |
| 3[a] | 3.63 | 69.71 | CH |
| 4 | 2.46/2.31 | 37.17 | CH2 |
| 5[b] | | 139.71 | olefinic =C |
| 6 | 5.4 | 120.71 | olefinic =CH |
| 7 | 2/1.57 | 31.14 | CH2 |
| 8 | 1.69 | 30.71 | CH |
| 9 | 0.99 | 49.57 | CH |
| 10 | | 35.99 | quaternary C |
| 11 | 1.58/1.54 | 20.57 | CH2 |
| 12 | 1.79/1.22 | 38.85 | CH2 |
| 13 | | 40.49 | quaternary C |
| 14 | 1.16 | 56.0 | CH |
| 15 | 2.04/1.29 | 30.04 | CH2 |
| 16 | 4.58 | 80.0 | CH |
| 17 | 1.79 | 62.0 | CH |
| 18 | 0.85 | 14.85 | CH3 |
| 19 | 1.07 | 17.85 | CH3 |
| 20 | 2.12 | 38.85 | CH |
| 21 | 1.02 | 14.0 | CH3 |
| 22 | | 109.92 | quaternary C |
| 23 | 2.13/1.75 | 35.85 | CH2 |
| 24 | 1.94/1.64 | 28.14 | CH2 |
| 25 | 1.76 | 32.43 | CH |
| 26[c] | 3.75 | 74.0 | CH2 |
| 27 | 0.97 | 15.28 | CH3 |

[a] The C-3 carbon connects to a sugar anomeric proton, 5.2 *ppm*, through HMBC.
[b] The C-5 carbon chemical shift, 139.71 *ppm*, was determined by HMBC.
[c] The C-26 carbon connects to a sugar anomeric proton, 4.26 *ppm*, through HMBC.

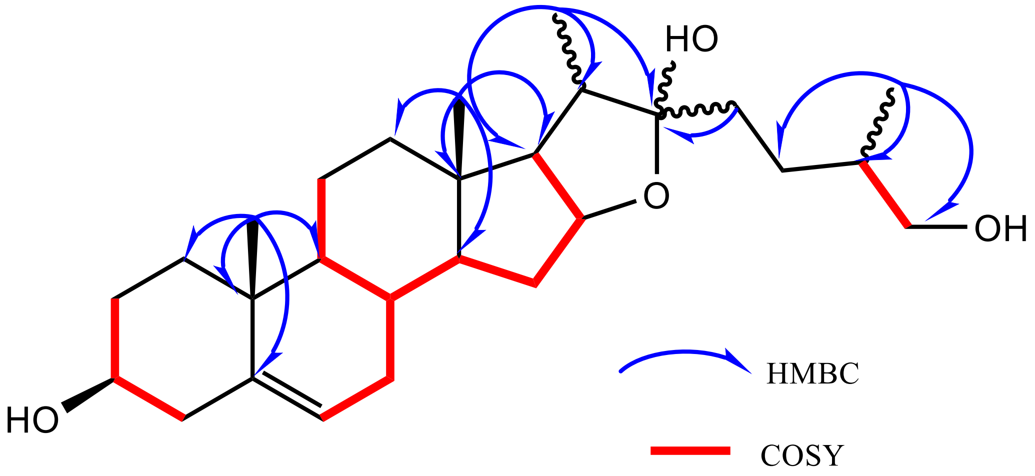


**Figure S13. The key COSY (red, bold) and HMBC (blue arrow) correlations for the aglycone of the switchgrass saponin SS1031.**

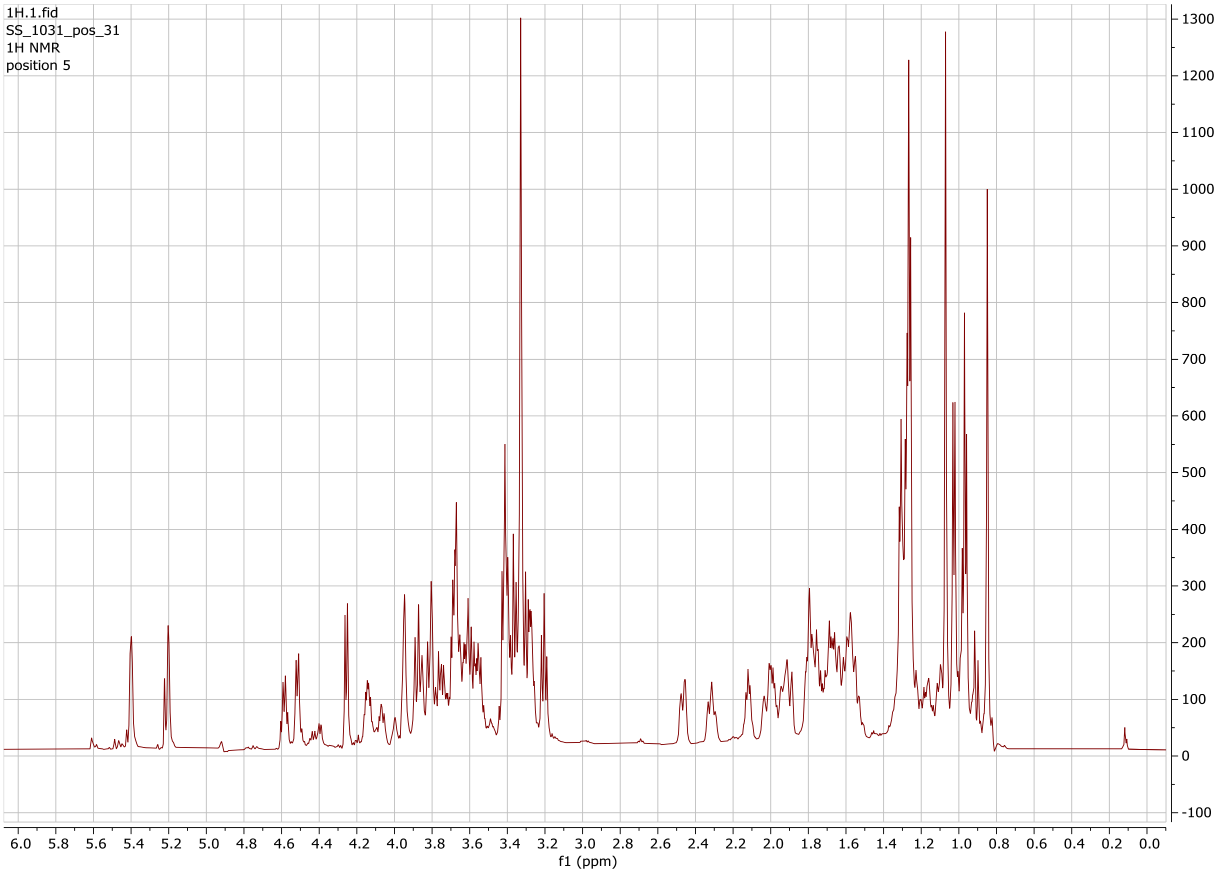


**Figure S14. $^1$H NMR spectrum for the intact molecule of the switchgrass saponin SS1031.**

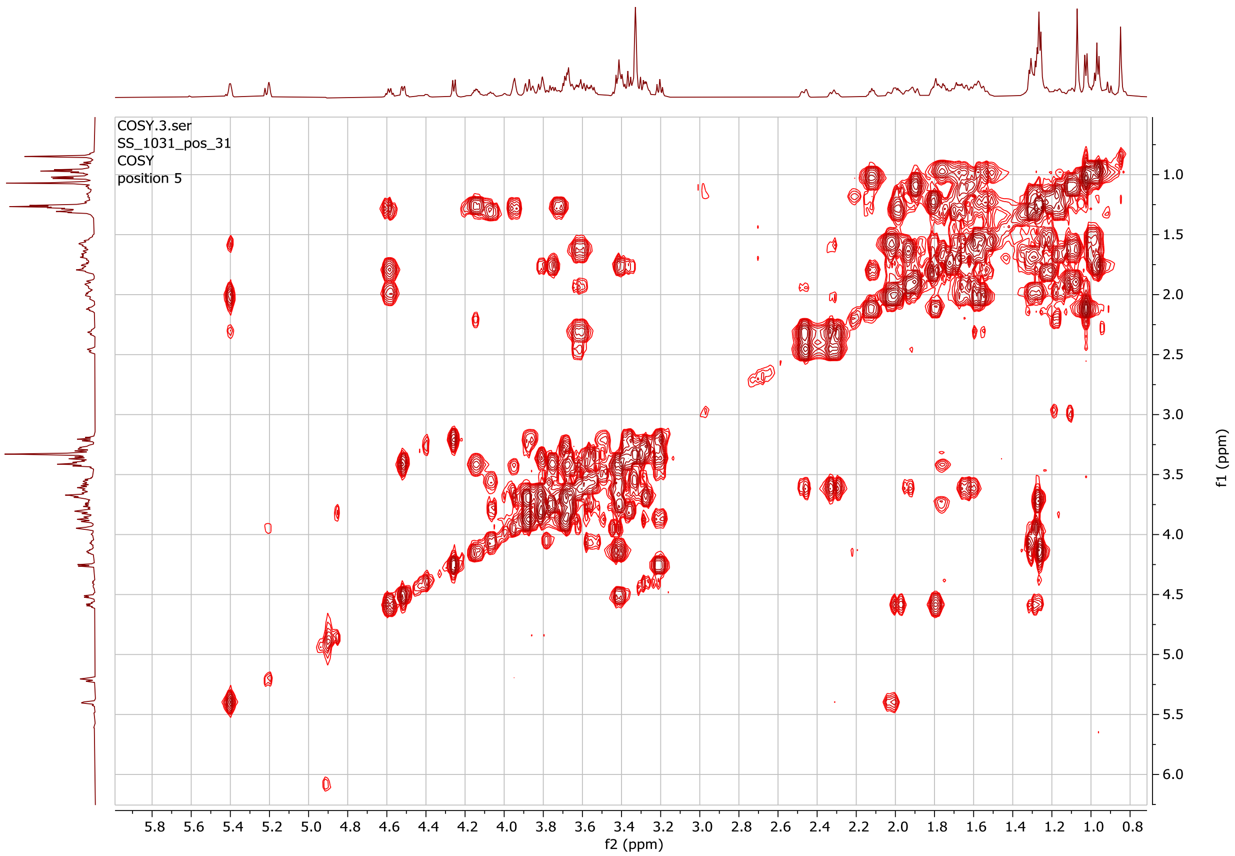


**Figure S15. gCOSY $^{1}$H-$^{1}$H NMR spectrum for the intact molecule of the switchgrass saponin SS1031.**

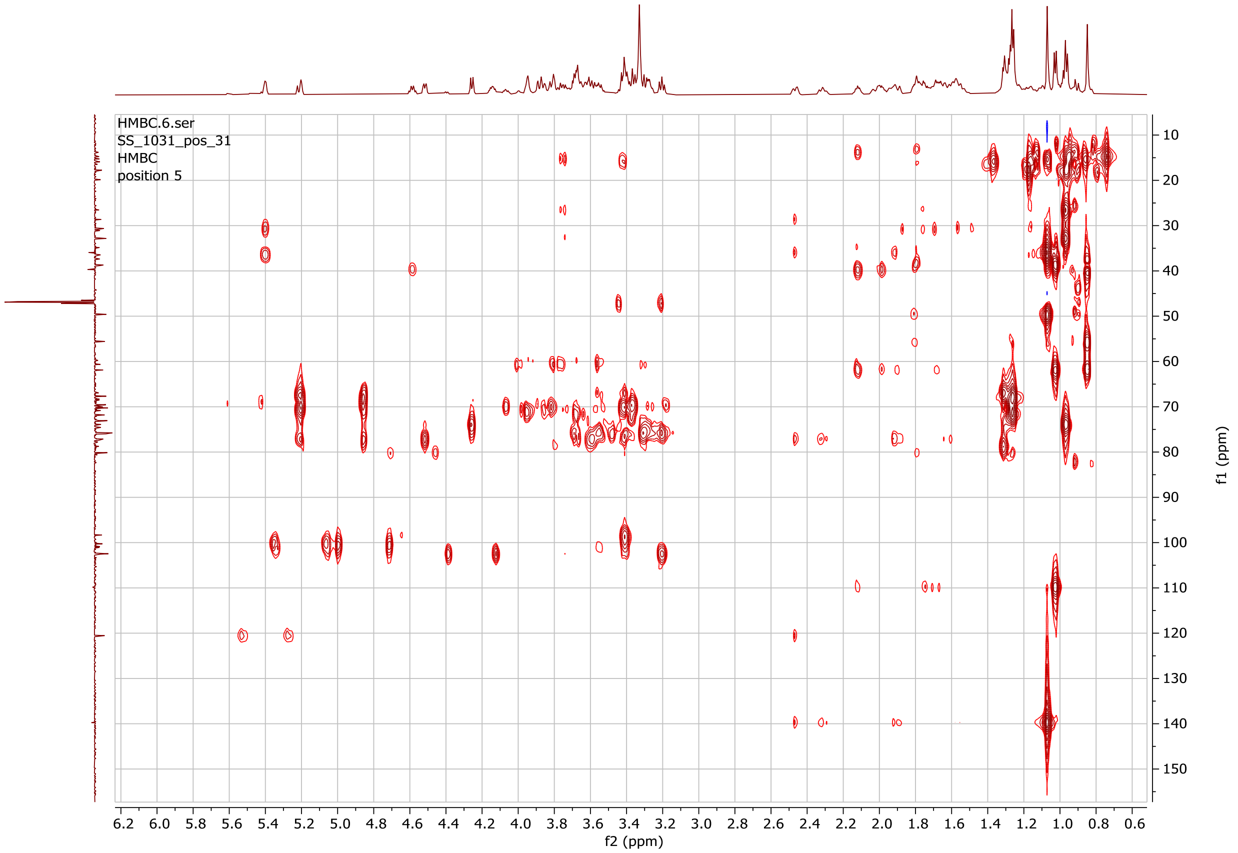


**Figure S16. gHMBCAD NMR spectrum for the intact molecule of the switchgrass saponin SS1031.**

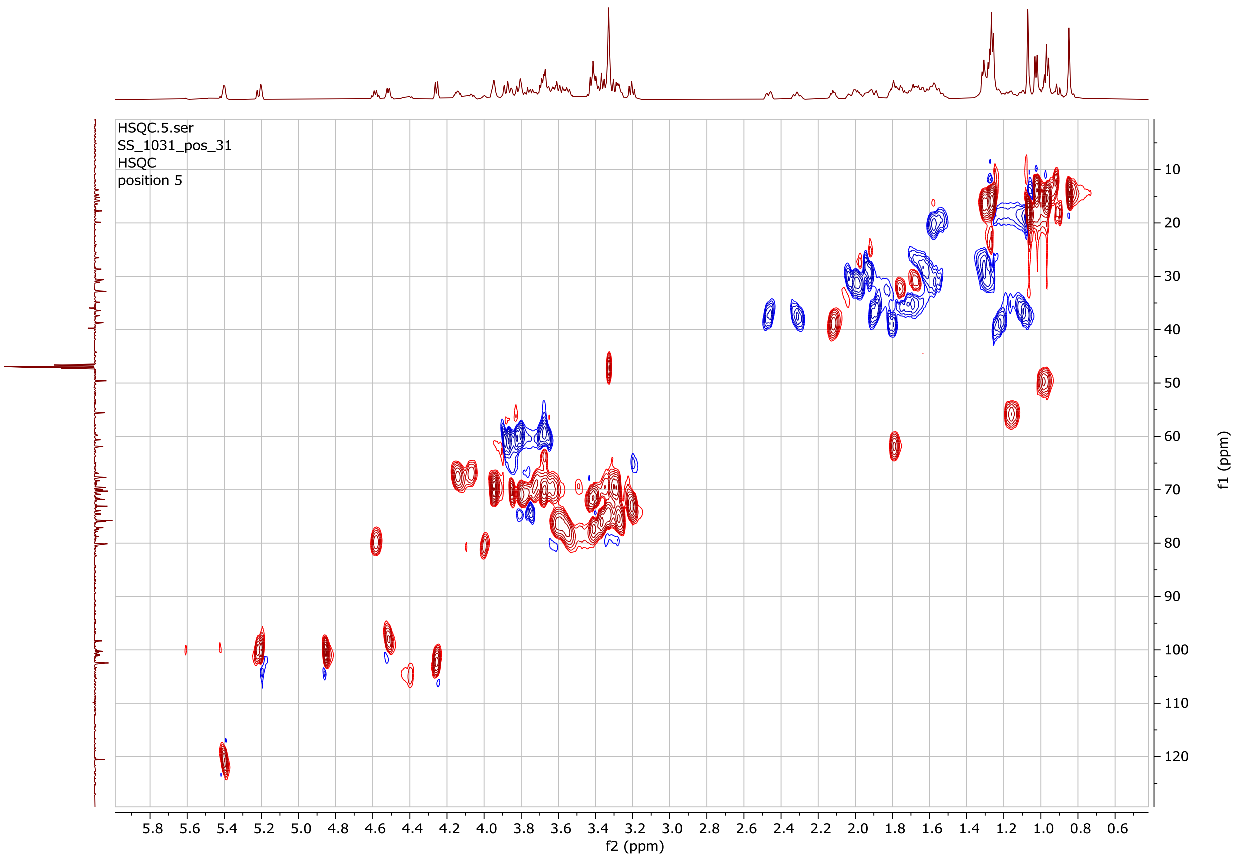


**Figure S17. gHSQCAD NMR spectrum for the intact molecule of the switchgrass saponin SS1031.**

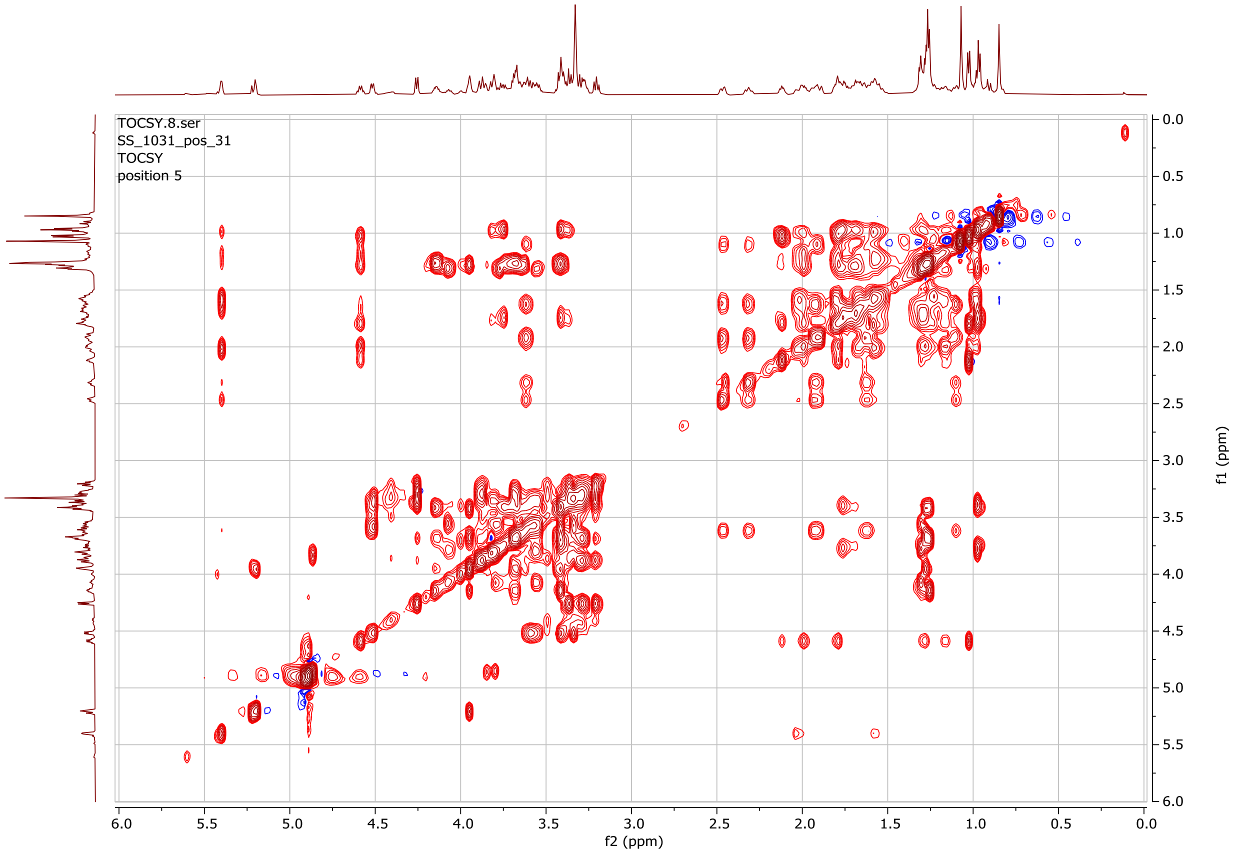


**Figure S18. gTOCSY NMR spectrum for the intact molecule of the switchgrass saponin SS1031.**

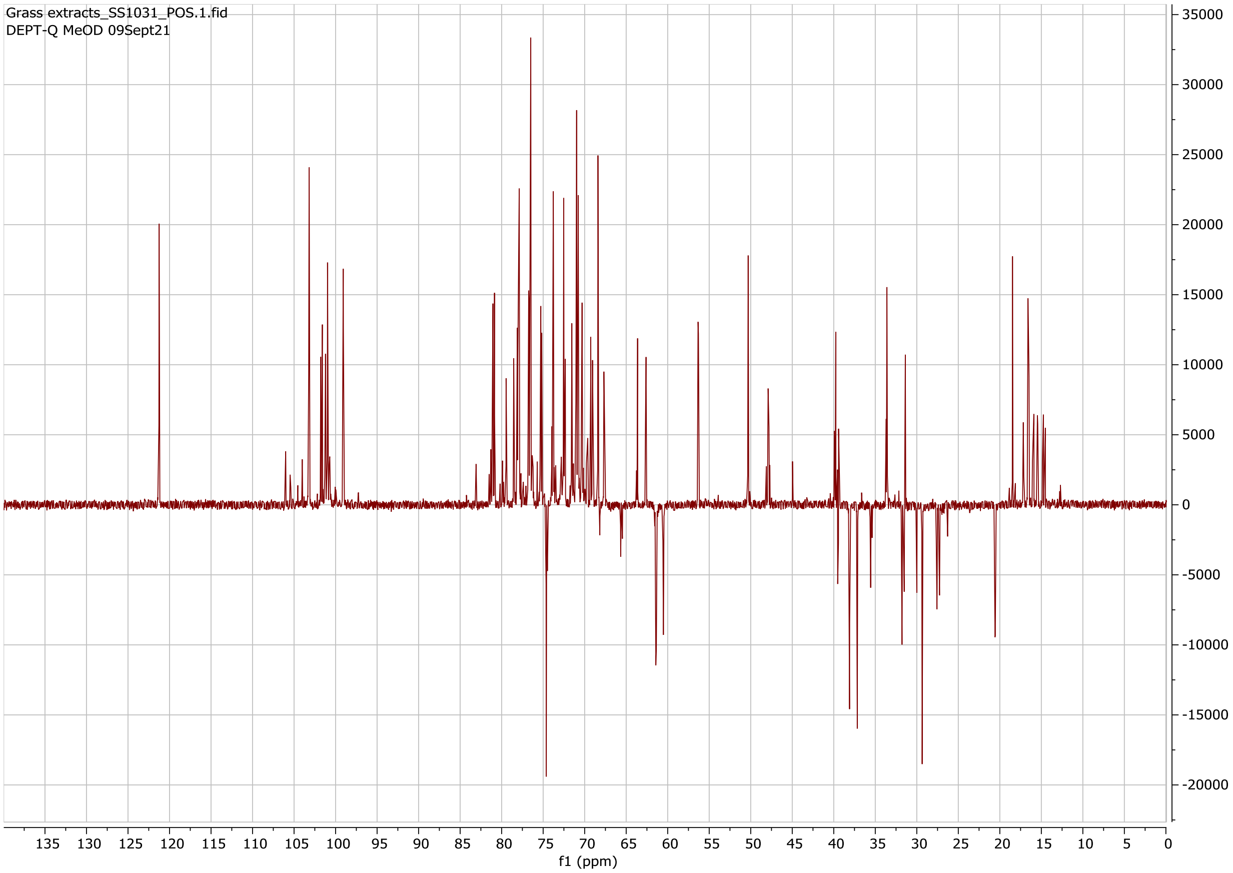


**Figure S19. DEPT-Q ($^{13}C$) NMR spectrum for the intact molecule of the switchgrass saponin SS1031.**

**Table S16. Chemical shift assignments of the aglycone of the switchgrass saponin SS1032. Chemical formula: $C_{27}H_{42}O_3$**

| Position | 1H δ (ppm) | 13C δ (ppm) | Annotation |
|---|---|---|---|
| 1 | 1.89/1.09 | 36.71 | CH2 |
| 2 | 1.94/1.64 | 28.44 | CH2 |
| 3[a] | 3.62 | 70.14 | CH |
| 4 | 2.46/2.31 | 37.13 | CH2 |
| 5[b] | | 139.92 | olefinic =C |
| 6 | 5.4 | 120.71 | olefinic =CH |
| 7 | 2.04/1.57 | 30.28 | CH2 |
| 8 | 1.7 | 27.28 | CH |
| 9 | 0.99 | 49.57 | CH |
| 10 | | 35.99 | quaternary C |
| 11 | 1.57/1.55 | 20.57 | CH2 |
| 12 | 1.78/1.22 | 38.85 | CH2 |
| 13 | | 39.42 | quaternary C |
| 14 | 1.16 | 56.0 | CH |
| 15 | 1.91/1.3 | 29.43 | CH2 |
| 16 | 4.42 | 80.43 | CH |
| 17 | 1.76 | 61.57 | CH |
| 18 | 0.83 | 17.85 | CH3 |
| 19 | 1.06 | 17.85 | CH3 |
| 20 | 1.93 | 41.0 | CH |
| 21 | 0.99 | 13.14 | CH3 |
| 22 | | 108.42 | quaternary C |
| 23 | 2/1.72 | 30.97 | CH2 |
| 24 | 1.64/1.44 | 28.44 | CH2 |
| 25 | 1.67 | 30.71 | CH |
| 26[c] | 3.45/3.34 | 65.43 | CH2 |
| 27 | 0.81 | 15.28 | CH3 |

[a] The C-3 carbon connects to a sugar anomeric proton, 5.2 *ppm*, through HMBC.
[b] The C-5 carbon chemical shift, 139.92 *ppm*, was determined by HMBC.
[c] One of the two C-26 protons, 3.45 *ppm*, correlates with the C-22 carbon, 108.42 *ppm*, through HMBC (suggesting a pyranosidic ring).

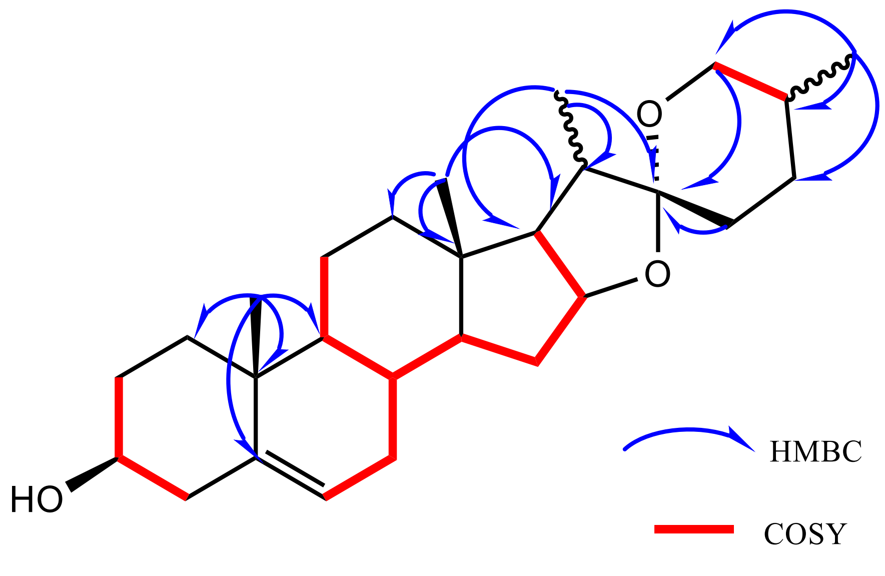


**Figure S20. The key COSY (red, bold) and HMBC (blue arrow) correlations for the aglycone of the switchgrass saponin SS1032.**

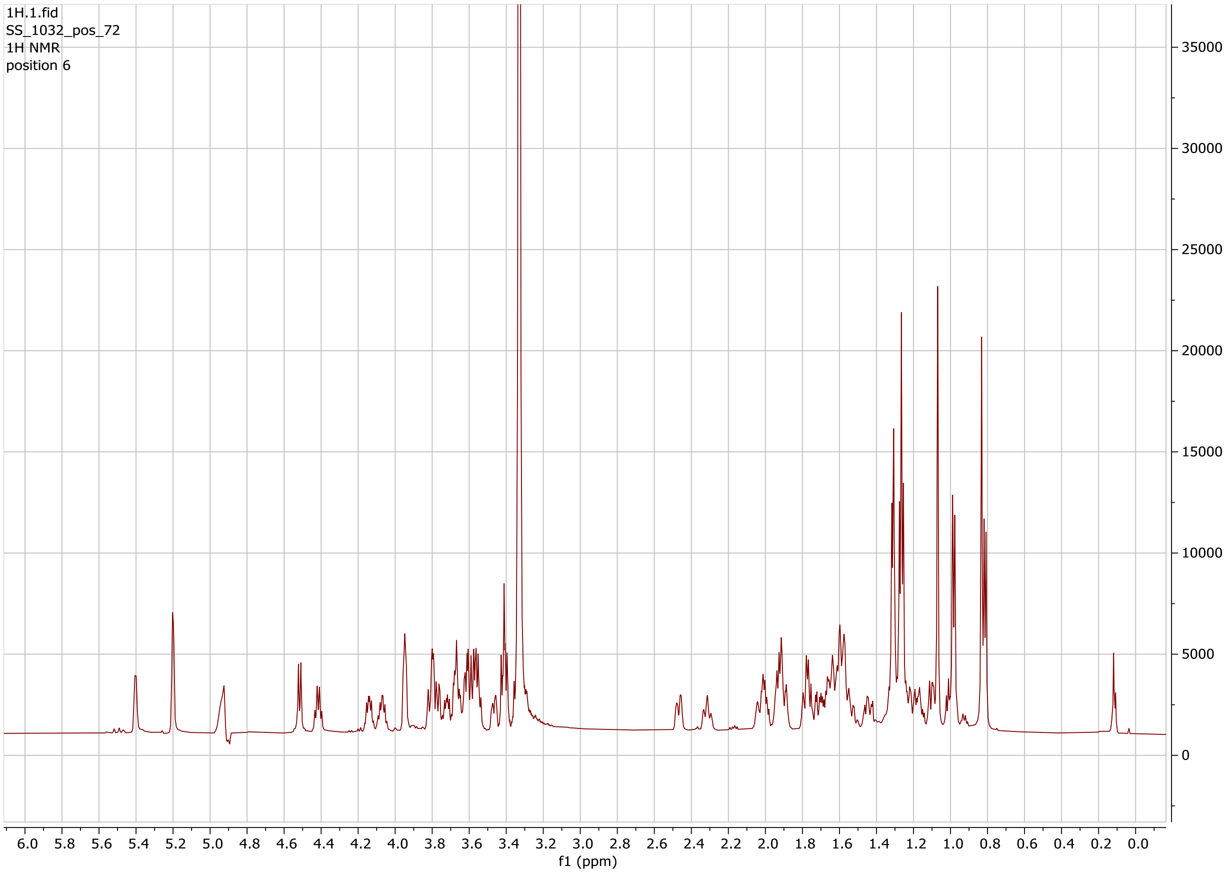


**Figure S21. $^{1}$H NMR spectrum for the intact molecule of the switchgrass saponin SS1032.**

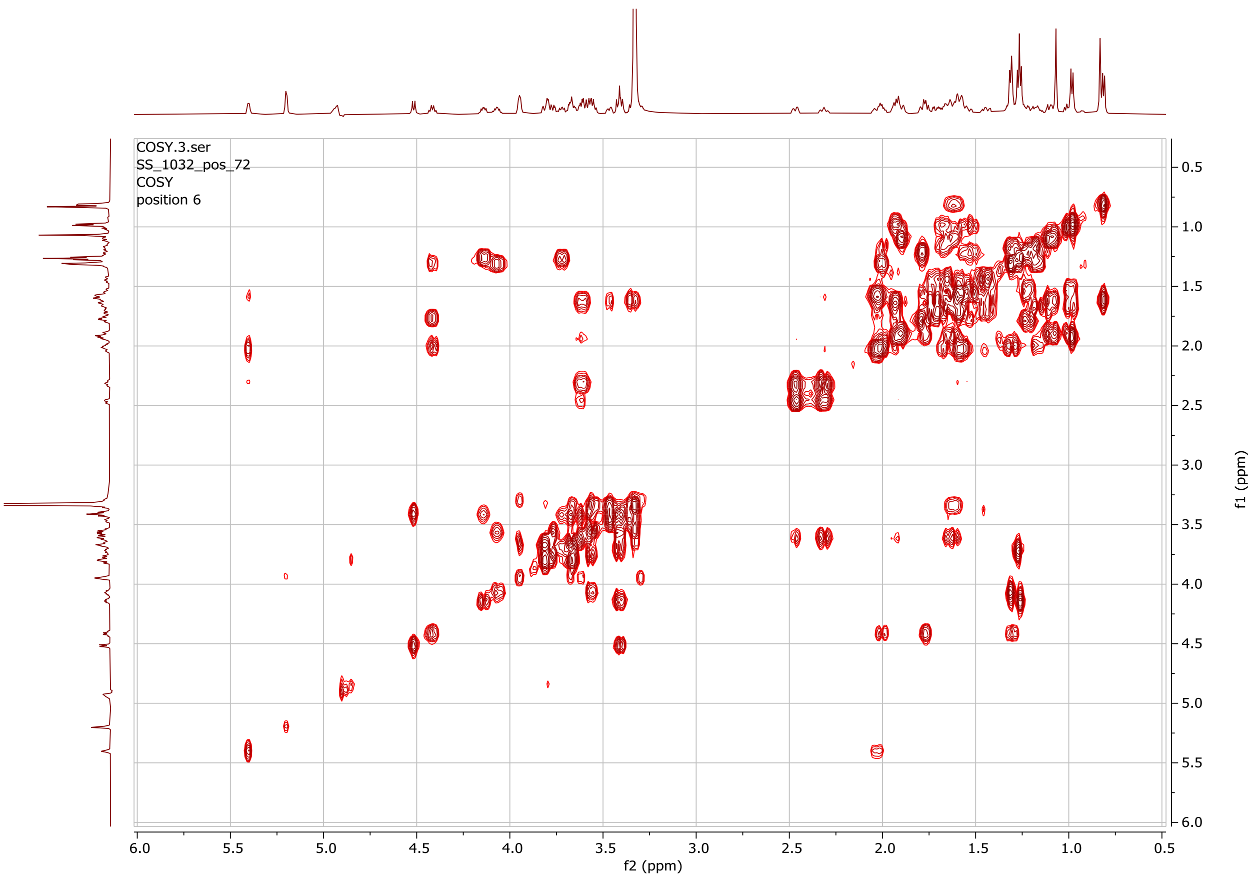


**Figure S22. gCOSY $^{1}H$-$^{1}H$ NMR spectrum for the intact molecule of the switchgrass saponin SS1032.**

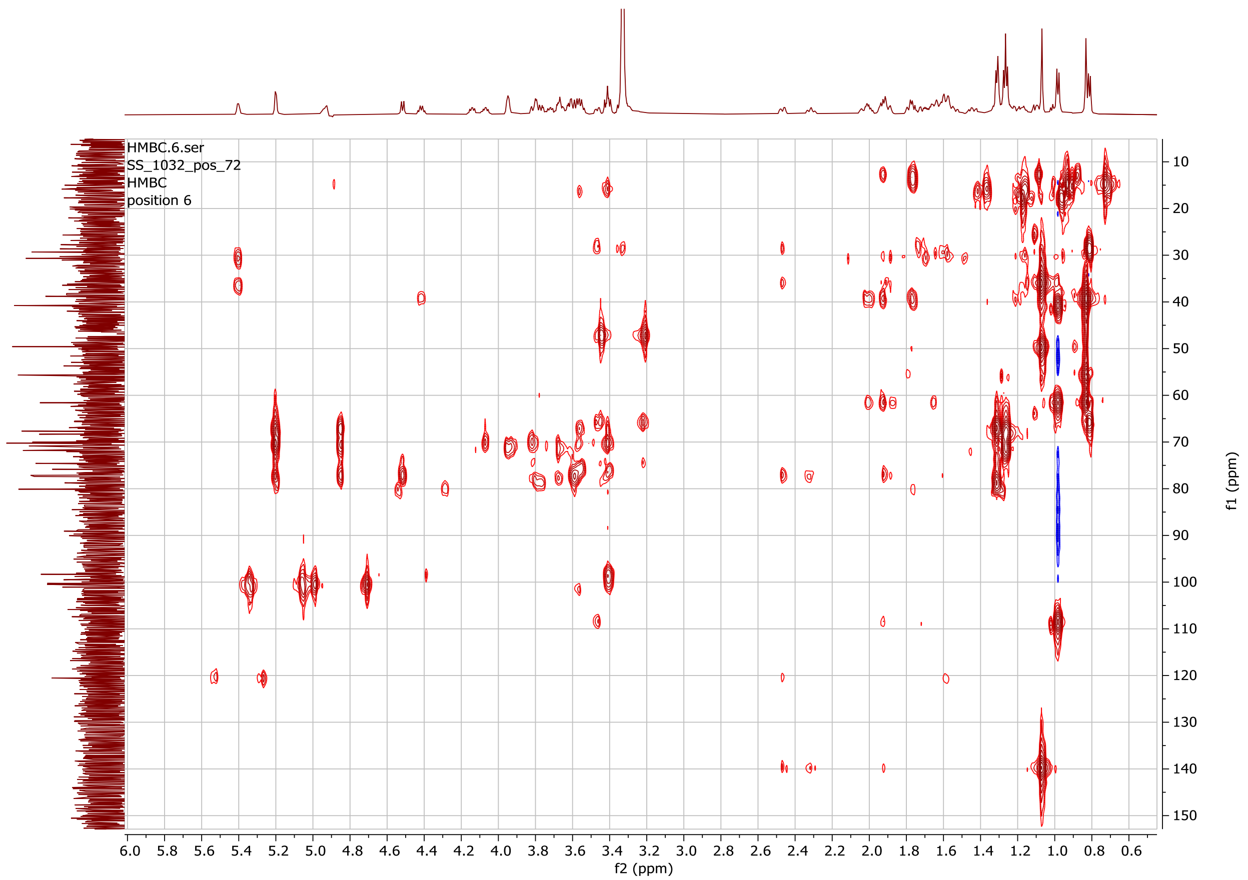


**Figure S23. gHMBCAD NMR spectrum for the intact molecule of the switchgrass saponin SS1032.**

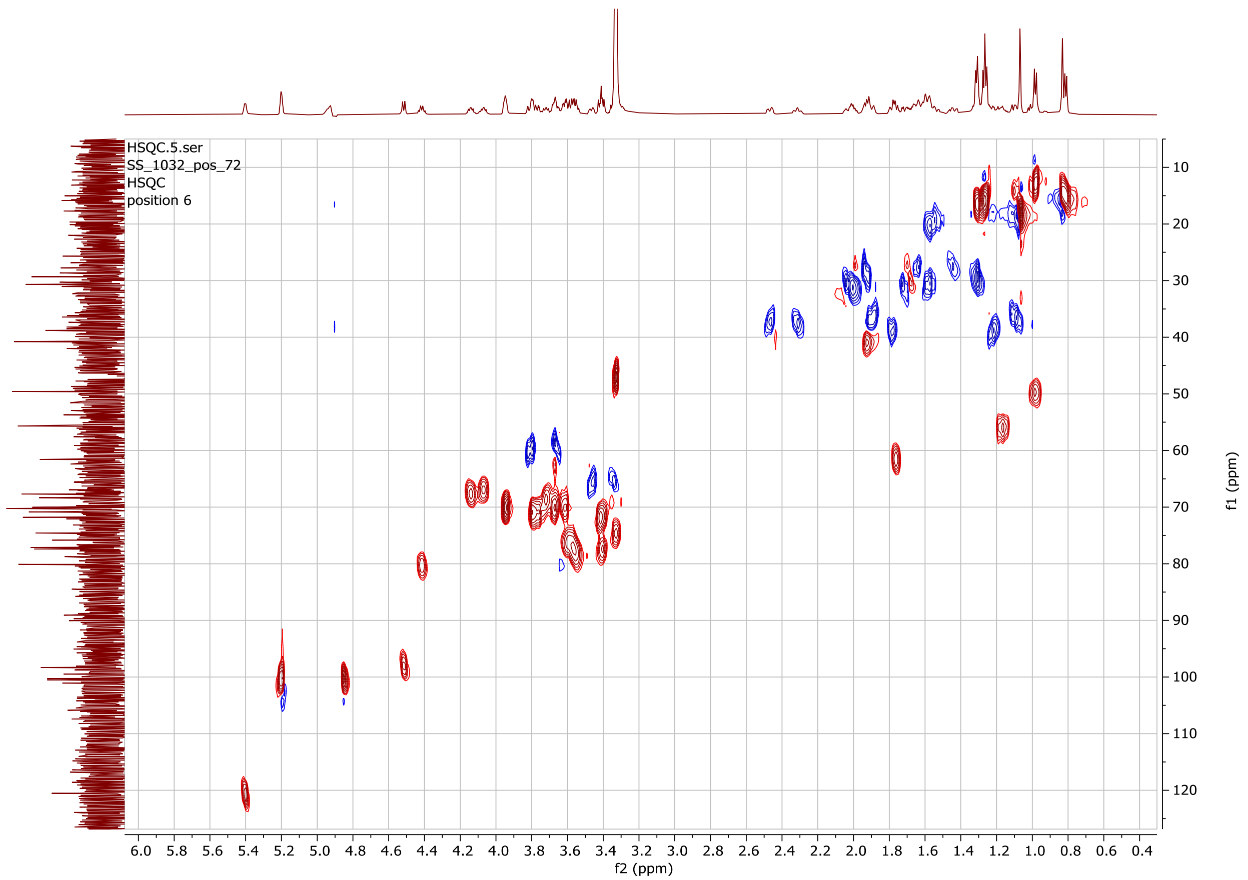


**Figure S24. gHSQCAD NMR spectrum for the intact molecule of the switchgrass saponin SS1032.**

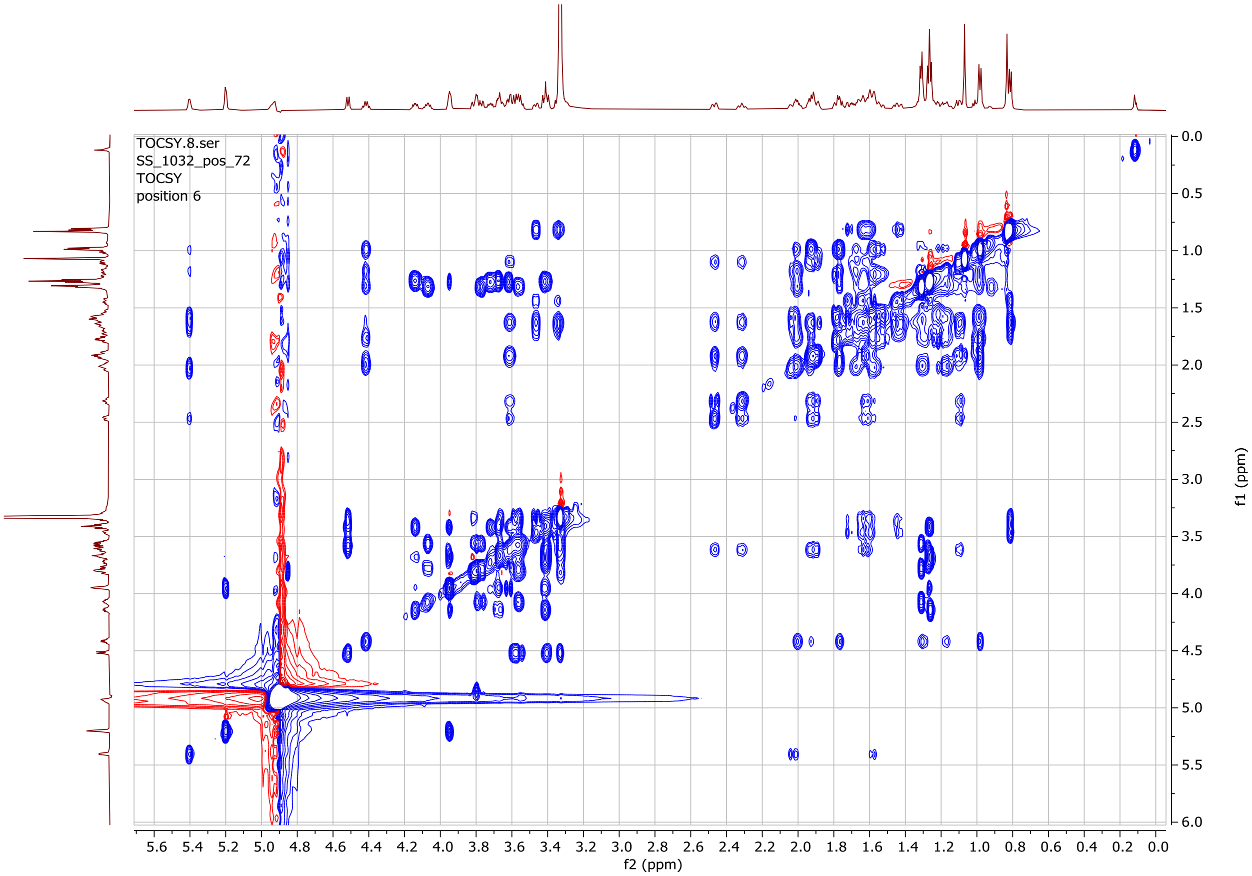


**Figure S25. gTOCSY NMR spectrum for the intact molecule of the switchgrass saponin SS1032.**

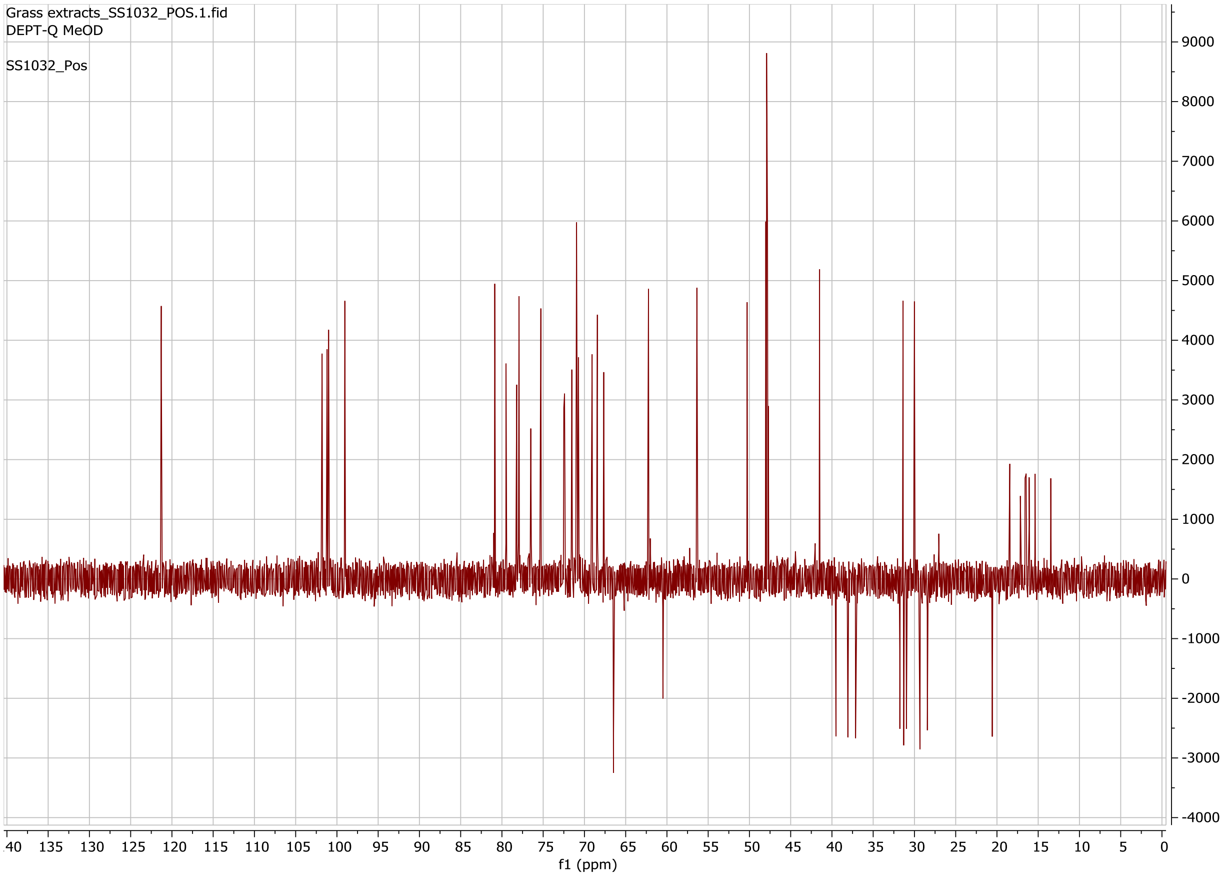


**Figure S26. DEPT-Q ($^{13}C$) NMR spectrum for the intact molecule of the switchgrass saponin SS1032.**

**Table S17. Chemical shift assignments of the aglycone of the switchgrass saponin SS1244. Chemical formular: $C_{27}H_{44}O_5$**

| Position | 1H δ (ppm) | 13C δ (ppm) | Annotation |
|---|---|---|---|
| 1 | 1.9/1.1 | 36.28 | CH2 |
| 2 | 1.94/1.64 | 28.14 | CH2 |
| 3[a] | 3.63 | 70.14 | CH |
| 4 | 2.46/2.31 | 37.57 | CH2 |
| 5[b] | | 139.71 | olefinic =C |
| 6 | 5.4 | 120.71 | olefinic =CH |
| 7 | 2.02/1.58 | 31.52 | CH2 |
| 8 | 1.65 | 31.57 | CH |
| 9 | 0.96 | 49.57 | CH |
| 10 | | 35.99 | quaternary C |
| 11 | 1.64/1.56 | 20.31 | CH2 |
| 12 | 1.93/1.3 | 38.14 | CH2 |
| 13 | | 44.14 | quaternary C |
| 14 | 1.71 | 51.71 | CH |
| 15 | 2.04/1.27 | 30.28 | CH2 |
| 16 | 4.18 | 88.57 | CH |
| 17 | | 89.57 | quaternary C |
| 18 | 0.87 | 15.28 | CH3 |
| 19 | 1.06 | 17.85 | CH3 |
| 20 | 2.27 | 41.43 | CH |
| 21 | 0.95 | 8.0 | CH3 |
| 22 | | 110.14 | quaternary C |
| 23 | 2.05/1.71 | 35.0 | CH2 |
| 24 | 1.44/1.71 | 26.0 | CH2 |
| 25 | 1.76 | 32.43 | CH |
| 26[c] | 3.76/3.4 | 74.0 | CH2 |
| 27 | 0.96 | 15.28 | CH3 |

[a] The C-3 carbon connects to a sugar anomeric proton, 5.61 *ppm*, through HMBC.
[b] The C-5 carbon chemical shift, 139.71 *ppm*, was determined by HMBC.
[c] The C-26 carbon connects to a sugar anomeric proton, 4.25 *ppm*, through HMBC.

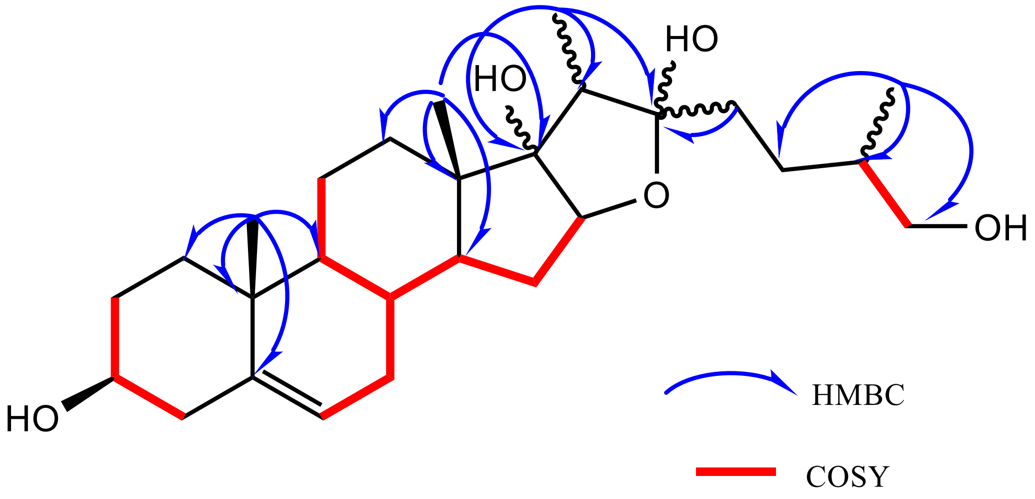


**Figure S27. The key COSY (red, bold) and HMBC (blue arrow) correlations for the aglycone of the switchgrass saponin SS1244.**

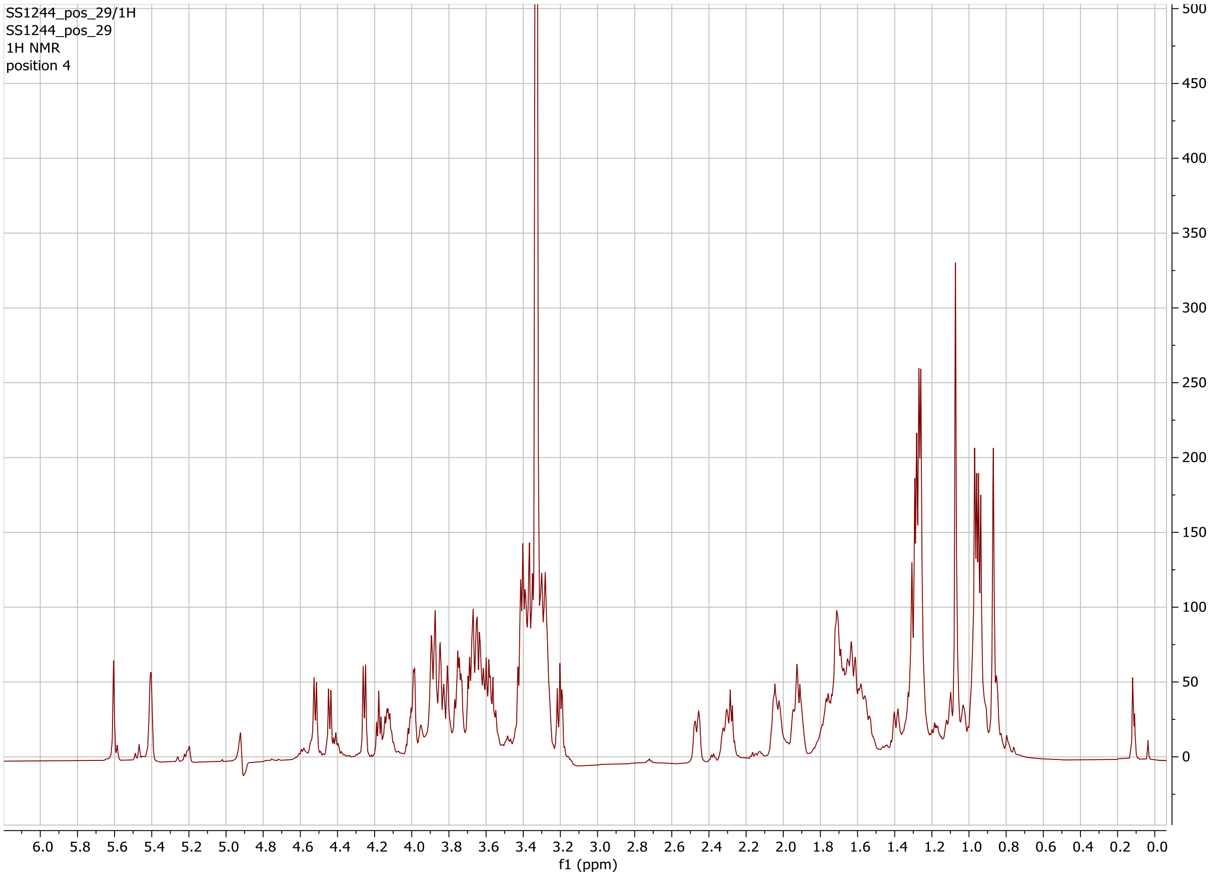


**Figure S28. ¹H NMR spectrum for the intact molecule of the switchgrass saponin SS1244.**

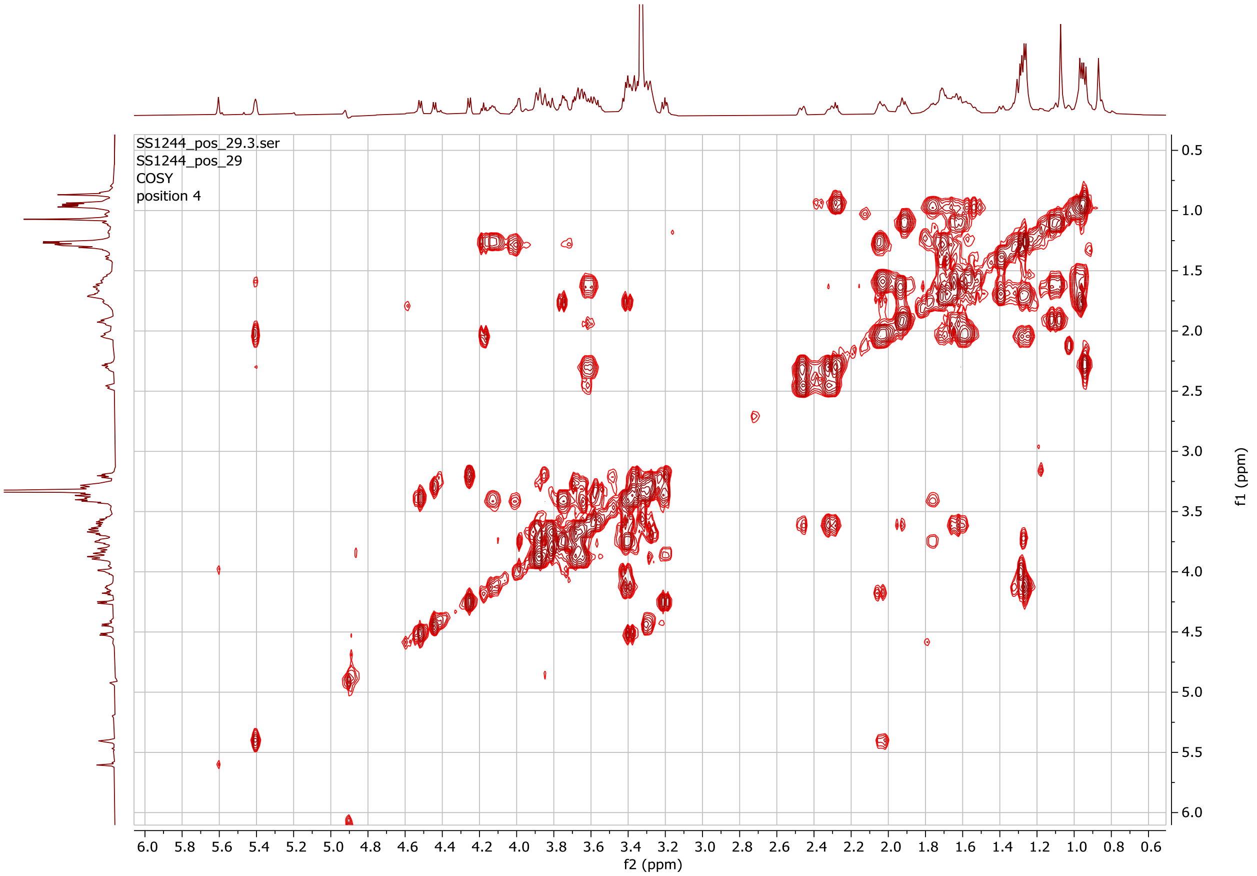


**Figure S29. gCOSY $^{1}$H-$^{1}$H NMR spectrum for the intact molecule of the switchgrass saponin SS1244.**

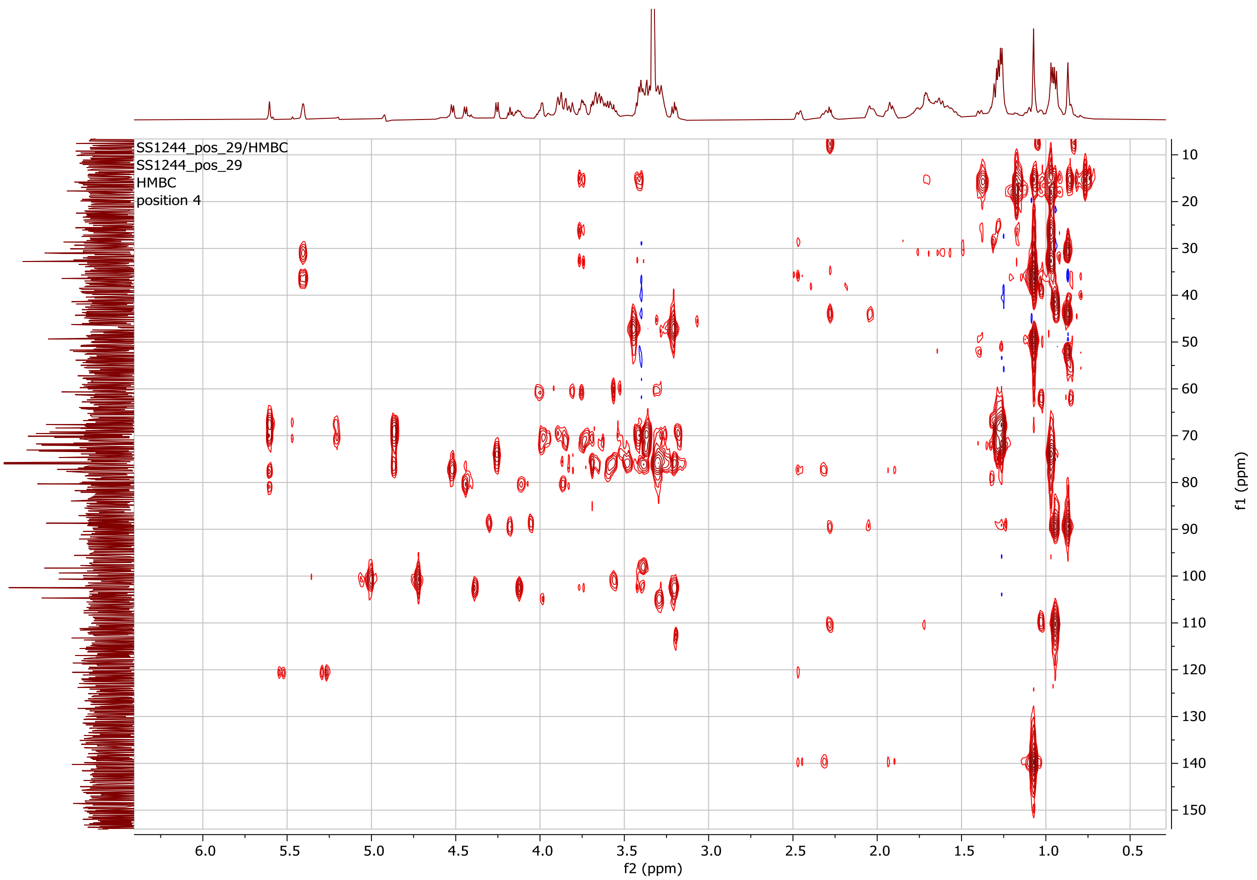


**Figure S30. gHMBCAD NMR spectrum for the intact molecule of the switchgrass saponin SS1244.**

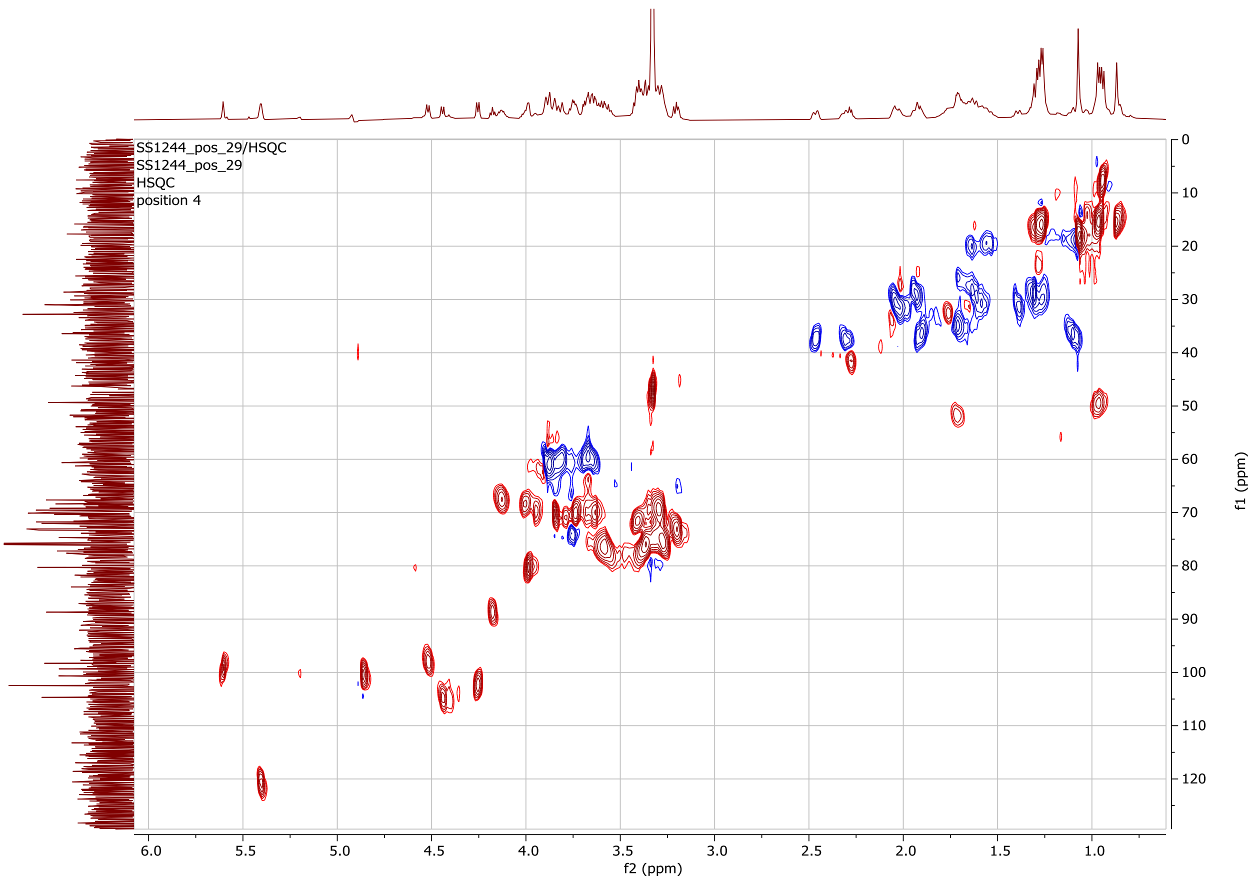


**Figure S31. gHSQCAD NMR spectrum for the intact molecule of the switchgrass saponin SS1244.**

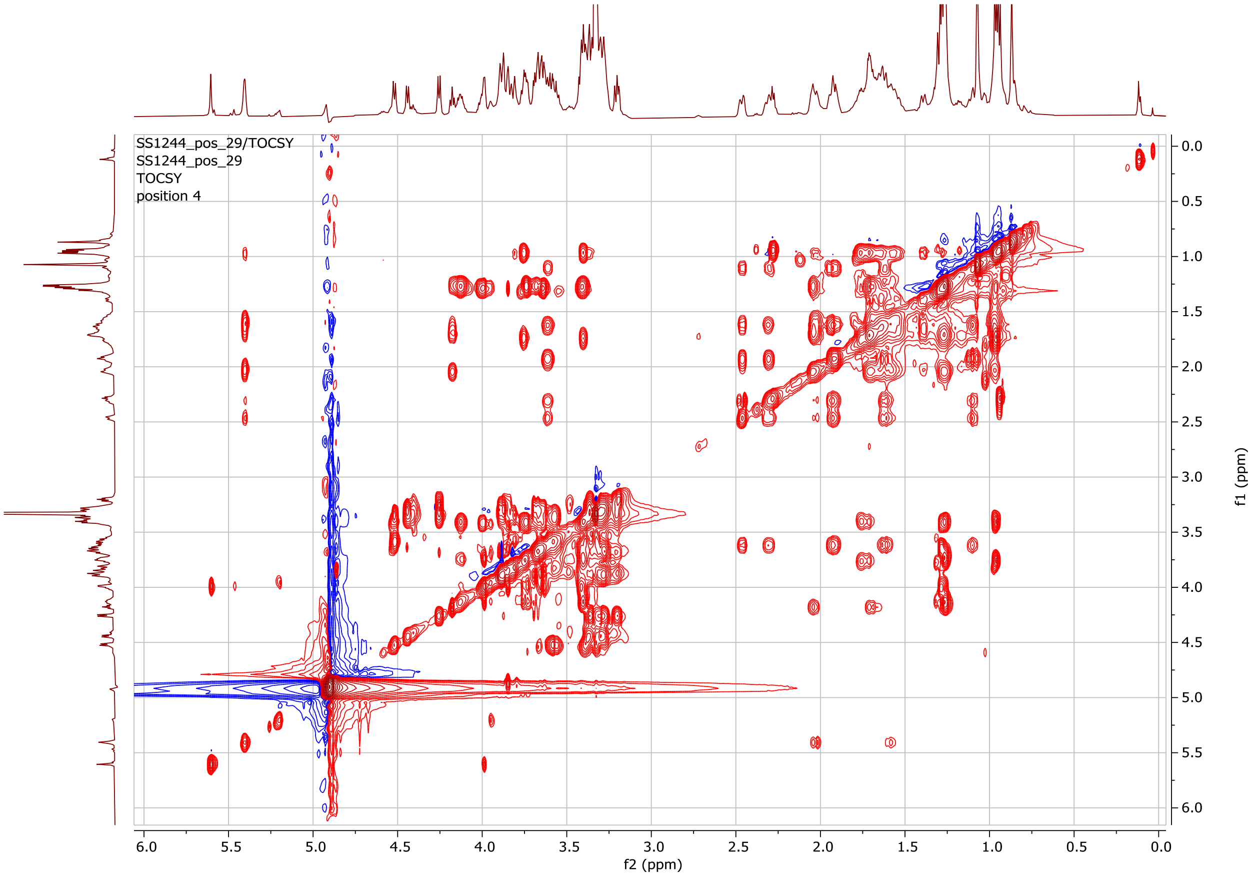


**Figure S32. gTOCSY NMR spectrum for the intact molecule of the switchgrass saponin SS1244.**

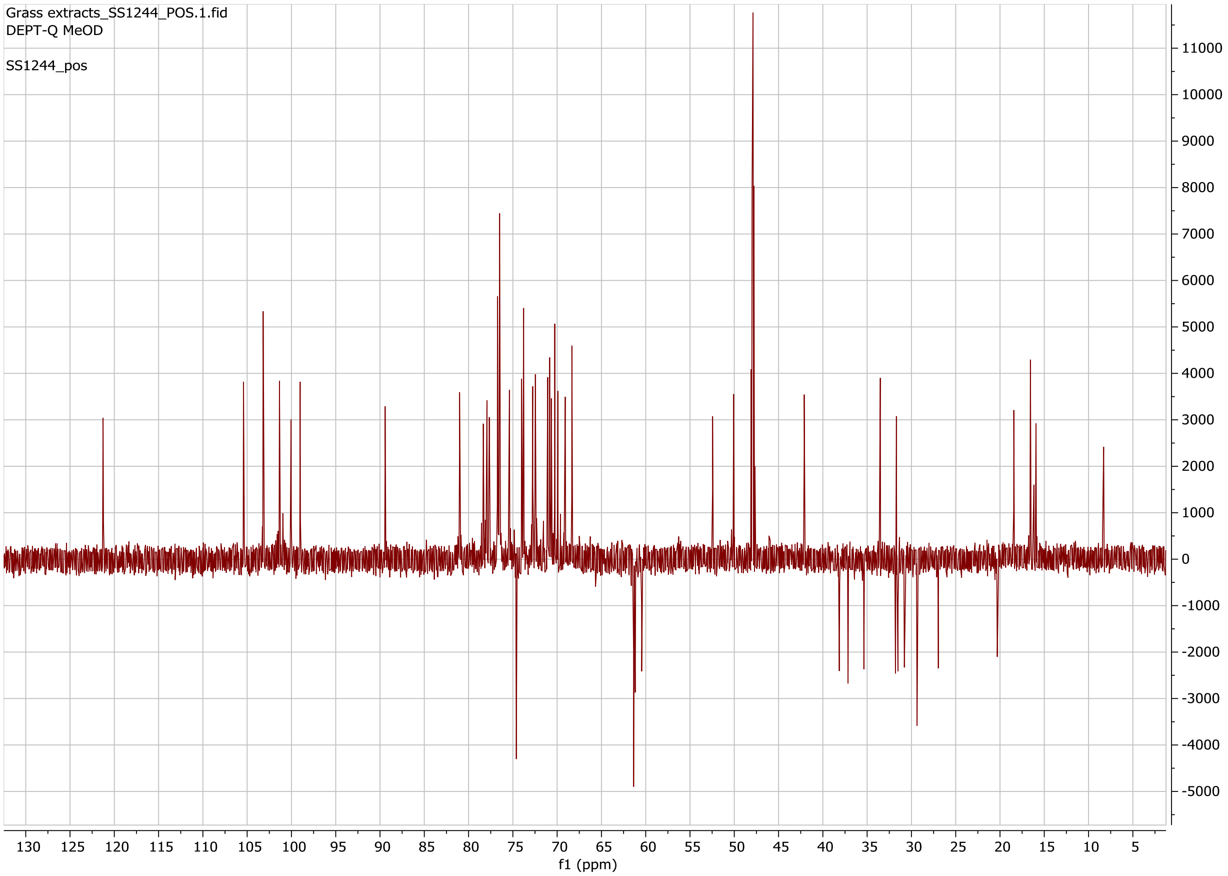


**Figure S33. DEPT-Q ($^{13}C$) NMR spectrum for the intact molecule of the switchgrass saponin SS1244.**

**Table S18. Chemical shift assignments of the aglycone of the switchgrass saponin SS1064. Chemical formular: $C_{27}H_{42}O_4$**

| Position | 1H δ (ppm) | 13C δ (ppm) | Annotation |
|---|---|---|---|
| 1 | 1.9/1.1 | 36.28 | CH2 |
| 2 | 1.95/1.64 | 27.28 | CH2 |
| 3[a] | 3.63 | 70.14 | CH |
| 4 | 2.46/2.31 | 37.17 | CH2 |
| 5[b] | | 139.71 | olefinic =C |
| 6 | 5.41 | 120.28 | olefinic =CH |
| 7 | 2.05/1.59 | 30.71 | CH2 |
| 8 | 1.64 | 27.28 | CH |
| 9 | 0.96 | 49.57 | CH |
| 10 | | 35.99 | quaternary C |
| 11 | 1.62/1.52 | 20.29 | CH2 |
| 12 | 1.93/1.3 | 38.14 | CH2 |
| 13 | | 43.71 | quaternary C |
| 14 | 1.74 | 51.71 | CH |
| 15 | 2.03/1.36 | 31.57 | CH2 |
| 16 | 4.03 | 88.57 | CH |
| 17 | | 89.14 | quaternary C |
| 18 | 0.85 | 15.28 | CH3 |
| 19 | 1.06 | 17.85 | CH3 |
| 20 | 2.11 | 43.57 | CH |
| 21 | 0.91 | 7.14 | CH3 |
| 22 | | 108.85 | quaternary C |
| 23 | 2.11/1.72 | 30.71 | CH2 |
| 24 | 1.94/1.43 | 28.14 | CH2 |
| 25 | 1.64 | 31.14 | CH |
| 26[c] | 3.49/3.35 | 66.3 | CH2 |
| 27 | 0.82 | 15.28 | CH3 |

[a] The C-3 carbon connects to a sugar anomeric proton, 5.61 *ppm*, through HMBC.
[b] The C-5 carbon chemical shift, 139.71 *ppm*, was determined by HMBC.
[c] One of the two C-26 protons, 3.49 *ppm*, correlates with the C-22 carbon, 108.85 *ppm*, through HMBC (suggesting a pyranosidic ring).

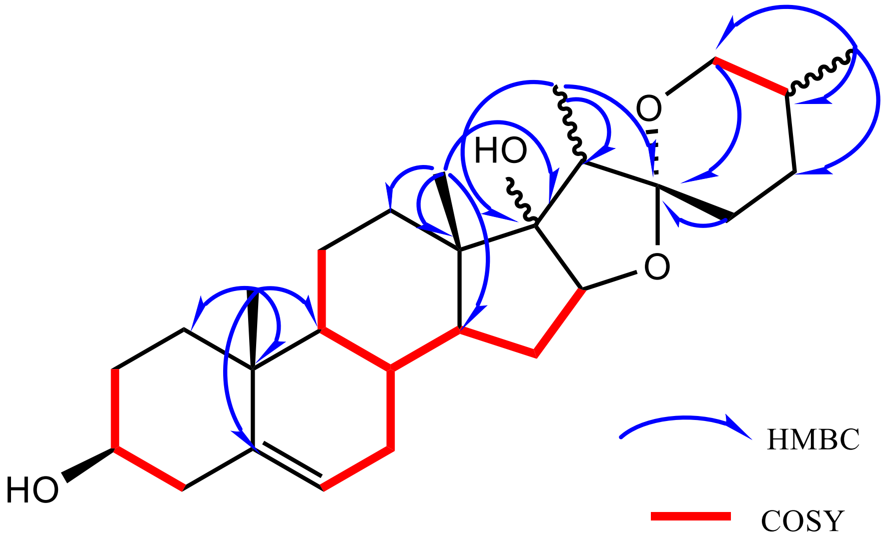


**Figure S34. The key COSY (red, bold) and HMBC (blue arrow) correlations for the aglycone of the switchgrass saponin SS1064.**

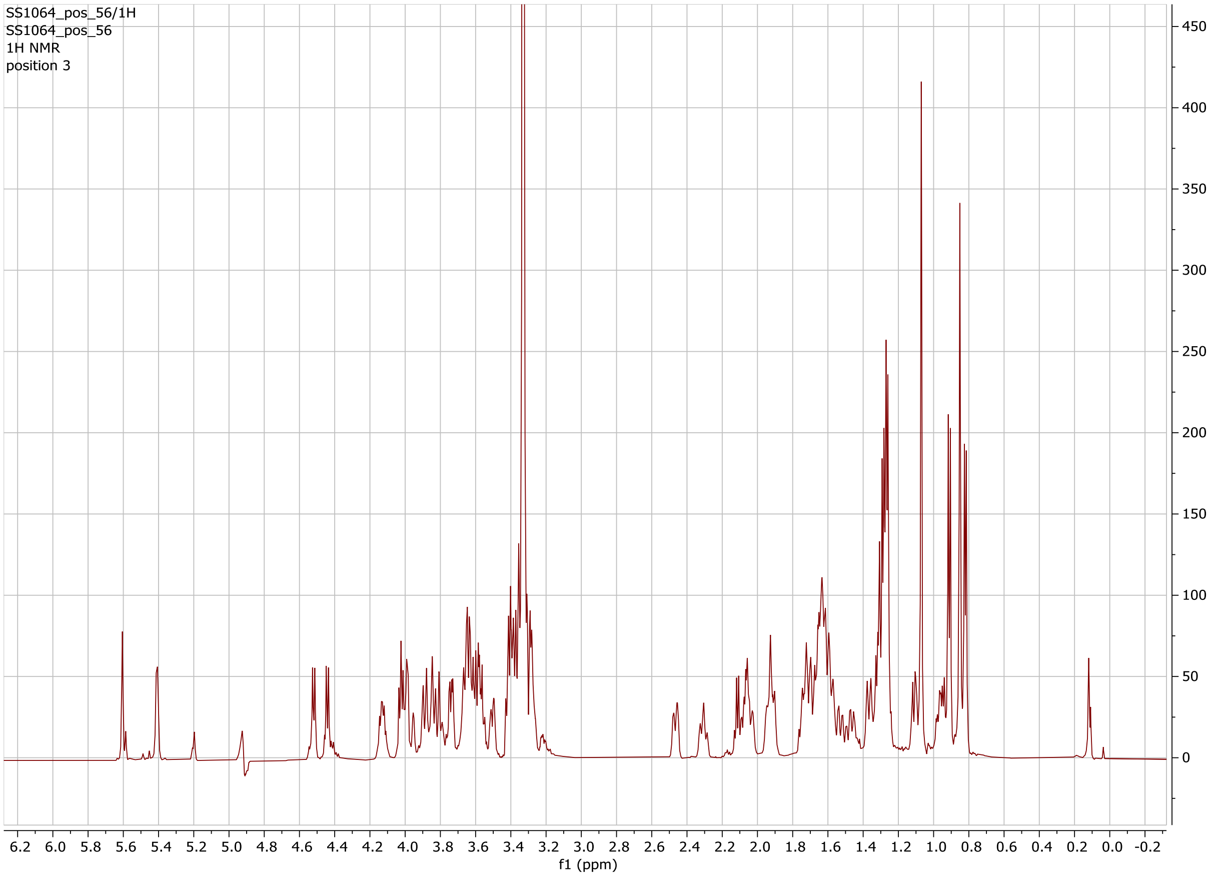


**Figure S35.** **$^1$H NMR spectrum for the intact molecule of the switchgrass saponin SS1064.**

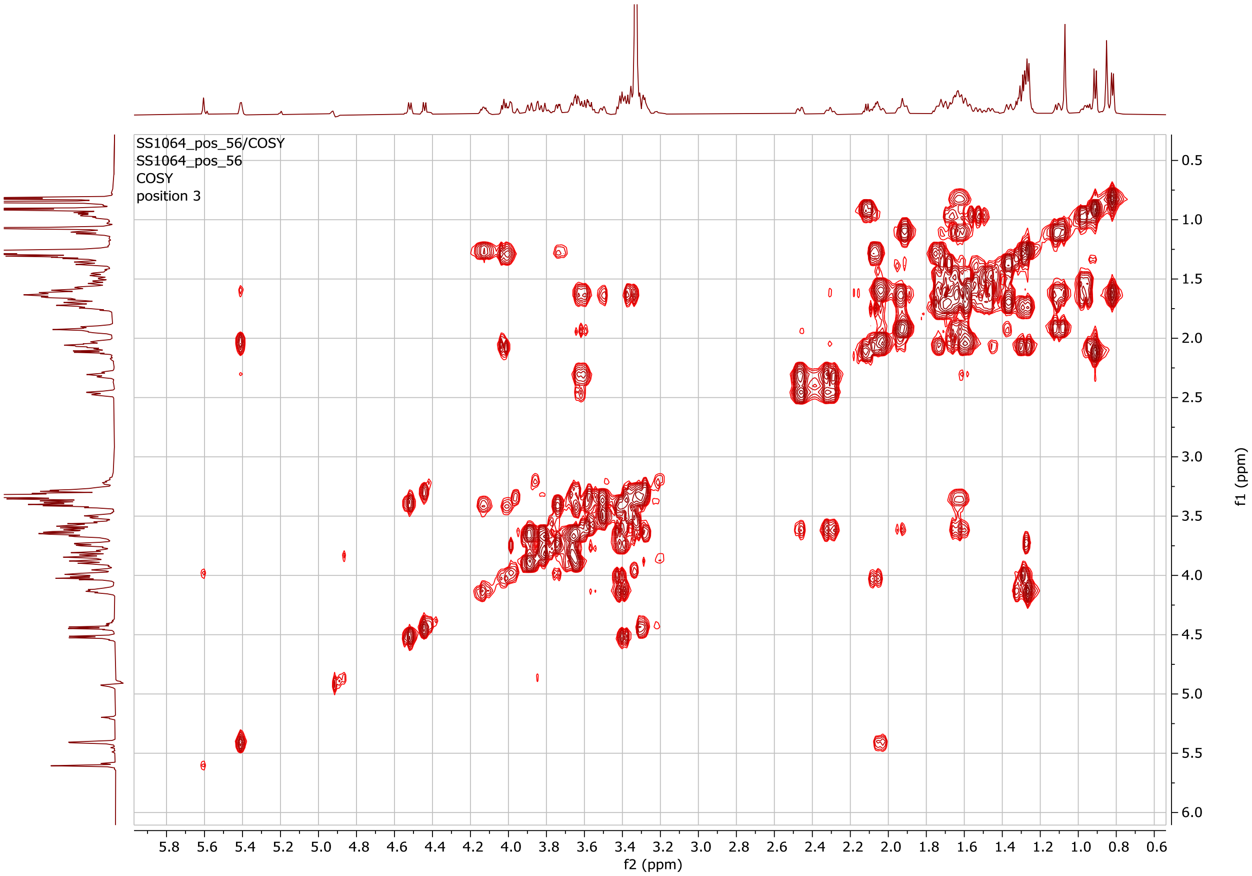


**Figure S36. gCOSY NMR spectrum for the intact molecule of the switchgrass saponin SS1064.**

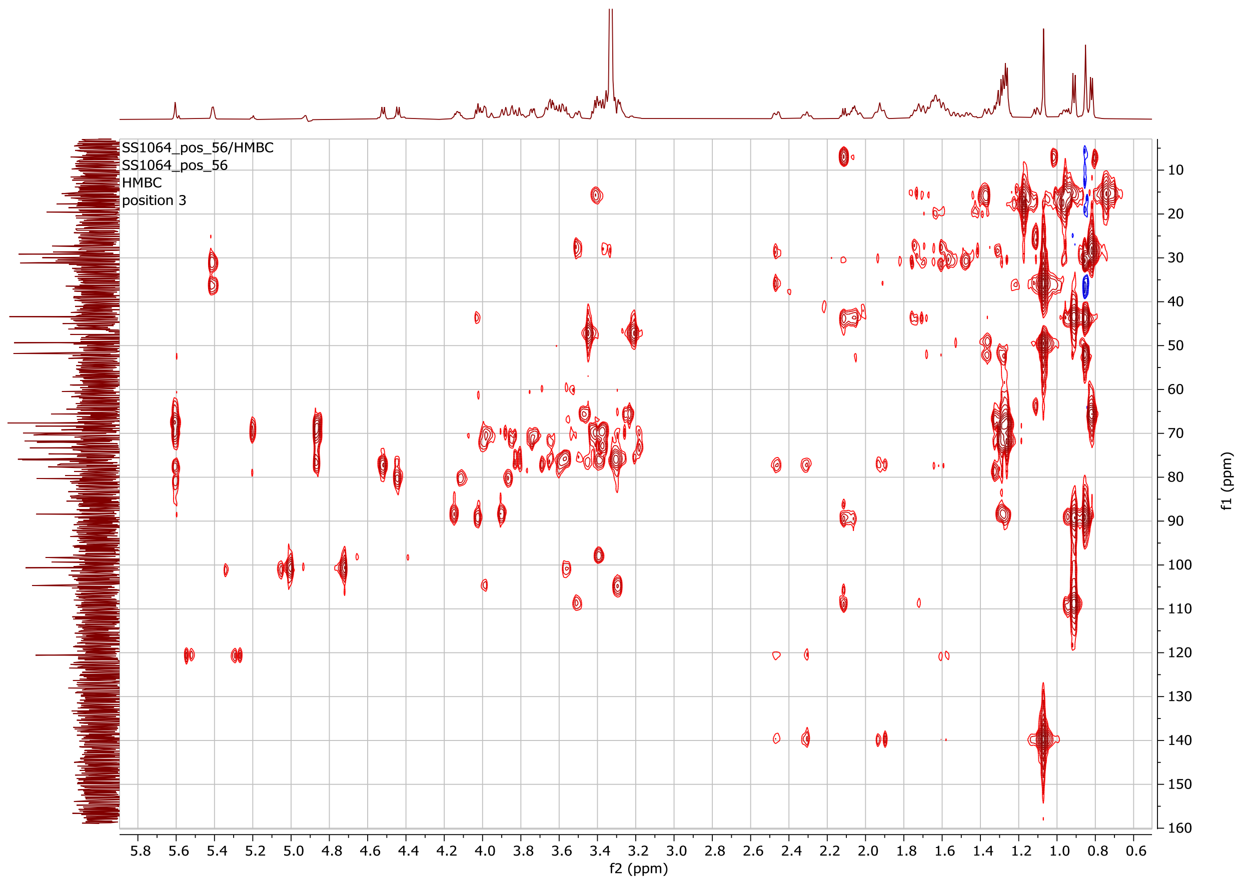


**Figure S37. gHMBCAD NMR spectrum for the intact molecule of the switchgrass saponin SS1064.**

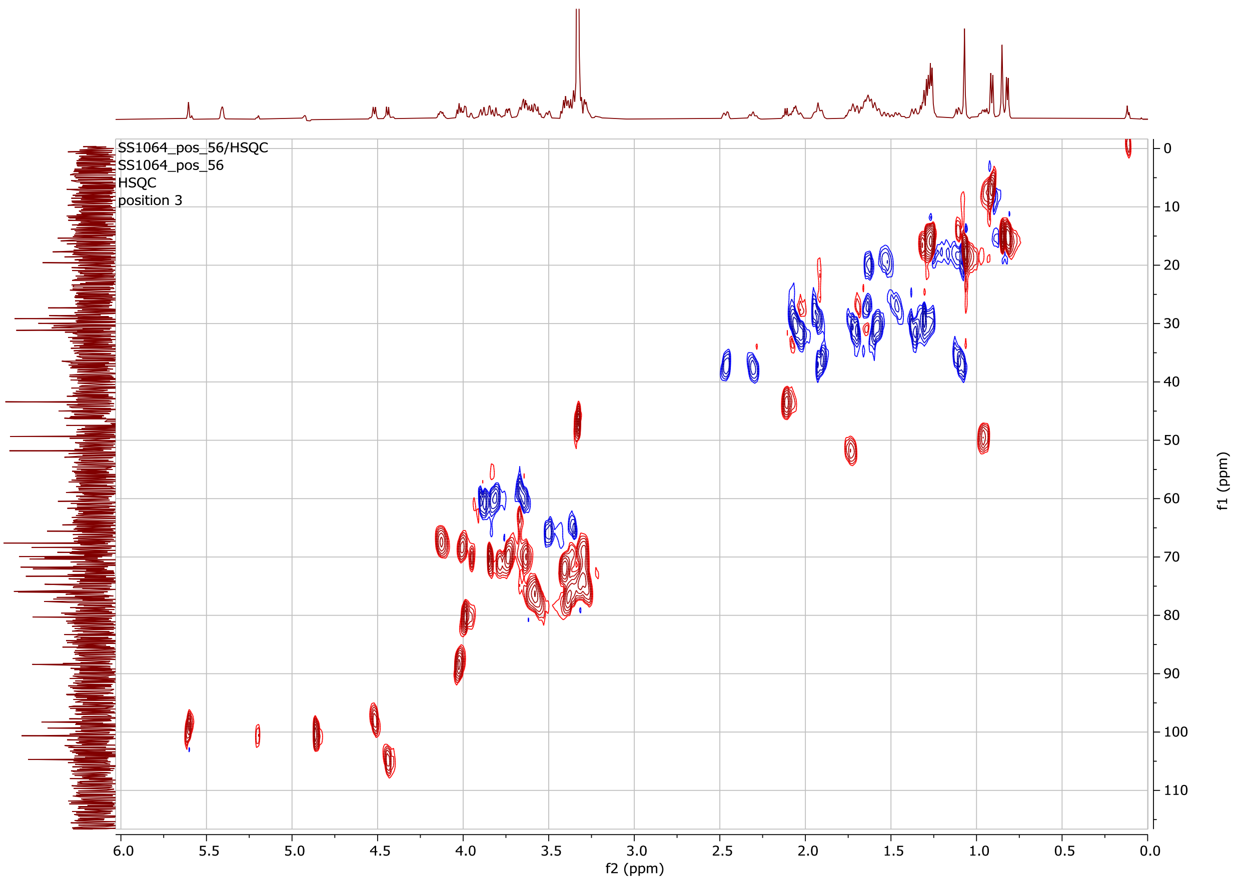


**Figure S38. gHSQCAD NMR spectrum for the intact molecule of the switchgrass saponin SS1064.**

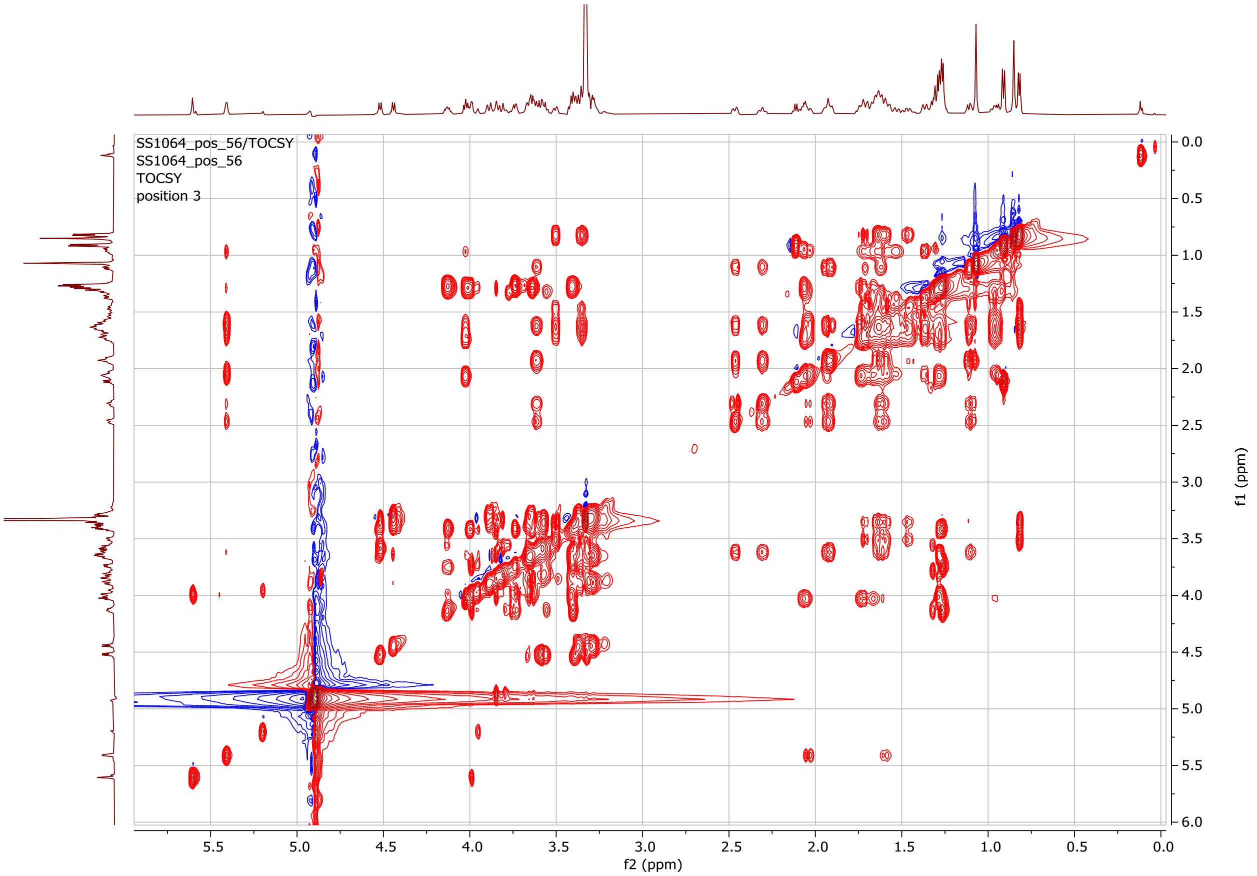


**Figure S39. gTOCSY NMR spectrum for the intact molecule of the switchgrass saponin SS1064.**

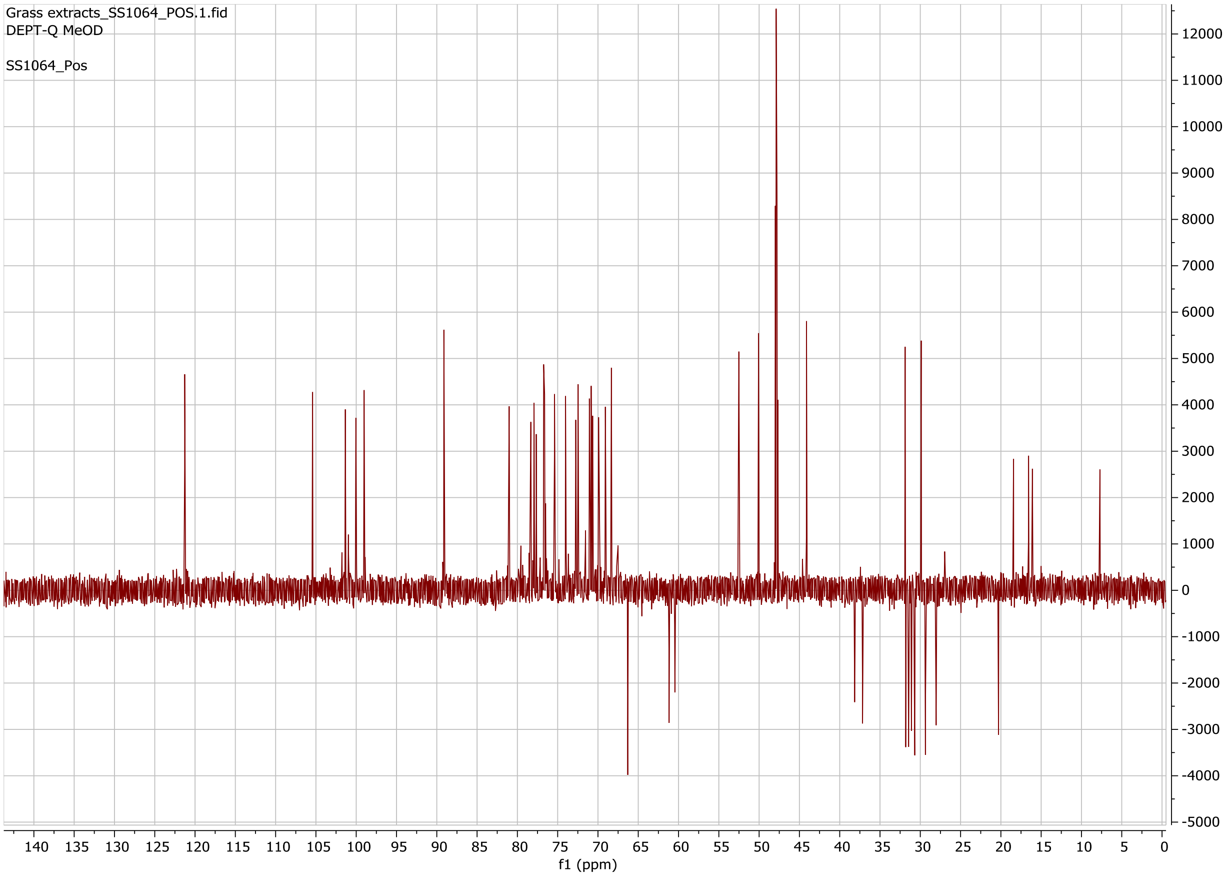


**Figure S40. DEPT-Q ($^{13}C$) NMR spectrum for the intact molecule of the switchgrass saponin SS1064.**

**Table S19. Chemical shift assignments of the aglycone of the switchgrass saponin SS1050. Chemical formular: $C_{27}H_{44}O_3$**

| Position | 1H δ (ppm) | 13C δ (ppm) | Annotation |
|---|---|---|---|
| 1 | 1.89/1.1 | 37.19 | CH2 |
| 2 | 1.94/1.64 | 28.14 | CH2 |
| 3[a] | 3.63 | 70.14 | CH |
| 4 | 2.45/2.31 | 38.08 | CH2 |
| 5[b] | | 139.28 | olefinic =C |
| 6 | 5.4 | 120.71 | olefinic =CH |
| 7 | 2/1.58 | 29.35 | CH2 |
| 8 | 1.5 | 31.14 | CH |
| 9 | 0.99 | 50 | CH |
| 10 | | 35.57 | quaternary C |
| 11 | 1.58/1.54 | 20.73 | CH2 |
| 12 | 2.03/1.3 | 38.88 | CH2 |
| 13 | | 42.42 | quaternary C |
| 14 | 1.08 | 55.57 | CH |
| 15 | 1.91/1.3 | 29.35 | CH2 |
| 16 | 1.91/1.08 | 37.14 | CH2 |
| 17 | 1.61 | 51.71 | CH |
| 18 | 0.77 | 10.14 | CH3 |
| 19 | 1.05 | 17.85 | CH3 |
| 20 | 2.61 | 48.71 | CH |
| 22[c] | | 215.57 | Ketone carbonyl C |
| 21 | 1.13 | 14.85 | CH3 |
| 23 | 2.56/1.97 | 31.57 | CH2 |
| 24 | 1.7/1.39 | 26.98 | CH2 |
| 25 | 1.76 | 32 | CH |
| 26[d] | 3.76/3.39 | 74 | CH2 |
| 27 | 0.96 | 15.28 | CH3 |

[a] The C-3 carbon connects to a sugar anomeric proton, 5.22 *ppm*, through HMBC.
[b] The C-5 carbon chemical shift, 139.28 *ppm*, was determined by HMBC.
[c] The C-22 carbon chemical shift, 215.57 *ppm*, was determined by HMBC.
[d] The C-26 carbon connects to a sugar anomeric proton, 4.26 *ppm*, through HMBC.

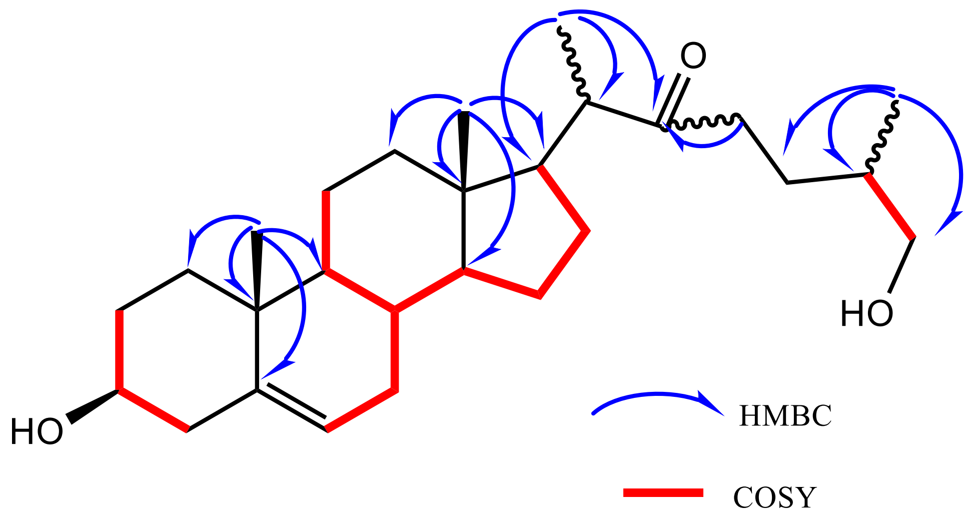


**Figure S41. The key COSY (red, bold) and HMBC (blue arrow) correlations for the aglycone of the switchgrass saponin SS1050.**

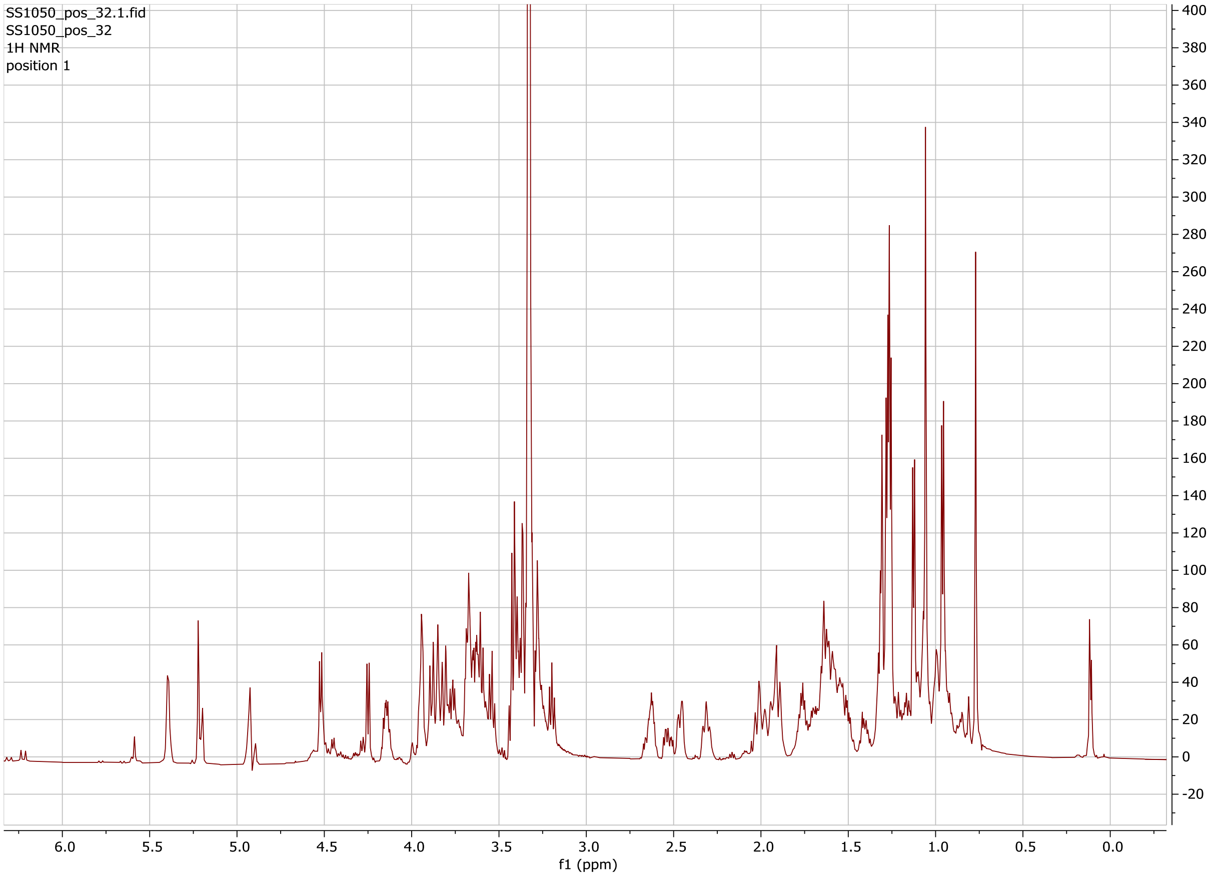


**Figure S42. $^{1}$H NMR spectrum for the intact molecule of the switchgrass saponin SS1050.**

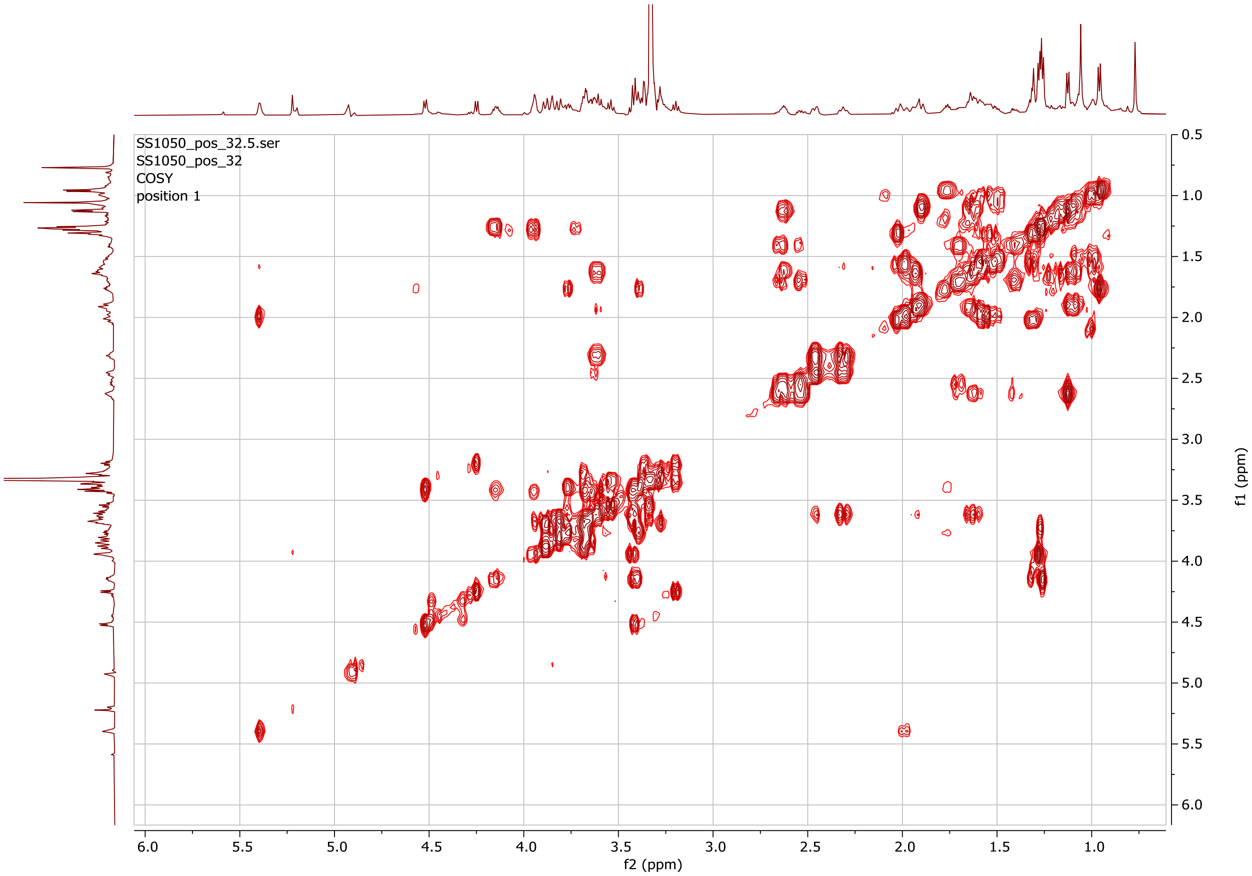


**Figure S43. gCOSY NMR spectrum for the intact molecule of the switchgrass saponin SS1050.**

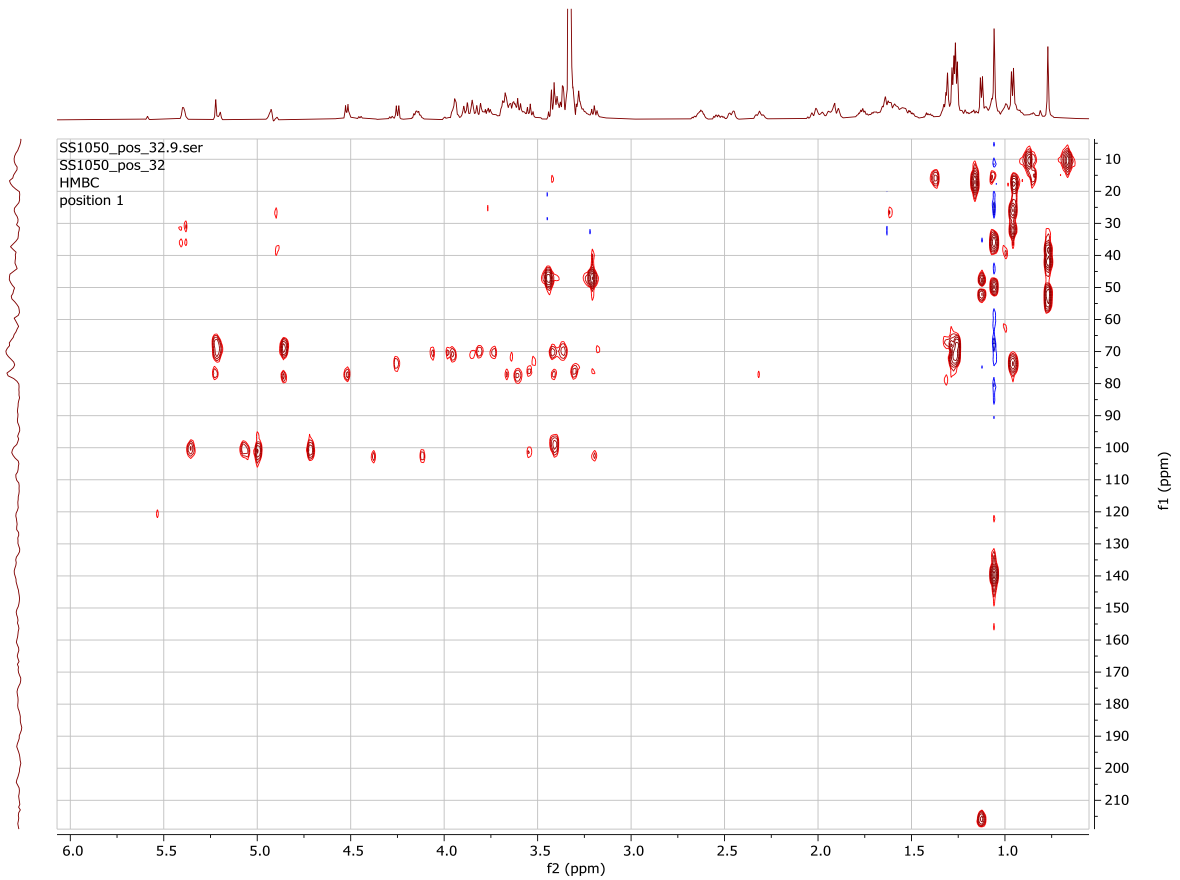


**Figure S44. gHMBCAD NMR spectrum for the intact molecule of the switchgrass saponin SS1050.**

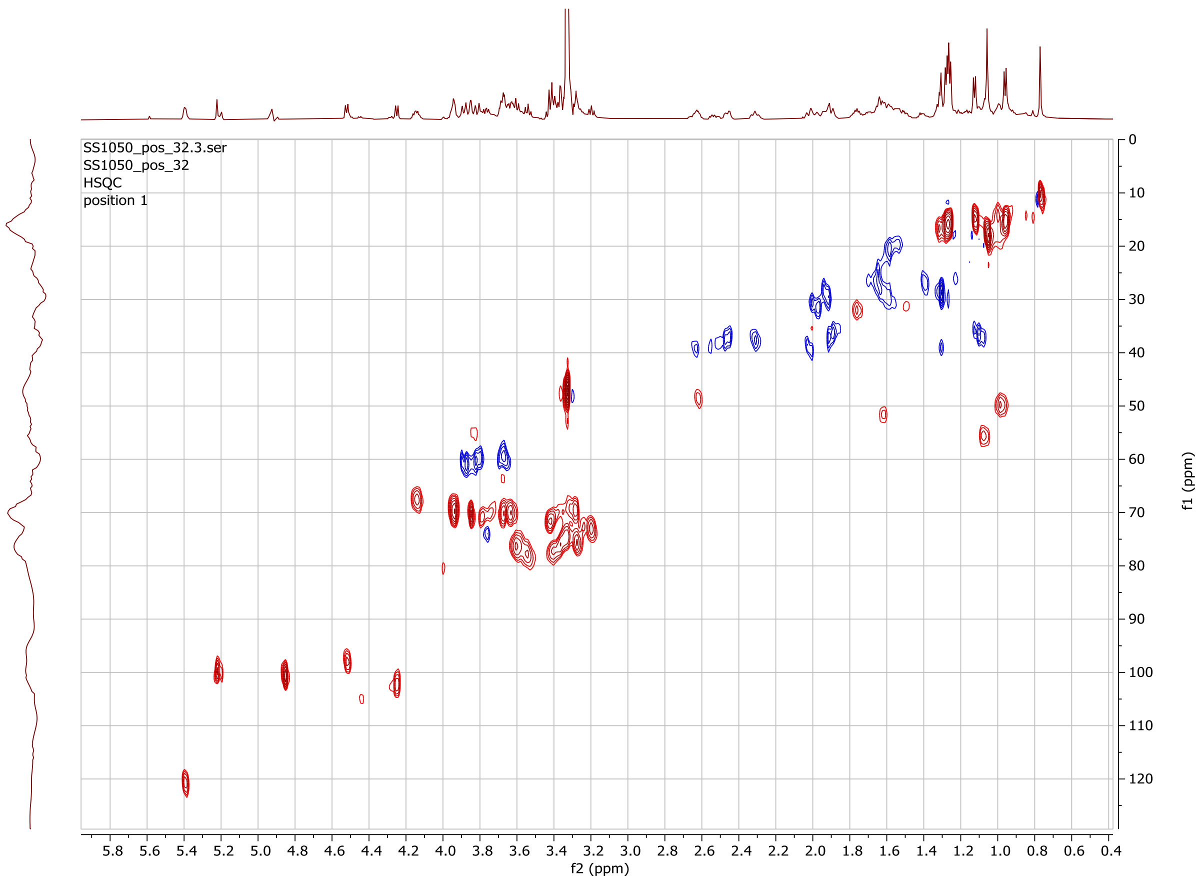


**Figure S45. gHSQCAD NMR spectrum for the intact molecule of the switchgrass saponin SS1050.**

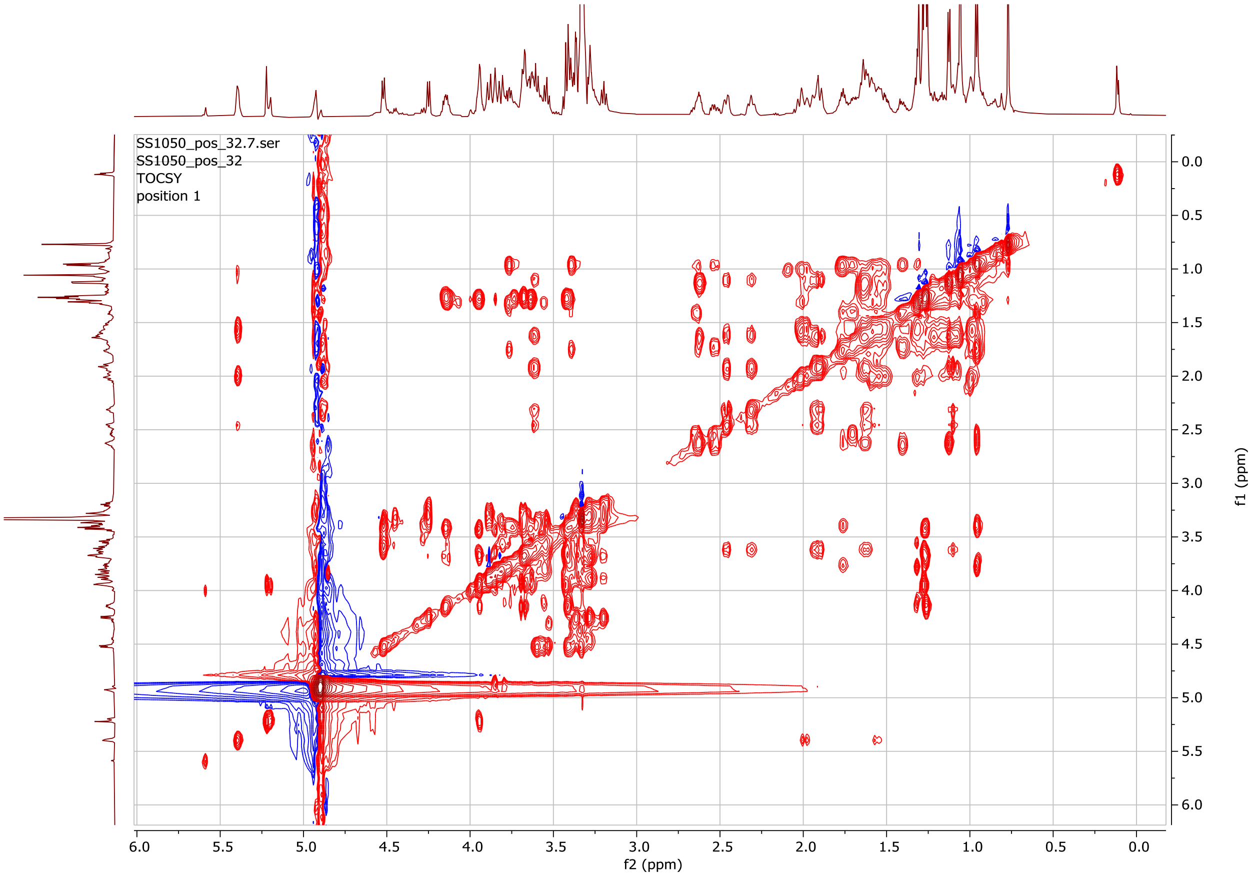


**Figure S46. gTOCSY NMR spectrum for the intact molecule of the switchgrass saponin SS1050.**

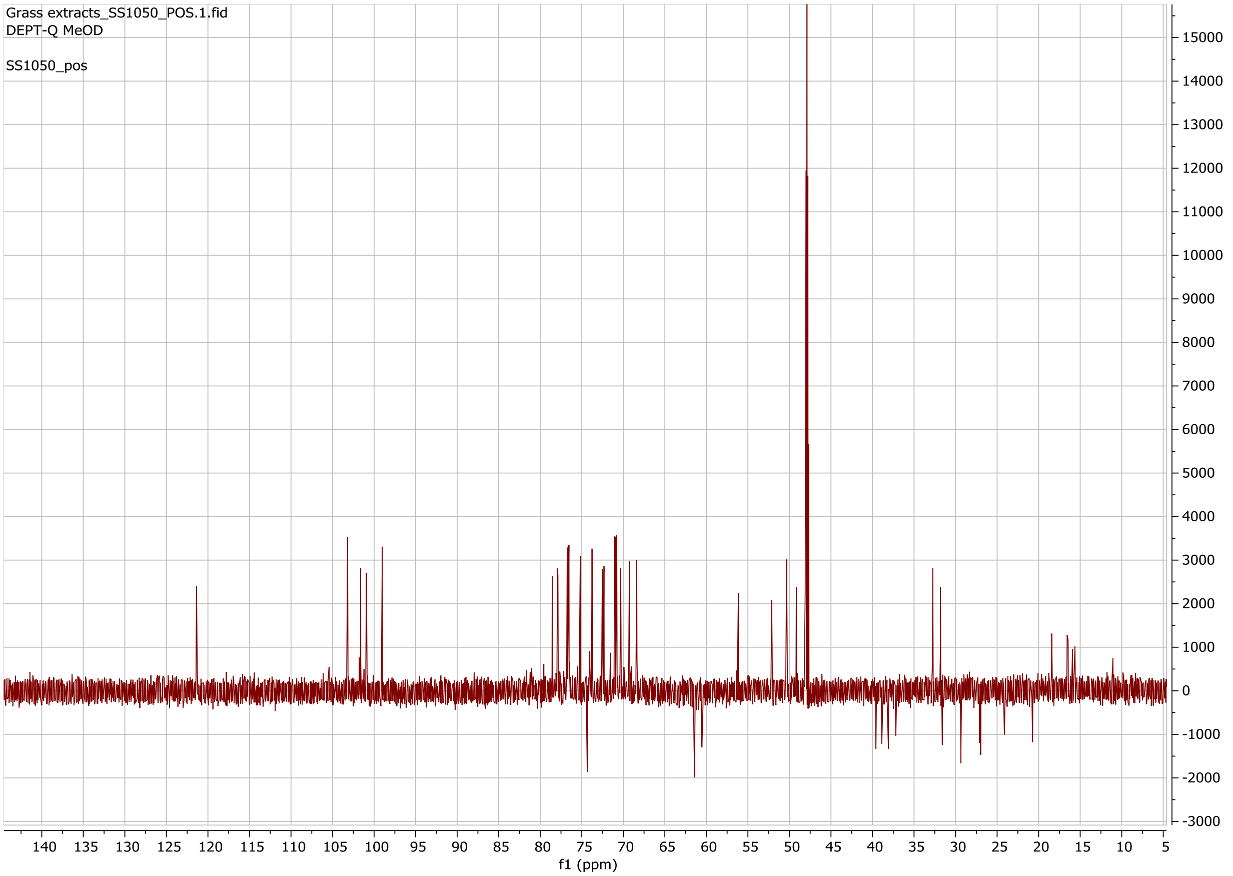


**Figure S47. DEPT-Q ($^{13}C$) NMR spectrum for the intact molecule of the switchgrass saponin SS1050.**

**Table S20. Chemical shift assignments of the aglycone of the switchgrass saponin SS1254. Chemical formular: $C_{29}H_{46}O_5$**

| Position | 1H δ (ppm) | 13C δ (ppm) | Annotation |
|---|---|---|---|
| 1 | 1.89/1.09 | 35.53 | CH2 |
| 2 | 1.92/1.61 | 29.36 | CH2 |
| 3[a] | 3.63 | 70.11 | CH3 |
| 4 | 2.45/2.3 | 37.14 | CH2 |
| 5[b] | | 139.97 | olefinic =C |
| 6 | 5.4 | 120.76 | olefinic =CH |
| 7 | 2/1.57 | 31.78 | CH2 |
| 8 | 1.67 | 30.76 | CH |
| 9 | 0.99 | 49.72 | CH |
| 10 | | 35.98 | quaternary C |
| 11 | 1.58/1.51 | 20.29 | CH2 |
| 12 | 1.81/1.23 | 38.14 | CH2 |
| 13 | | 40.47 | quaternary C |
| 14 | 1.16 | 55.79 | CH |
| 15 | 2.04/1.28 | 31.53 | CH2 |
| 16 | 4.58 | 79.91 | CH |
| 17 | 1.79 | 61.94 | CH |
| 18 | 0.85 | 14.59 | CH3 |
| 19 | 1.07 | 17.96 | CH3 |
| 20 | 2.1 | 38.92 | CH |
| 21 | 1.02 | 13.88 | CH3 |
| 22 | | 109.71 | quaternary C |
| 23 | 1.74/1.65 | 35.45 | CH2 |
| 24 | 1.58/1.34 | 27.16 | CH2 |
| 25 | 1.78 | 31.73 | CH |
| 26 | 3.97/3.9 | 68.93 | CH2 |
| 27 | 0.97 | 15.01 | CH3 |
| 28[c] | | 171.08 | Ester carbonyl C |
| 29 | 2.05 | 18.56 | CH3 |

[a] The C-3 carbon connects to a sugar anomeric proton, 5.59 *ppm*, through HMBC.
[b] The C-5 carbon chemical shift, 139.97 *ppm*, was determined by HMBC.
[c] The C-28 carbon chemical shift, 171.08 *ppm*, was determined by HMBC.

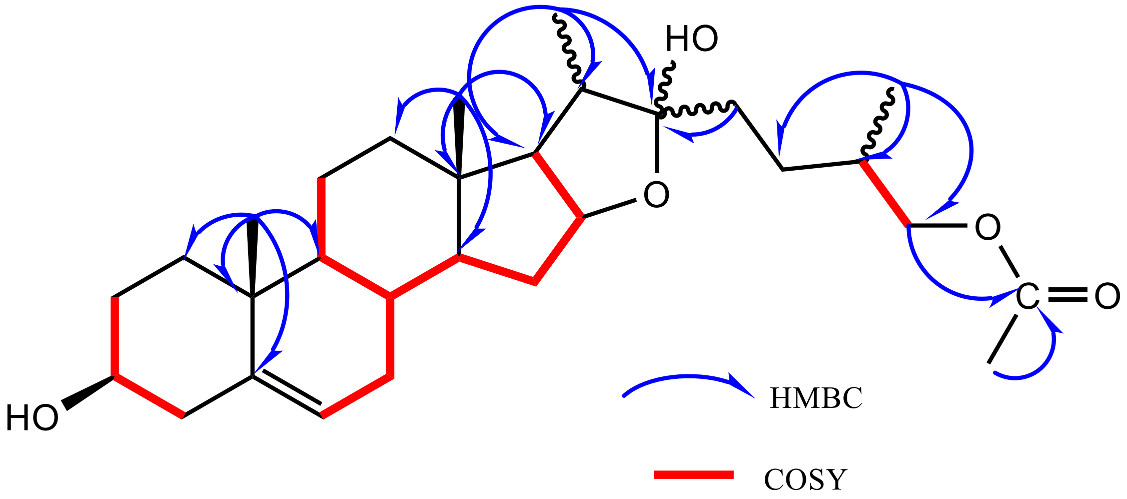


**Figure S48. The key COSY (red, bold) and HMBC (blue arrow) correlations for the aglycone of the switchgrass saponin SS1254.**

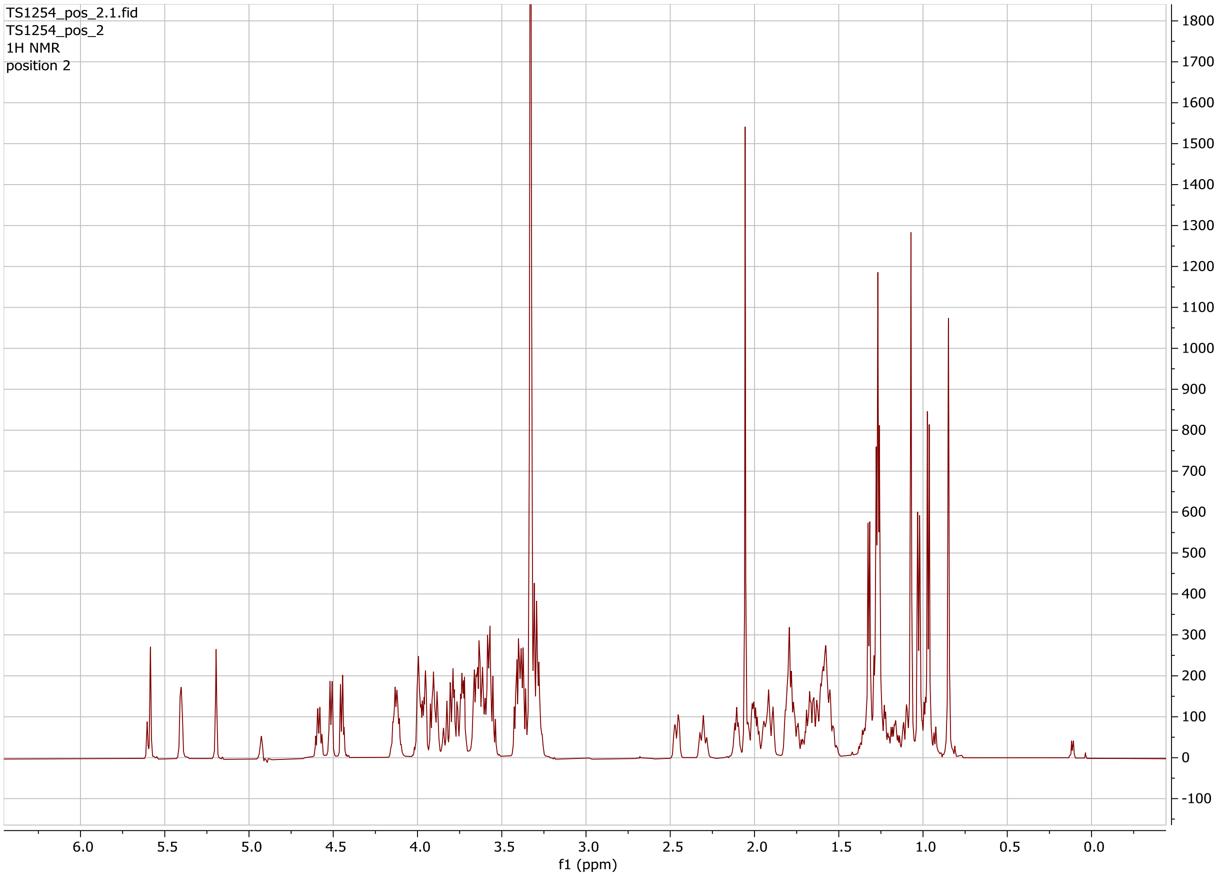


**Figure S49. $^{1}H$ NMR spectrum for the intact molecule of the switchgrass saponin SS1254.**

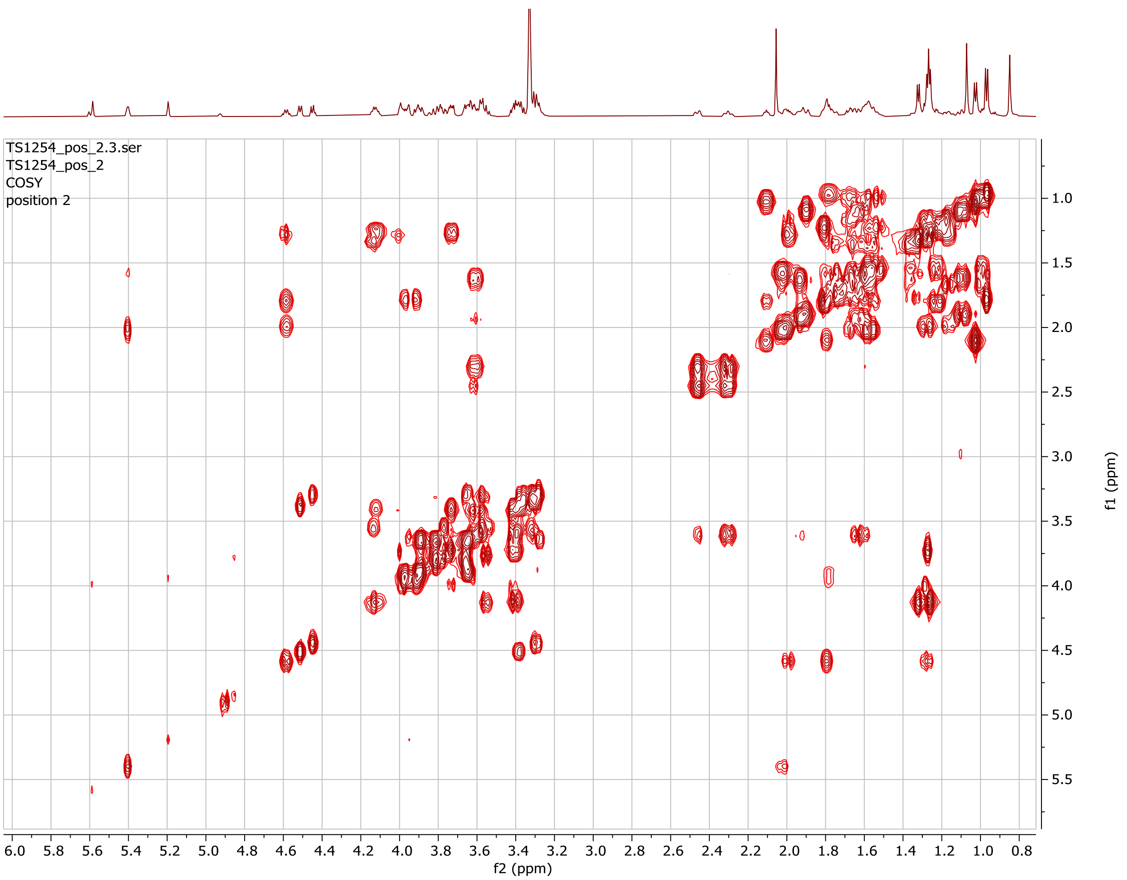


**Figure S50. gCOSY NMR spectrum for the intact molecule of the switchgrass saponin SS1254.**

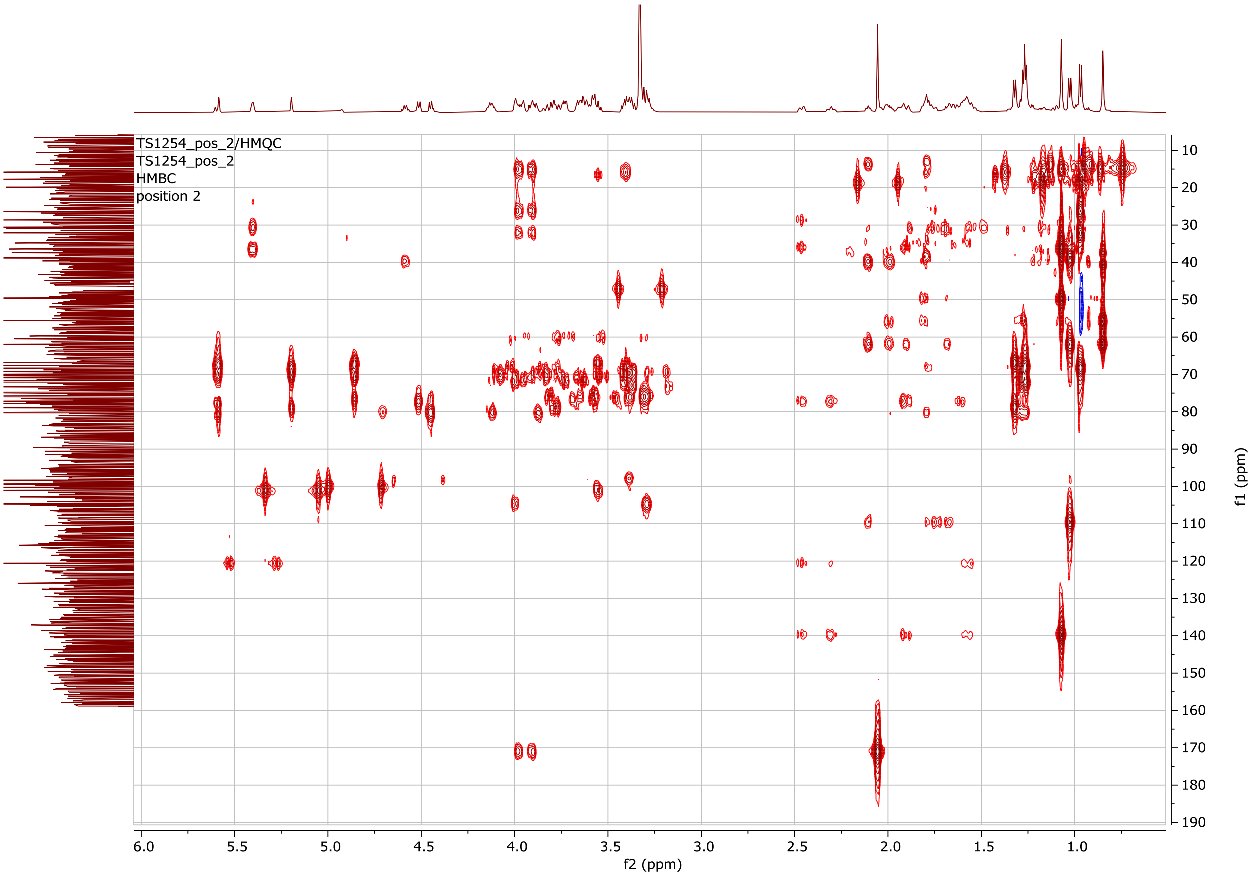


**Figure S51. gHMBCAD NMR spectrum for the intact molecule of the switchgrass saponin SS1254.**

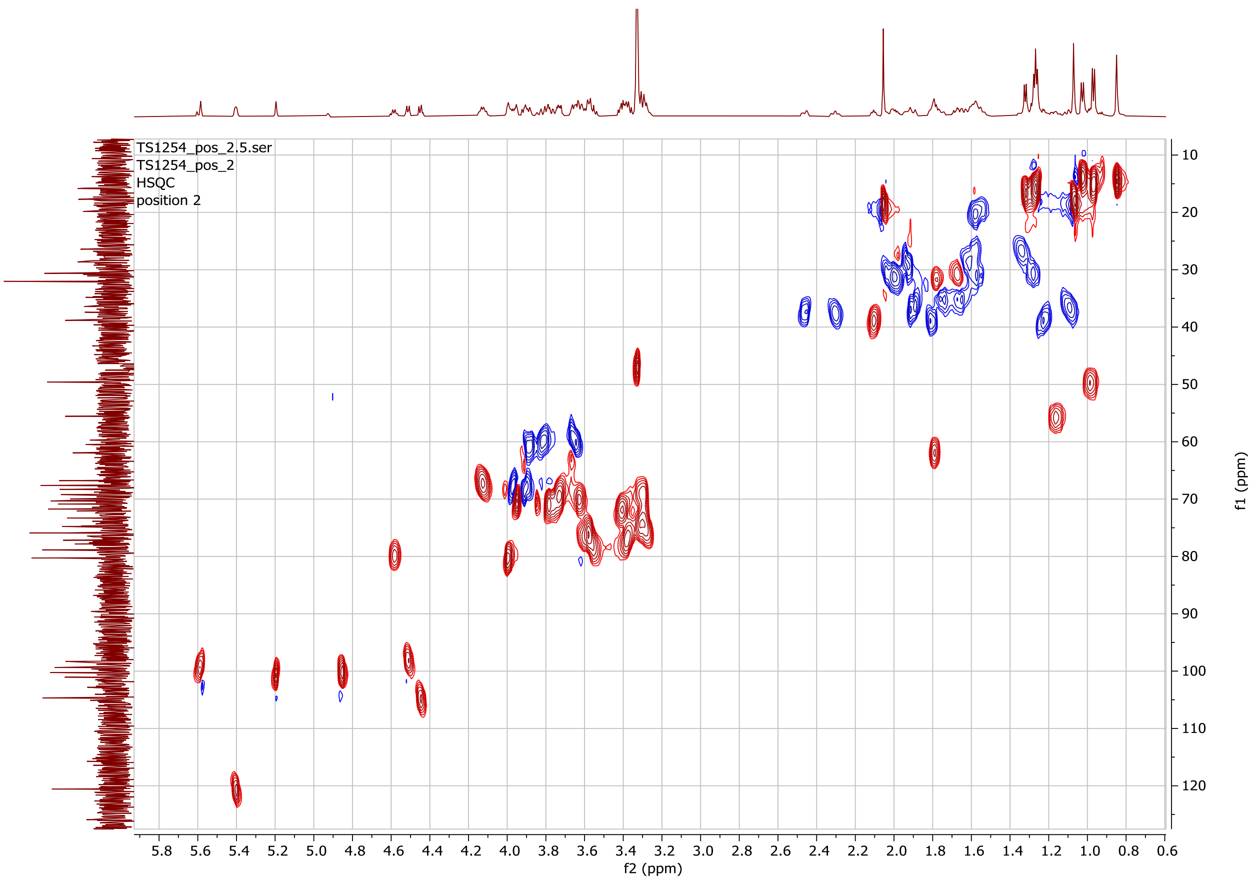


**Figure S52. gHSQCAD NMR spectrum for the intact molecule of the switchgrass saponin SS1254.**

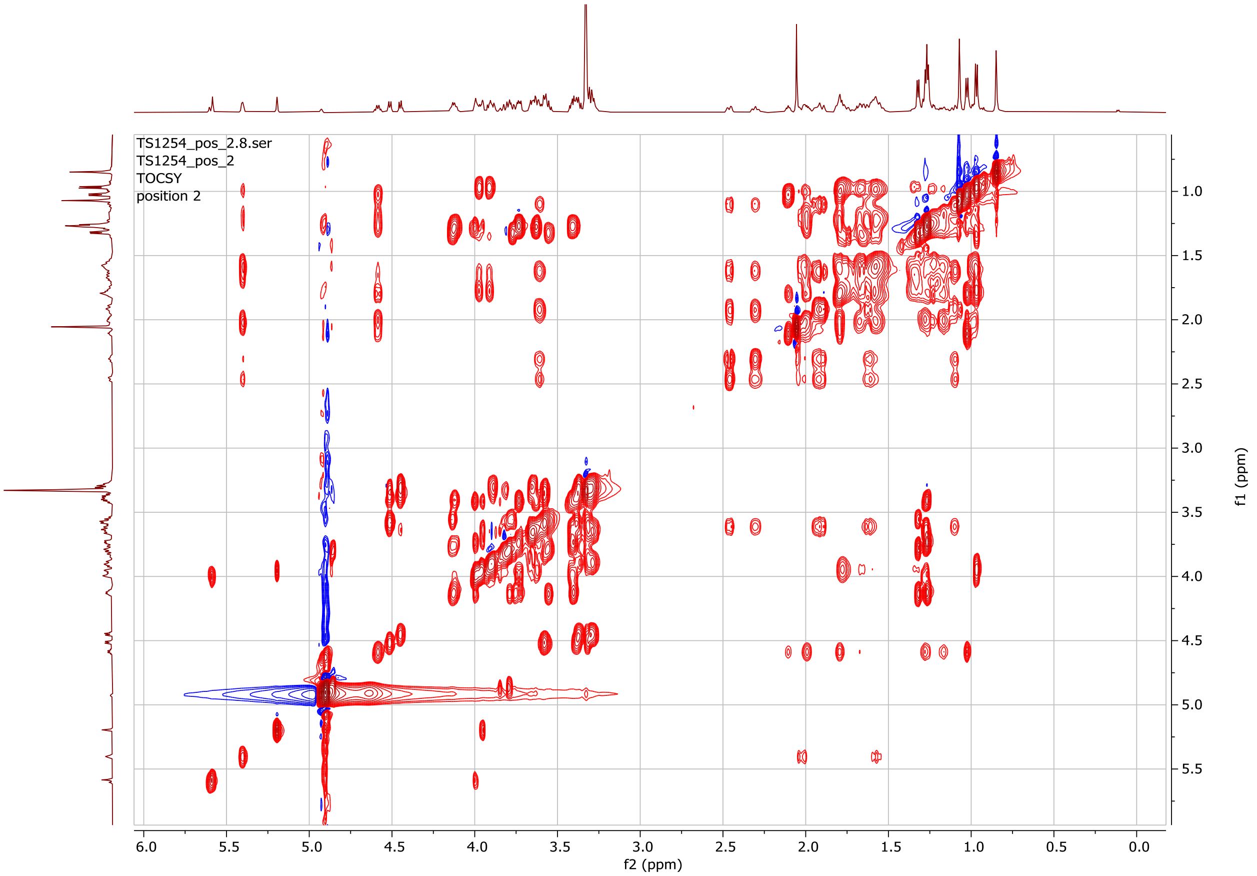


**Figure S53. gTOCSY NMR spectrum for the intact molecule of the switchgrass saponin SS1254.**

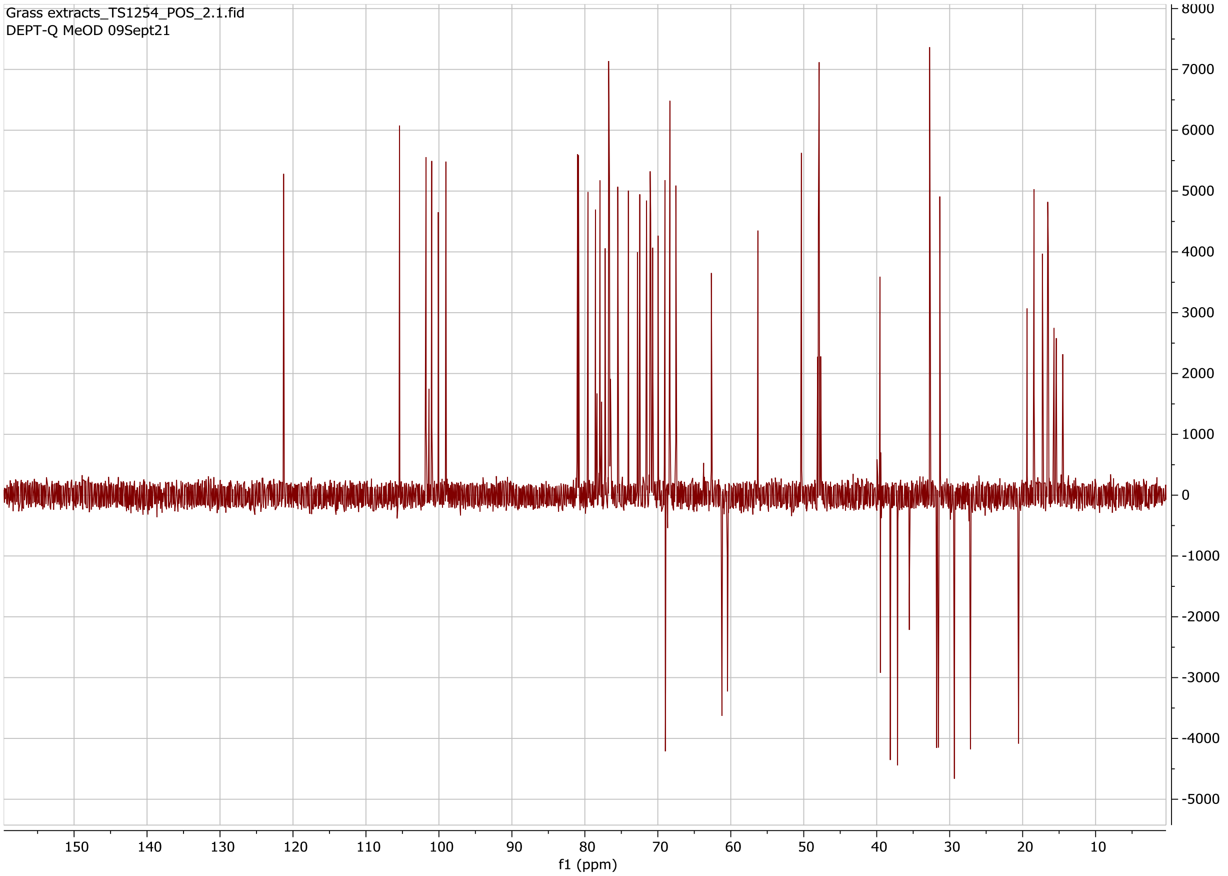


**Figure S54. DEPT-Q ($^{13}C$) NMR spectrum for the intact molecule of the switchgrass saponin SS1254.**

**Table S21. Chemical shift assignments of the aglycone of the switchgrass saponin SS1254. Chemical formular: $C_{29}H_{46}O_6$**

| Position | 1H δ (ppm) | 13C δ (ppm) | Annotation |
|---|---|---|---|
| 1 | 1.9/1.1 | 35.35 | CH2 |
| 2 | 1.94/1.61 | 29.36 | CH2 |
| 3[a] | 3.63 | 70.05 | CH |
| 4 | 2.45/2.3 | 37.16 | CH2 |
| 5[b] | 1.07 | 139.71 | olefinic =C |
| 6 | 5.4 | 120.77 | olefinic =CH |
| 7 | 2.04/1.59 | 31.5 | CH2 |
| 8 | 1.65 | 31.31 | CH |
| 9 | 0.96 | 49.53 | CH |
| 10 | 1.07 | 35.88 | quaternary C |
| 11 | 1.64/1.54 | 20.3 | CH2 |
| 12 | 1.93/1.31 | 38.14 | CH2 |
| 13 | 2.27 | 43.98 | quaternary C |
| 14 | 1.72 | 51.54 | CH |
| 15 | 2.05/1.25 | 30.83 | CH2 |
| 16 | 4.17 | 88.6 | CH |
| 17 | | 89.37 | quaternary C |
| 18 | 0.86 | 15.24 | CH3 |
| 19 | 1.06 | 17.92 | CH3 |
| 20 | 2.26 | 41.63 | CH |
| 21 | 0.94 | 7.93 | CH3 |
| 22 | 1.76 | 110.14 | quaternary C |
| 23 | 1.78/1.7 | 35.25 | CH2 |
| 24 | 1.56/1.36 | 26.82 | CH2 |
| 25 | 1.78 | 31.69 | CH |
| 26 | 3.97/3.91 | 68.93 | CH2 |
| 27 | 0.97 | 15.07 | CH3 |
| 28 | 3.97 | 171.12 | Ester carbonyl C |
| 29 | 2.05 | 18.5 | CH3 |

[a] The C-3 carbon connects to a sugar anomeric proton, 5.6 *ppm*, through HMBC.
[b] The C-5 carbon chemical shift, 139.71 *ppm*, was determined by HMBC.

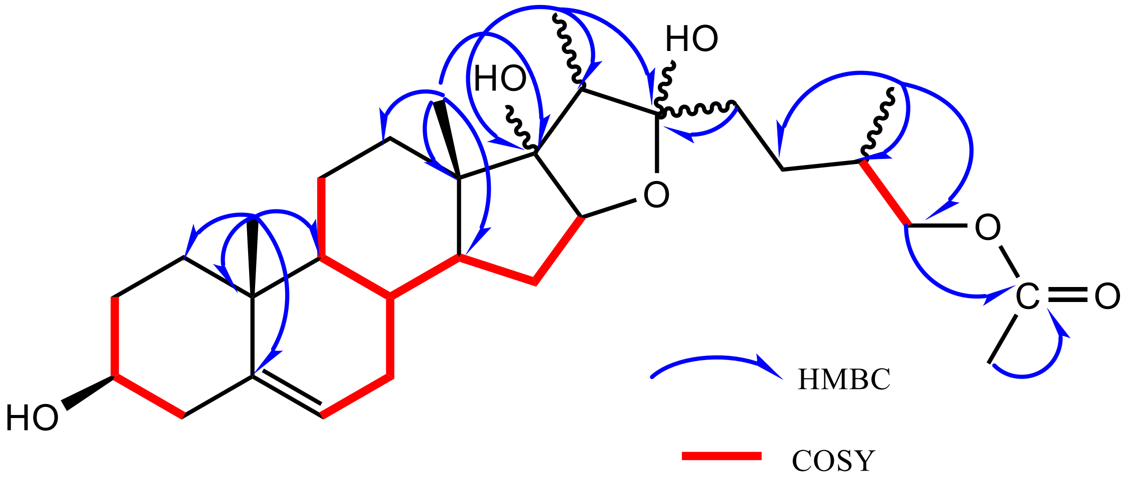


**Figure S55. The key COSY (red, bold) and HMBC (blue arrow) correlations for the aglycone of the switchgrass saponin SS1089.**

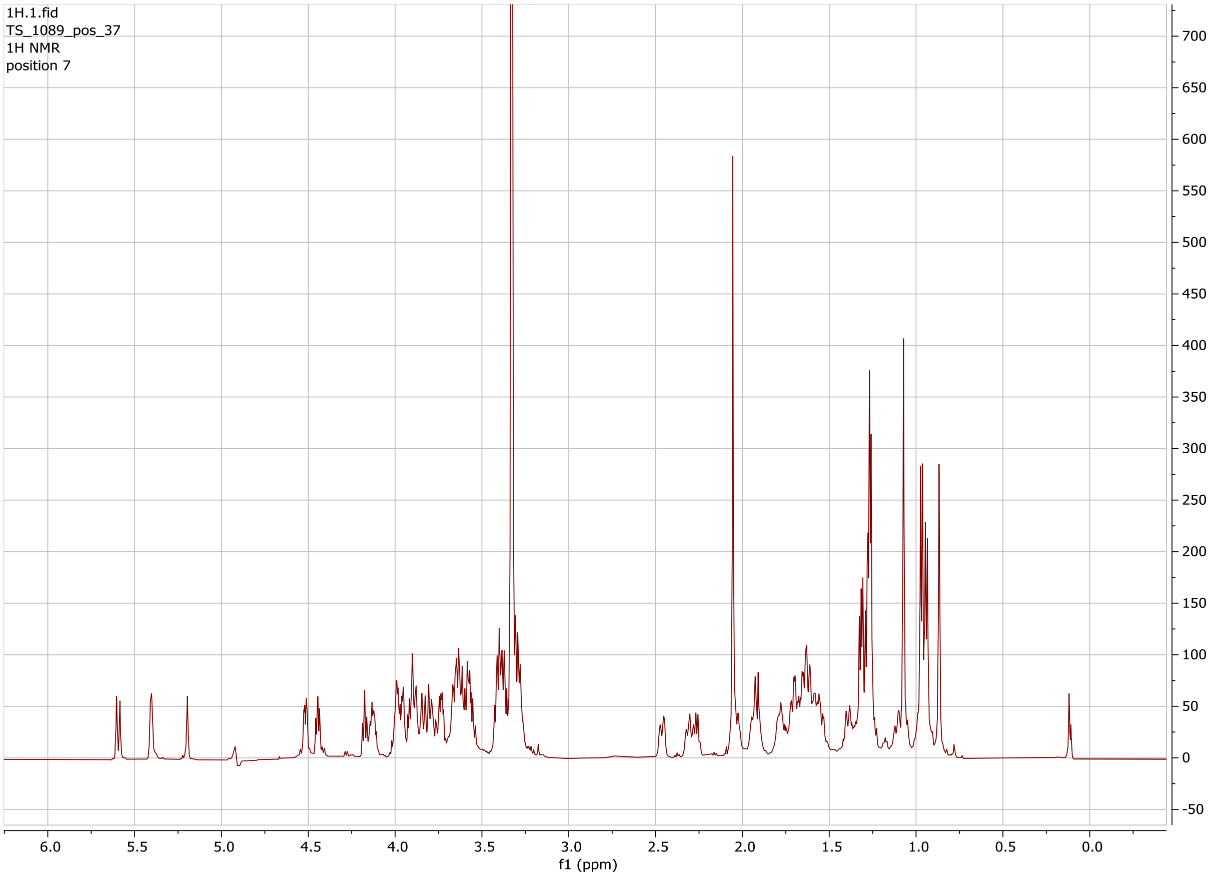


**Figure S56. [1]H NMR spectrum for the intact molecule of the switchgrass saponin SS1089.**

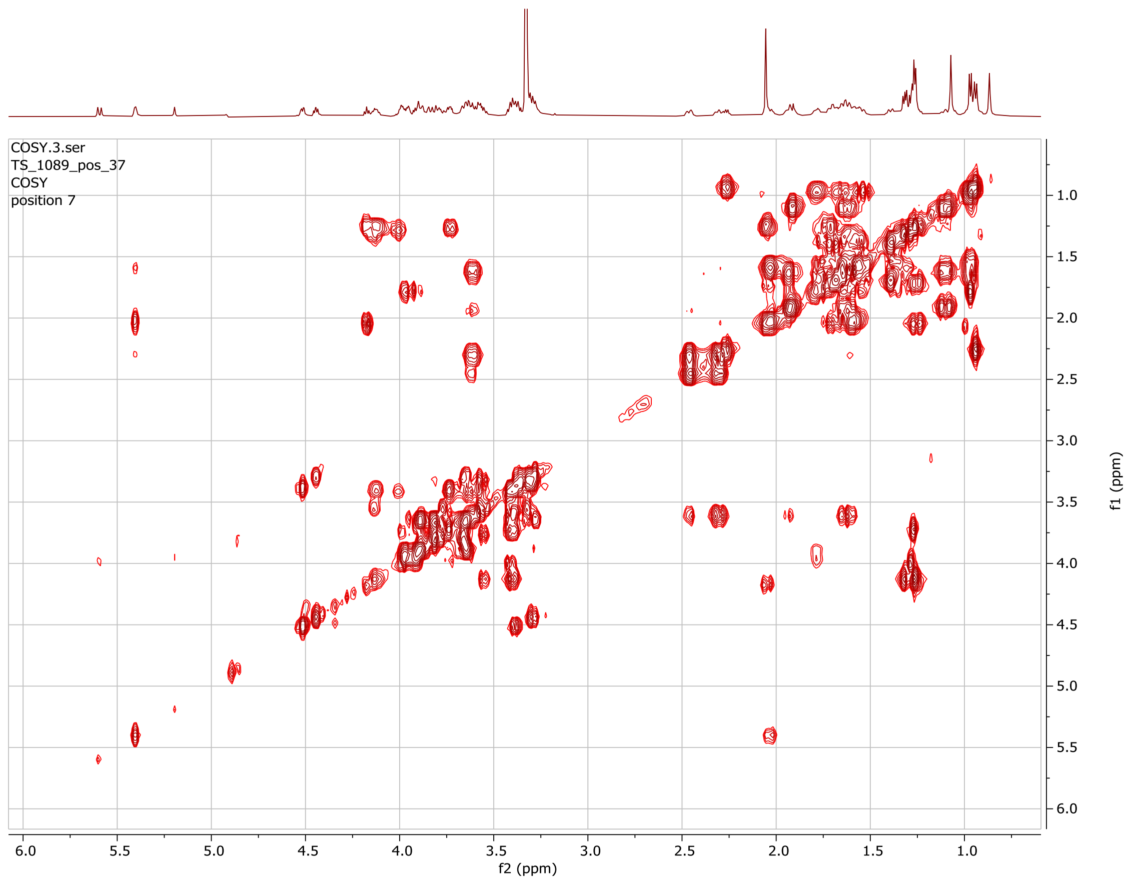


**Figure S57. gCOSY NMR spectrum for the intact molecule of the switchgrass saponin SS1089.**

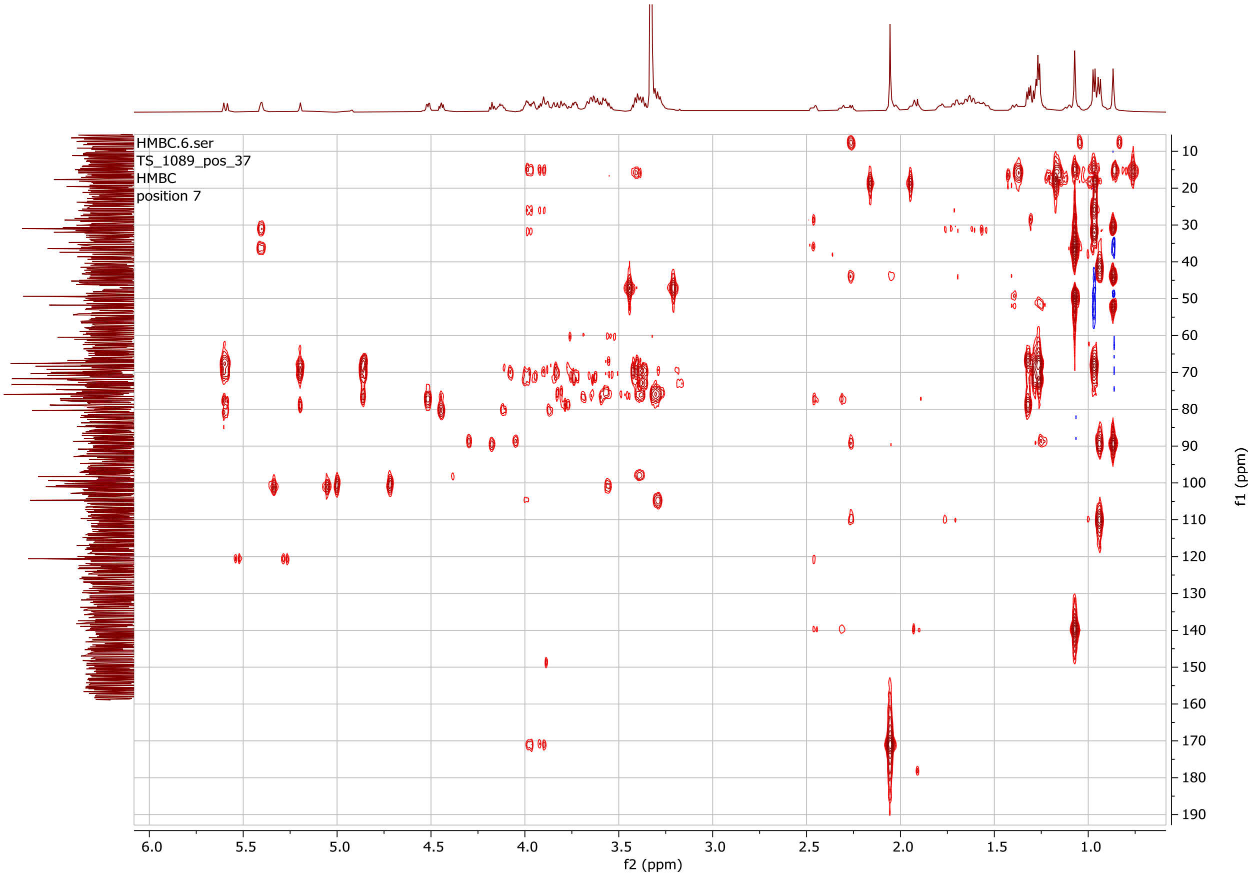


**Figure S58. gHMBCAD NMR spectrum for the intact molecule of the switchgrass saponin SS1089.**

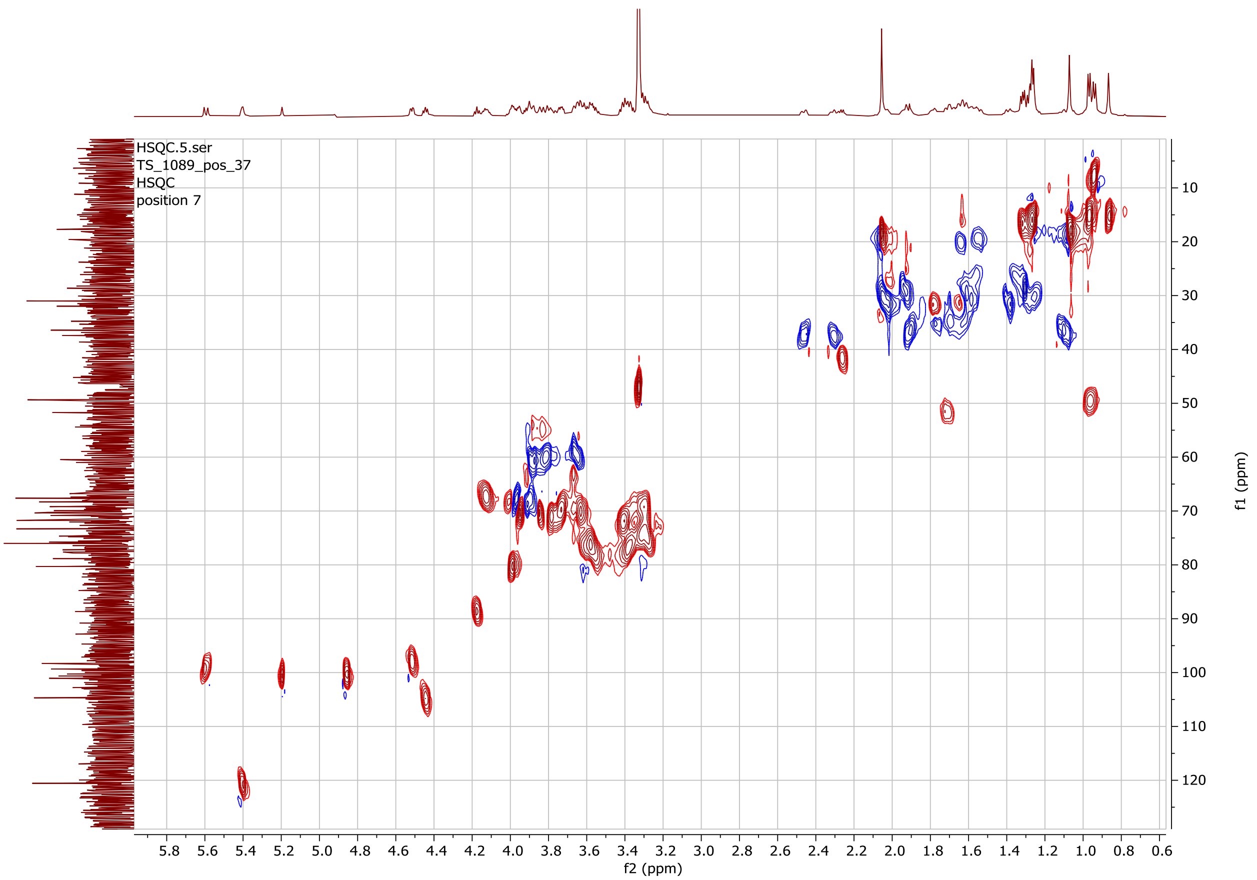


**Figure S59. gHSQCAD NMR spectrum for the intact molecule of the switchgrass saponin SS1089.**

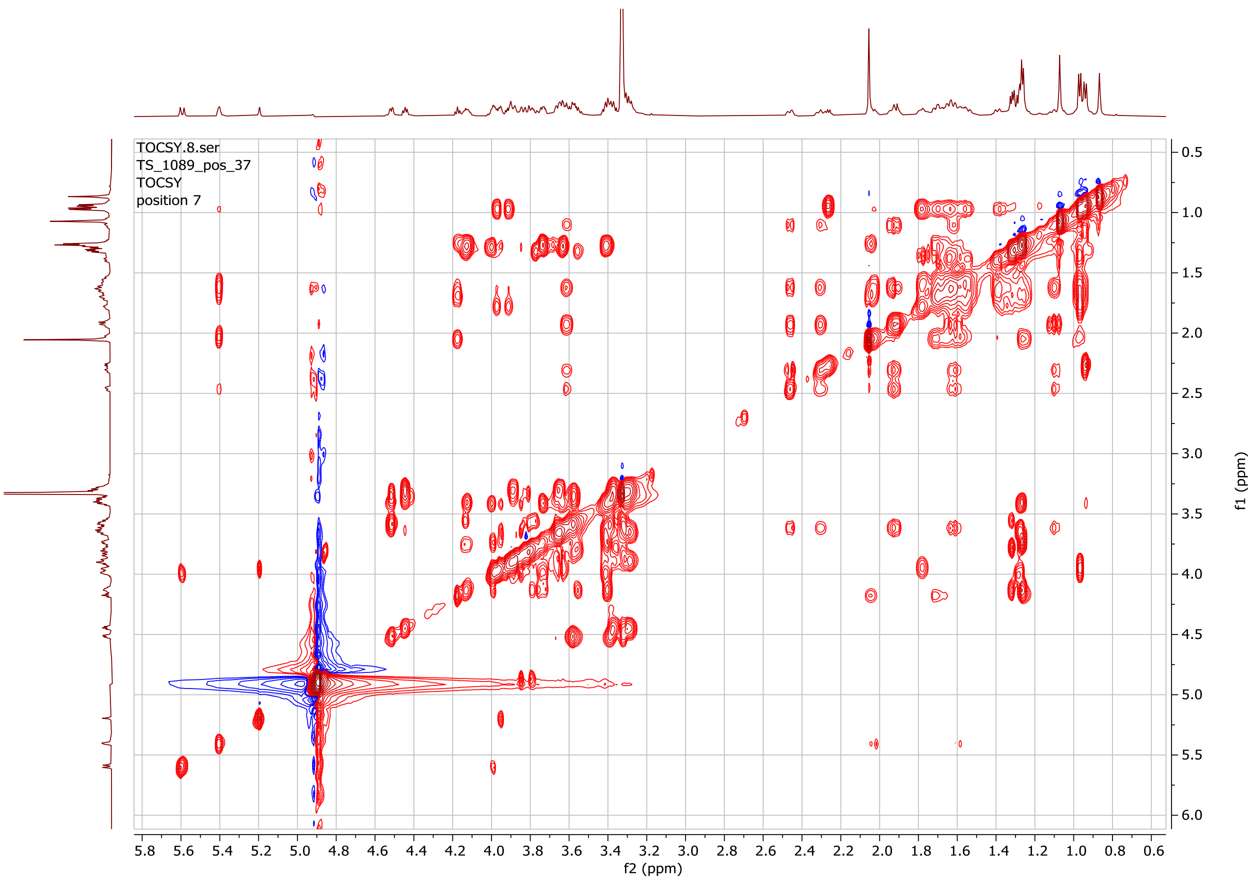


**Figure S60. gTOCSY NMR spectrum for the intact molecule of the switchgrass saponin SS1089.**

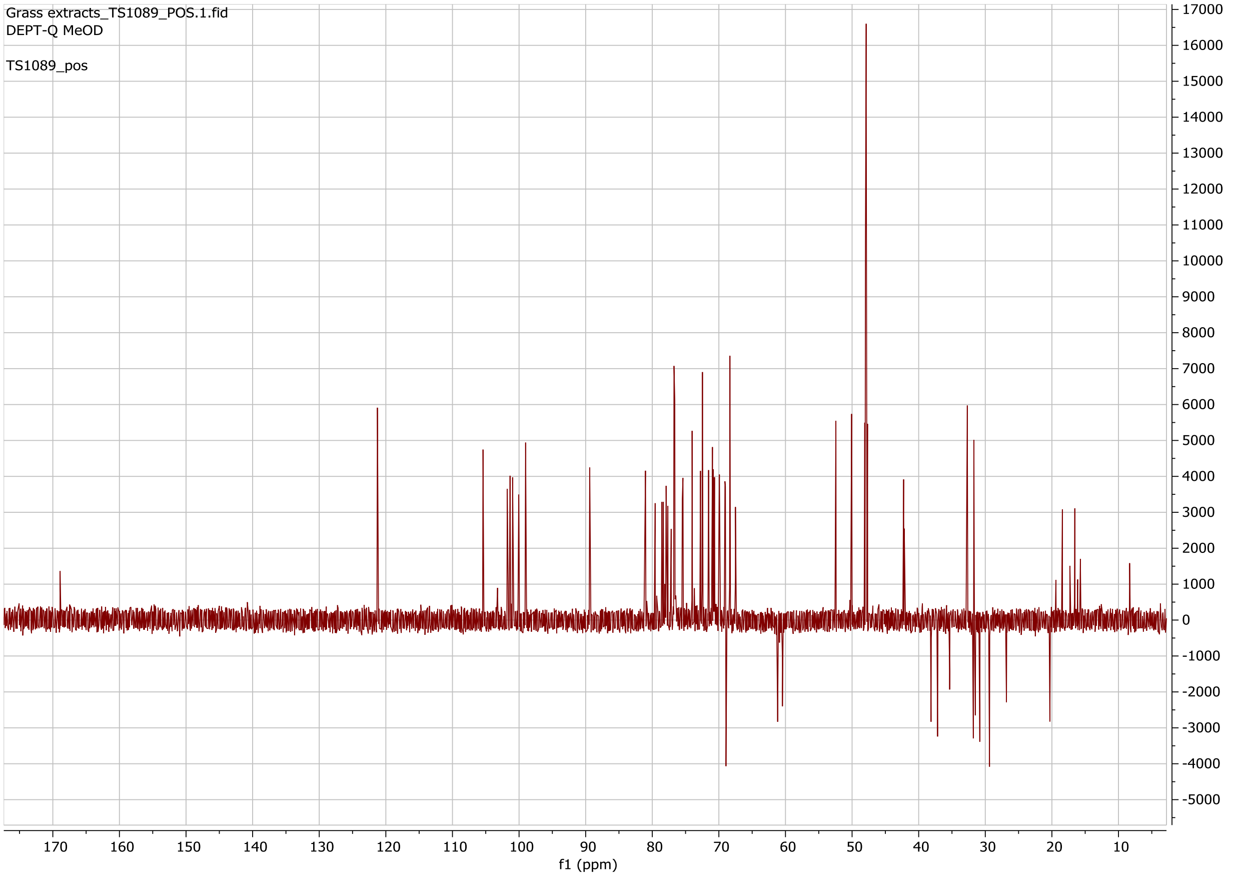


**Figure S61. DEPT-Q ($^{13}C$) NMR spectrum for the intact molecule of the switchgrass saponin SS1089.**

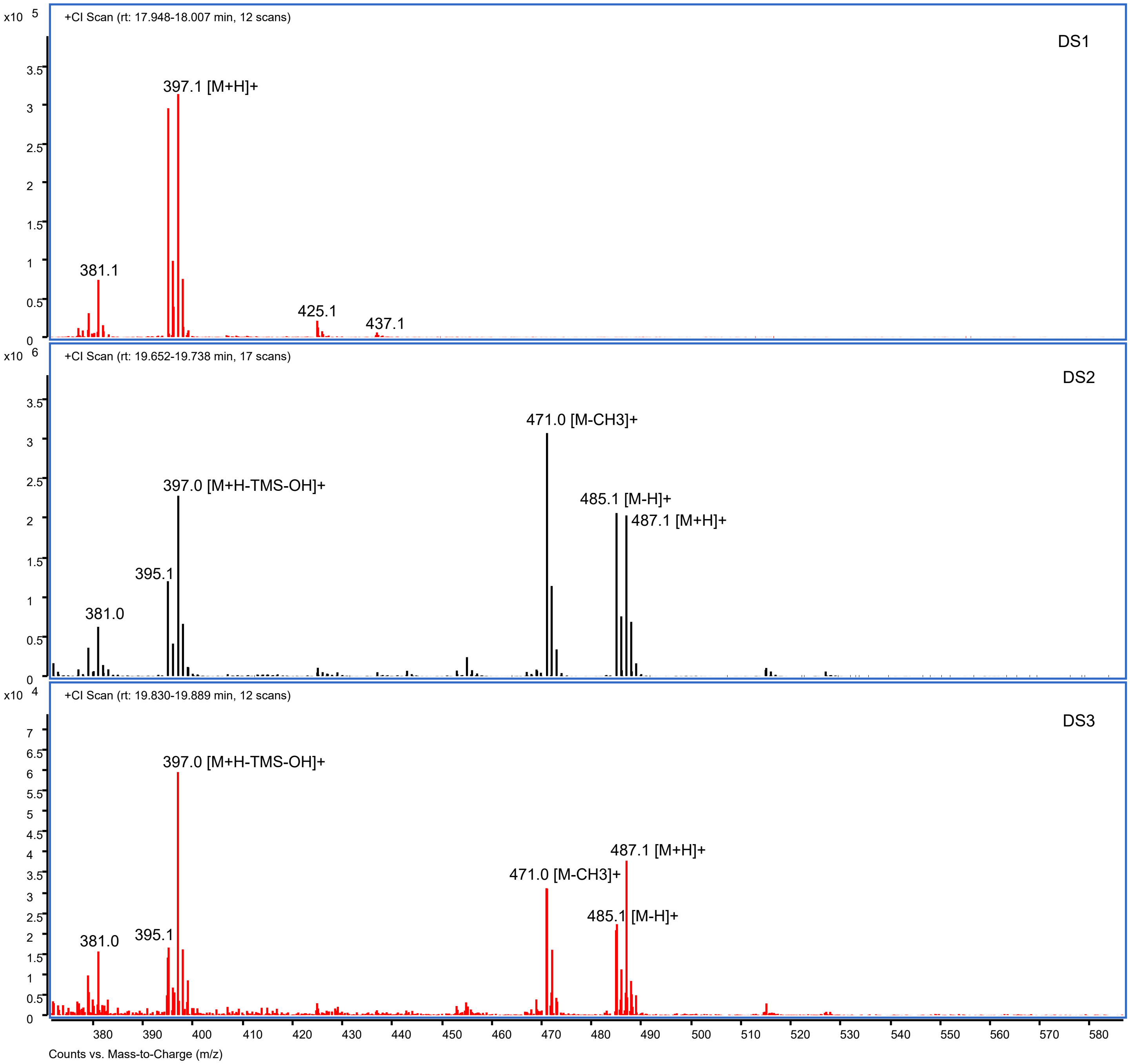


**Figure S62. The (CI) GC-MS spectral patterns of the sapogenin peaks detected in the standard and switchgrass samples. The key precursor ionsfor each TMS derivatized sapogenin peak were documented in Table 2.**

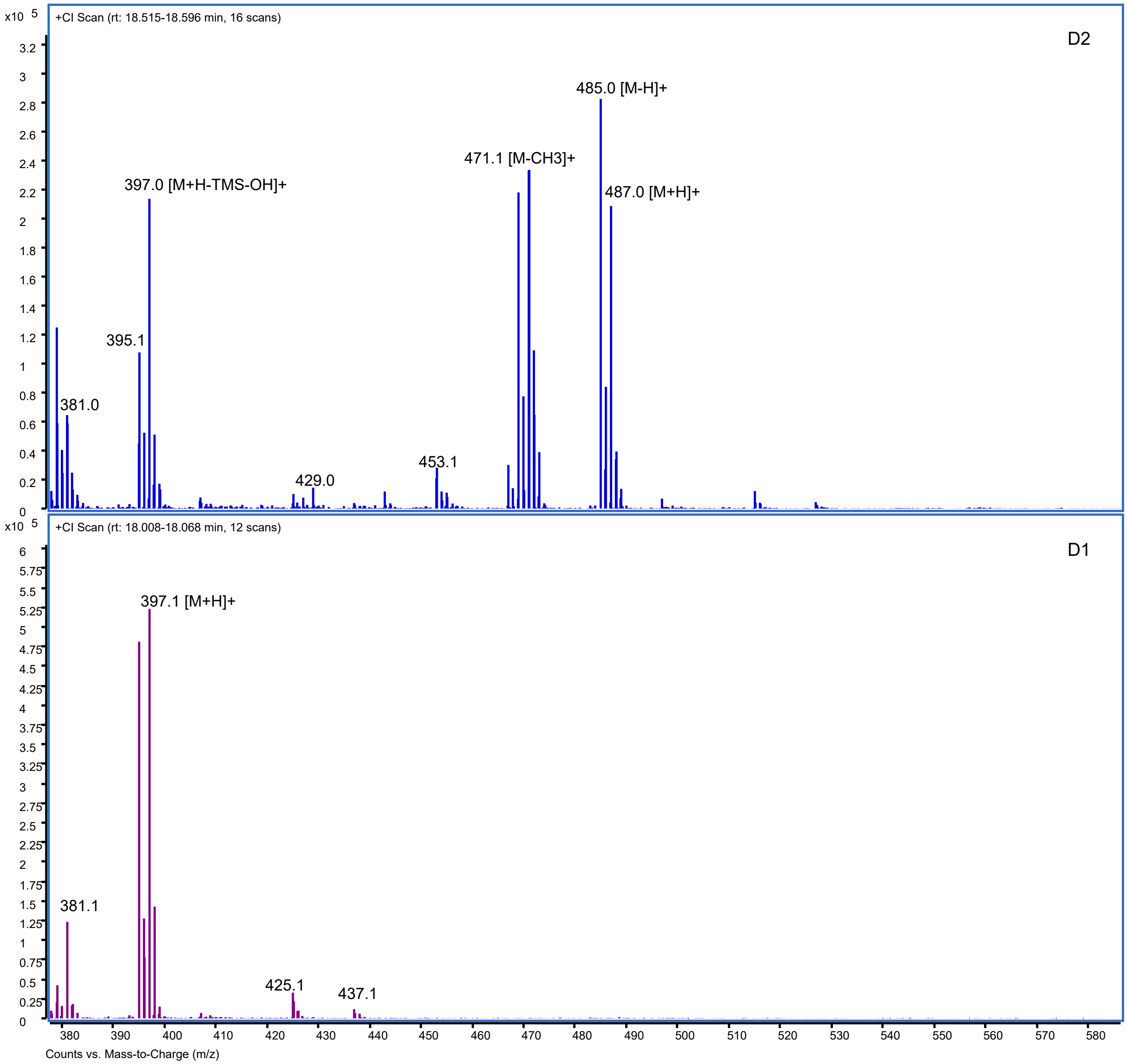


**Figure S62 (continued).**

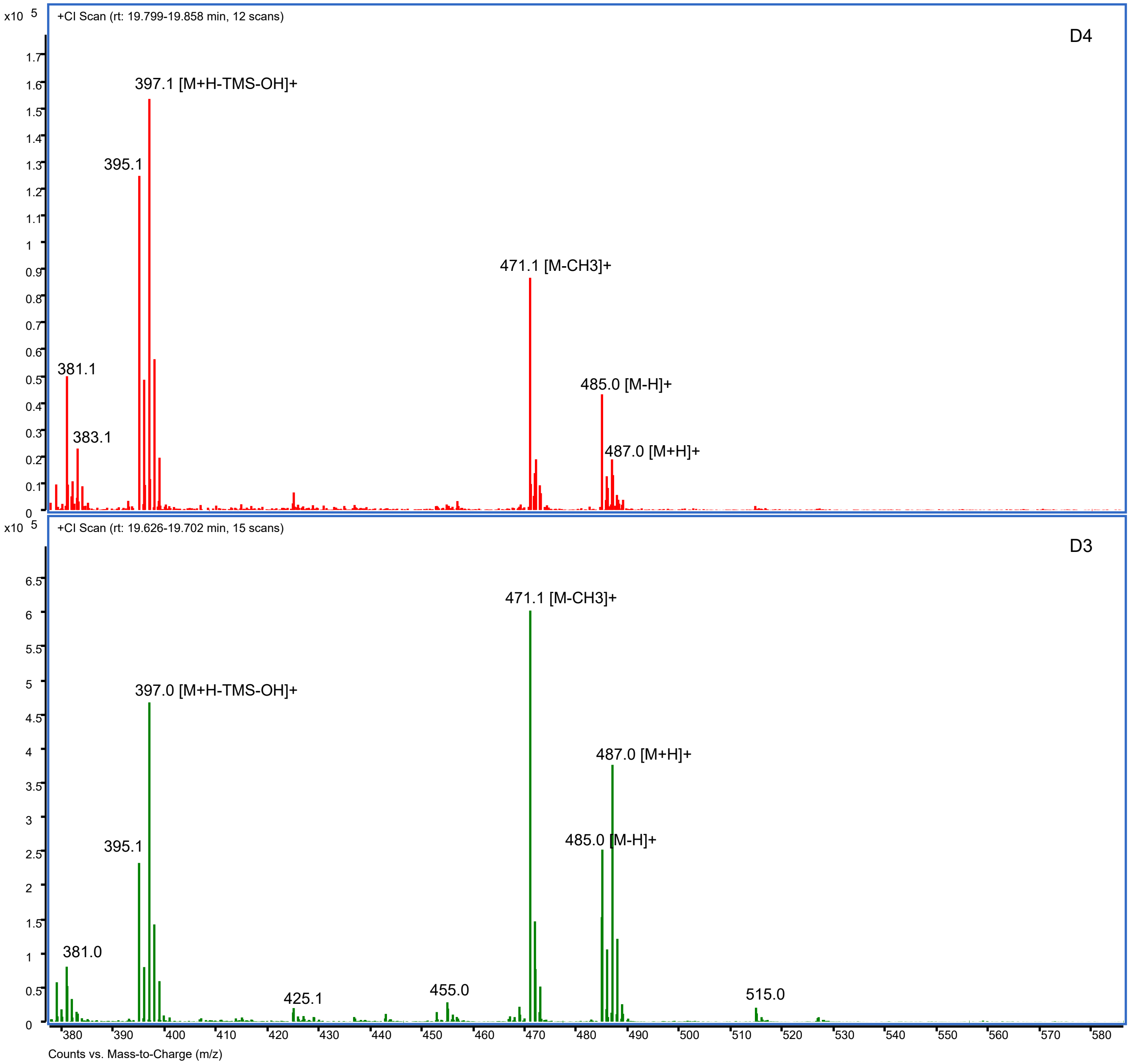


**Figure S62 (continued).**

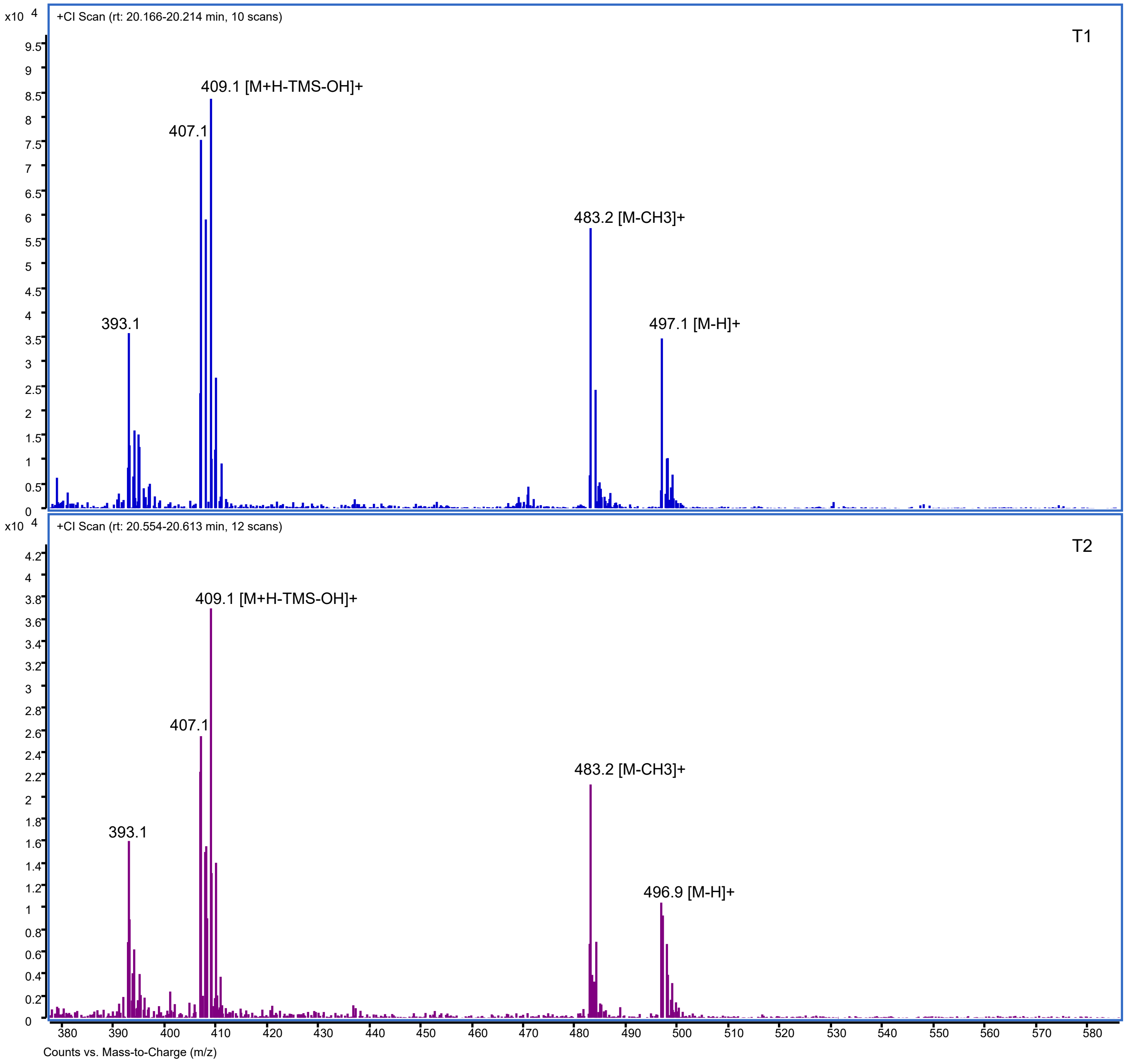


**Figure S62 (continued).**

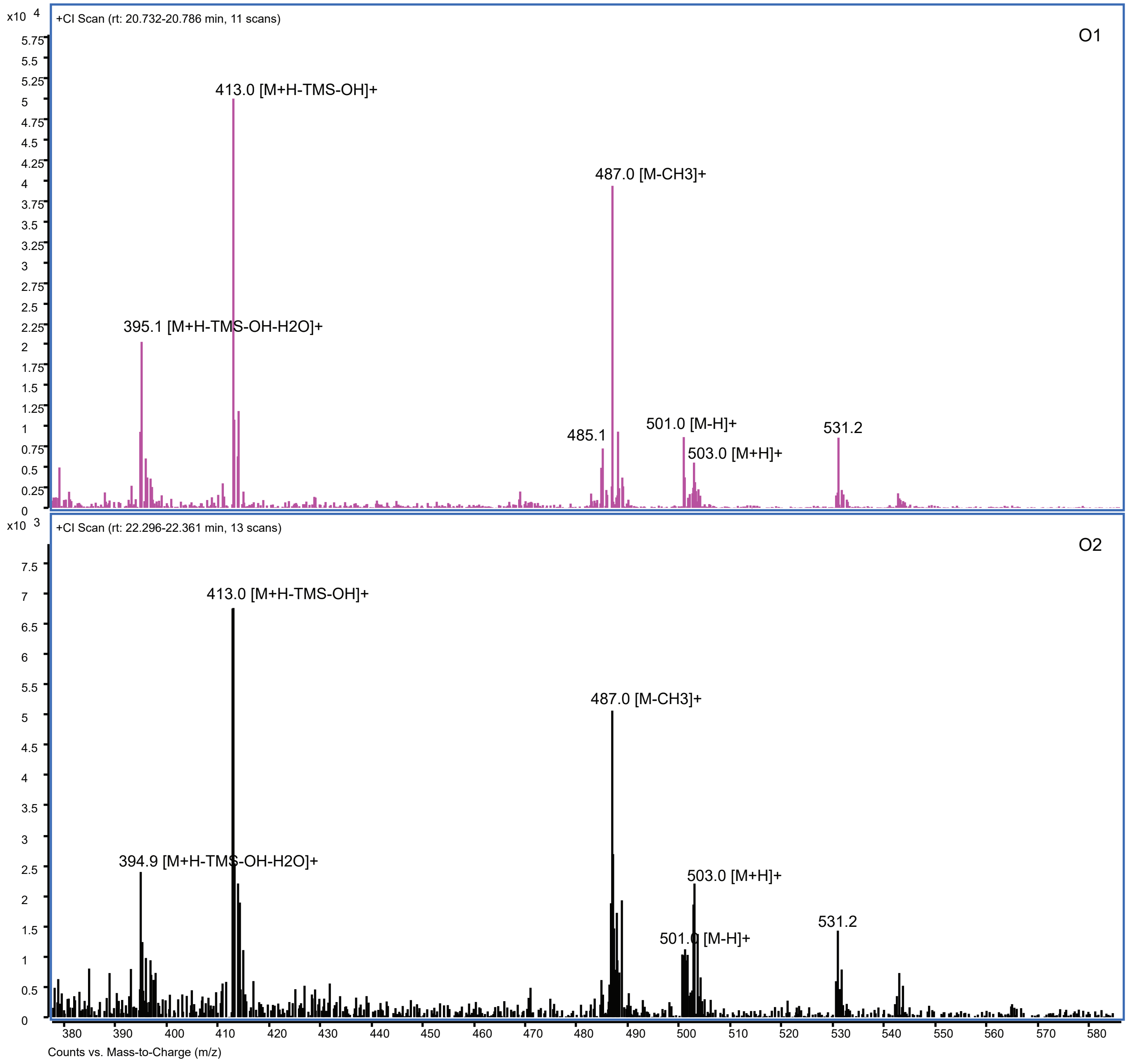


**Figure S62 (continued).**

**Table S22**. **Relative sapogenin concentrations in different tissues of the six switchgrass cultivars in this study.** Relative concentrations were calculated based on the total ion chromatograms (TIC) peak areas of the eight sapogenins relative to the peak areas of the diosgenin standard (**Materials and Methods**). Values are the average of triplicates ± SEM in ng/mg dw. ND, not detected or below detection level; ng/mg dw, nanograms per milligram of dry tissue weight; LB, leaf blade; LS, leaf sheath, St, stem; Rhi, rhizome; Rt, root.

| Cultivar Name | Tissue Type | D1 | D2 | D3 | D4 | O1 | O2 | T1 | T2 | Total |
|---|---|---|---|---|---|---|---|---|---|---|
| Dacotah | LB | 140±2 | 71±13 | 783±168 | 649±62 | ND | 194±75 | 99±13 | 32±10 | 1968±261 |
| | LS | 53±3 | 71±6 | 168±19 | 115±9 | 24±0 | 51±4 | 16±8 | 24±7 | 522±25 |
| | St | 111±8 | 36±11 | 51±13 | 49±9 | 2±1 | 1ND | 2±1 | 6±3 | 268±11 |
| | Rhi | 95±31 | 171±18 | 461±161 | 146±37 | 28±8 | 210±11 | ND | 5±0 | 1116±232 |
| | Rt | 79±16 | 697±107 | 160±22 | 117±7 | 124±35 | 62±16 | 1±0 | ND | 1240±193 |
| Summer | LB | 148±25 | 56±15 | 969±165 | 509±50 | 1±0 | 57±24 | 157±64 | 100±16 | 1997±254 |
| | LS | 69±14 | 51±15 | 164±13 | 108±14 | 4±3 | 15±2 | 11±4 | 26±5 | 449±32 |
| | St | 83±28 | 49±8 | 38±6 | 34±6 | 1±1 | 1±0 | 4±0 | 1±0 | 212±35 |
| | Rhi | 123±61 | 189±40 | 779±133 | 217±62 | 80±21 | 194±48 | ND | 7±0 | 1589±366 |
| | Rt | 132±27 | 898±192 | 166±50 | 149±28 | 158±41 | 52±14 | ND | ND | 1554±337 |
| Cave-in-Rock | LB | 169±21 | 37±15 | 1516±125 | 293±155 | ND | 40±29 | 173±54 | 6±4 | 2233±210 |
| | LS | 89±37 | 48±5 | 218±32 | 95±14 | 2±0 | 6±4 | 33±13 | 2±1 | 493±54 |
| | St | 136±20 | 39±17 | 47±15 | 42±12 | 1±1 | 1±0 | 3±2 | 3±2 | 273±44 |
| | Rhi | 232±35 | 155±10 | 1912±48 | 454±259 | 183±33 | 92±47 | 118±0 | 18±7 | 3165±99 |
| | Rt | 91±16 | 553±31 | 156±11 | 114±23 | 584±84 | 128±19 | 4±3 | ND | 1632±149 |
| Alamo | LB | 184±48 | 59±0 | 1078±305 | 769±330 | ND | 1±1 | 271±9 | 40±12 | 2403±630 |
| | LS | 136±24 | 34±5 | 75±12 | 137±34 | 2±0 | 1±0 | 10±3 | 29±14 | 423±68 |
| | St | 144±35 | 17±0 | 32±9 | 78±29 | 1±1 | ND | 3±2 | 8±4 | 283±57 |
| | Rhi | 239±9 | 74±1 | 1653±234 | 634±126 | ND | 132±61 | 2±0 | 15±5 | 2748±360 |
| | Rt | 253±71 | 72±3 | 2412±400 | 286±72 | 1±1 | 510±222 | ND | 3±0 | 3536±350 |
| Kanlow | LB | 204±54 | 69±54 | 1619±54 | 441±54 | 75±54 | 73±54 | 191±54 | 40±54 | 2713±54 |
| | LS | 122±17 | 39±6 | 134±32 | 159±34 | ND | ND | 20±6 | 29±9 | 502±56 |
| | St | 89±11 | 37±1 | 40±3 | 80±9 | 2±1 | ND | 5±2 | 10±1 | 263±13 |
| | Rhi | 365±28 | 94±10 | 3349±532 | 689±118 | ND | 3±1 | 2±0 | 22±4 | 4523±624 |
| | Rt | 434±24 | 105±7 | 4238±911 | 276±50 | ND | 4±3 | 2±1 | 7±2 | 5067±952 |
| BoMaster | LB | 173±42 | 51±1 | 1262±280 | 584±192 | ND | 8±5 | 183±39 | 59±21 | 2321±103 |
| | LS | 100±11 | 1ND | 98±30 | 182±16 | ND | 3±2 | 19±8 | 50±14 | 462±22 |
| | St | 84±19 | 69±12 | 43±11 | 106±20 | 4±1 | ND | ND | 13±3 | 320±21 |
| | Rhi | 217±46 | 40±11 | 1108±506 | 435±62 | ND | 63±31 | 1±0 | 11±3 | 1874±490 |
| | Rt | 271±70 | 87±13 | 3211±649 | 979±338 | ND | 183±159 | 3±2 | 5±3 | 4739±396 |
